# Cyanine dye conjugates of a 2’-deoxycytidine-based auto- and mitophagy activator extend *Caenorhabditis elegan*s lifespan

**DOI:** 10.64898/2026.01.23.701232

**Authors:** Ekaterina A. Guseva, Polina N. Kamzeeva, Sofya Y. Sokolskaya, Boris P. Myasnikov, Julia A. Golubeva, Vera A. Alferova, Rose-Lys Zaranaina, Valeriya B. Vays, Irina M. Vangeli, Eugene S. Belyaev, Olga A. Potapova, Natalia N. Gotmanova, Anna V. Bacheva, Lora E. Bakeeva, Elena I. Marusich, Maria P. Rubtsova, Olga A. Dontsova, Peter V. Sergiev, Andrey V. Aralov

## Abstract

**Background:** Autophagy and mitophagy are essential for cellular homeostasis and play key roles in longevity and healthy aging, whereas their age-associated decline contributes to the development of age-related diseases. The identification of small-molecule activators of these pathways therefore represents an important therapeutic objective.

**Methods:** In this study, we investigated a series of compounds based on a 2′-deoxycitidine-derived scaffold and systematically analyzed the impact of structural substitutions on their ability to induce autophagy and mitophagy. Chemical optimization and functional assays were combined with pathway analysis, cellular readouts of proteostasis, and in vivo lifespan assessment in *Caenorhabditis elegans*.

**Results:** The lead compound enhanced autophagy predominantly via activation of the AMPK–ULK1 signaling pathway and induced mitophagy in a Parkin-independent manner. It promoted autophagosome formation and facilitated functional clearance of aggregation-prone mutant huntingtin. Conjugation of the lead compound with the mitochondria-targeting Cy5 dye further potentiated mitophagy induction, likely through preferential mitochondrial accumulation, while reducing cytotoxicity. Importantly, the conjugated compound significantly extended *C. elegans* lifespan at lower concentrations compared with the unconjugated analogue.

**Conclusions:** Together, these results identify a promising chemical scaffold for the development of auto-and mitophagy activators and validate mitochondria-targeted conjugation as an effective strategy to enhance their biological performance. The demonstrated in vivo efficacy supports the potential relevance of these compounds for interventions aimed at preserving proteostasis and mitochondrial quality control, with possible implications for geroprotective applications.

## Introduction

Autophagy is a pivotal intracellular degradation process necessary for the elimination of damaged organelles [1]. Disruption of this process with age has been associated with the development of storage diseases, neurodegenerative diseases (including Parkinson’s and Huntington’s), and cardiovascular diseases [2]. These pathologies are particularly impacted by impaired mitochondrial degradation, which occurs through a specific subtype of autophagy called mitophagy [3].

Autophagy and mitophagy are tightly regulated cellular processes in which AMPK serves as a key activator. During autophagy, AMPK is activated by energy starvation and initiates the ULK1-dependent pathway [4]. In the context of mitophagy, AMPK is activated in parallel with the PINK/Parkin-mediated system of mitochondrial phospho-ubiquitination, and this activation is necessary for the targeting of mitochondria for autophagosomal engulfment [5].

Autophagy levels are known to decline naturally with age [6], and the use of autophagy activators, both natural and synthetic, is considered a promising approach for the development of geroprotective compounds [7–9]. For instance, natural products such as urolithin A, a derivative of ellagic acid [10,11], resveratrol [12], and spermidine [12] have demonstrated efficacy in prolonging lifespan and improving health [13] in ageing rodent [14–16] and worm models [17,18], as well as in human trials [19–21]. As for synthetic inducers, an increasing number of active molecules are being discovered among nucleoside analogs. Indeed, following the discovery of the pharmacological modulator of autophagy, 5-aminoimidazole-4-carboxamide ribonucleoside (AICAr) [22], several activators were found among (2’-deoxy)adenosine [23–26], guanosine [27], and, recently, (2’-deoxy)cytidine analogs [28].

In the present study, we investigated the ability of a set of (2’-deoxy)cytidine and cytosine derivatives with extended heteroaromatic systems, namely benzo[4,5]- and naphtho[2,1:4,5]imidazo[1,2-c]pyrimidinonyl moieties, to induce autophagy/mitophagy using a fluorescent cell reporter assay, immunoblotting, and transmission electron microscopy (TEM) analyses. For the lead compound, the specificity of the influence on the AMPK-dependent pathway and functionally productive autophagic degradation were also demonstrated. To enhance mitophagy-inducing properties, two approaches based on conjugation of the identified leader via the 7-*O*-attached linker group with mitochondria-targeting molecules were explored. After evaluating the biological properties of the synthesized conjugates, the lead compound from the primary screening and the prospective conjugates were assessed for their ability to influence nematodes’ lifespan.

## Methods

### Synthesis of compounds

For details of the synthesis of the compounds, see the supplementary file (purity exceeded 95% according to HPLC analysis).

### Generation and maintenance of mammalian cell lines

The mouse neuroblastoma (N2a) cells was kindly provided by Dr. A.A. Kudriaeva (Shemyakin-Ovchinnikov Institute of Bioorganic Chemistry, Russian Academy of Sciences, Moscow, Russia). The human neuroblastoma (SH-SY5Y), human embryonic kidney (HEK293) and mouse neuroblastoma (N2a) cell lines were cultured in Dulbecco’s modified Eagle medium (DMEM; ServiceBio, China) and were maintained at 37 °C in a humidified atmosphere containing 5% CO₂. Media was supplemented with 10% (v/v) fetal bovine serum (FBS), 2 mM HiGlutaXL (HiMedia, PA, USA), and 1× penicillin–streptomycin (HiMedia, PA, USA).

To assess the ability of the synthesized compounds to induce autophagy, we used an SH-SY5Y cell line stably expressing the GFP-LC3-RFP fluorescent reporter system [26,28,29]. For the evaluation of mitophagy induction, we employed a monoclonal HEK293 cell line expressing the TOM20MTS-mCherry-EGFP-Tet-On fluorescent reporter system [30].

The reporter-expressing cell lines were generated by lentiviral transduction. Lentivirus was produced by transfecting HEK293T cells with either pMXs GFP-LC3-RFP (Addgene #117413, [29]) or TOM20MTS-mCherry-EGFP-Tet-On (Addgene #109016, [30]), together with the packaging plasmids pCMV-VSV-G (Addgene #8454) and psPAX2 (Addgene #12260), using Lipofectamine 3000 (Thermo Fisher Scientific, USA). Wild-type SH-SY5Y and HEK293T cells were then infected with the lentivirus, and stable transformants were isolated by fluorescence-activated cell sorting (BD FACSAria III), resulting in monoclonal cell lines.

To investigate the mechanism of action of **1e**, we used an SH-SY5Y cell line with an inactivated *AMPK* gene, as previously described [26,28].

To obtain an SH-SY5Y cell line with an inactivated *PRKN* gene, guide RNAs for Cas9 (gCgACgACCCCAgAAACgCgg) targeting the third coding exon of *PRKN* were cloned into the PX458 plasmid (Addgene #48138). Then, SH-SY5Y cells were transfected with the obtained pX458 plasmids containing the gRNA sequences using a Lipofectamine 3000 reagent (Thermo Fisher Scientific, MA, USA). The next day, GFP-positive cells were sorted using BD FACS Aria III (BD Biosciences, CA, USA). After this, the cells were seeded into monoclonal cell lines, which were then analyzed by sequencing and Western blotting. Individual clones were analyzed by sequencing of PCR amplification products of the *PRKN* (5D-TCAGCATTCTATTGTGTTTCACGT-3D and 5D-CCTACAGTGATGTCTCCTTGTAGT-3D) and Western blotting.

To obtain N2a expressing huntingtin gene variants, cells were co-transfected with 1 μg of total plasmid DNA (950 ng pSBtet-Neo-HttQ15/HttQ138 + 50 ng SB100x transposase coding vector) in Opti-MEM media (Gibco, USA) using a Lipofectamine 3000 transfection kit (Invitrogen, USA) according to the manufacturer’s instructions. After co-transfection, 1 mg/ml G418 (Sigma-Aldrich, USA) was added to culture medium for 10-12 days until 100% cell death was achieved in the negative control. A portion of the resulting polyclonal culture was then transferred to a 96-well culture plate, seeding 1, 5, and 10 cells/ml into each 32 wells and then incubated on 1 mg/ml G418 for 10-14 days. As a result, 4 candidate Neuro-2a monoclones with each huntingtin gene variant were selected. Htt/mHtt expression levels in Neuro-2a transgenic cells were verified by qPCR after induction by 1 μg/ml doxycycline (Sigma-Aldrich, USA).

### Screening of autophagy inducers

In the GFP-LC3-RFP reporter system, RFP is diffusely distributed throughout the cell cytoplasm, and autophagy does not affect the level of its fluorescence. GFP, together with LC3, is incorporated into autophagosomes upon the activation of autophagy. When the autophagosome and the lysosome fuse, the pH inside the vesicle is reduced, which leads to a decrease in the cellular GFP fluorescent signal. Thus, by comparing the ratio of red and green fluorescent signals in cells under different conditions, we can draw a conclusion about the level of autophagy activity.

For the screening of autophagy activity, cells of the reporter cell lines were seeded into 96-well plates (655866, Greiner Bio-one, Germany) in DMEM medium (ServiceBio, China) at a concentration of 2*10^4^ cells per well. After 24 h incubation, cells were treated with the compounds under study, control compound or blank solution (FCCP 5 μM and DMSO 0.1%, respectively). After 24 h incubation, 4 microphotographs (10х lens) of each well were taken using a CELENA® X High Content Imaging System (Logos Biosystems) (provided by the Moscow State University Development Program). The CELENA X Cell Analyser software was used to calculate the average intensity of RFP and GFP signals in each cell individually. At least 500 cells were analyzed for each experiment. The ANOVA-test was used to compare the RFP/GFP ratio of cell populations under different conditions.

### Fluorescent microscopy

To evaluate the ability of **1e** and **1f** to induce mitophagy, a HEK293T cell line expressing the TOM20MTS-mCherry-EGFP-Tet-On reporter system was used. Cells were treated with doxycycline (2 ug/mL, Terno Fisher Scientific, USA) 24 h before experiment. The seeding and treatment procedures were the same as for the screening procedure. Microphotographs were acquired at 60× magnification using a CELENA® X High Content Imaging System (Logos Biosystems, South Korea). Image analysis was performed using Fiji 20250514.1117 [31].

Human neuroblastoma cells were seeded into 96-well plates (655866, Greiner Bio-one, Germany) at a density of 2 × 10^4^ cells per well. After 24 h incubation with **8a-d**, SH-SY5Y cells were washed twice with PBS and stained with MitoTracker Green (200 nM, 15 min), LysoTracker Red (150 nM, 15 min) or LysoTracker Green (150 nM, 15 min). Images were acquired using a Nikon C2 fluorescence confocal microscope system (Tokyo, Japan).

### Immunoblotting

Immunoblotting of cell lysates was performed as previously described [32]. Briefly, 24h before the experiment, production of HTT or mut-HTT was induced with doxycycline (2 ug/mL, Terno Fisher Scientific, USA) in N2a cells. Further, N2a or SH-SY5Y cells were incubated with the compounds for 24 h and lysed in a RIPA buffer (150 mM sodium chloride, 50 mM Tris-HCl, pH 8.0, 0.5% Nonidet P-40, 1% sodium deoxycholate, 0.5% SDS) supplemented with a protease inhibitor cocktail (Thermo Fisher Scientific, USA). Lysates were separated on 12–15% SDS-PAGE gels and transferred onto PVDF membranes (0.45 μm, Thermo Fisher Scientific, USA) using wet transfer. Membranes were blocked for 1 h at room temperature in TBST (10 mM Tris-HCl, pH 7.5, 150 mM NaCl, 0.1% Tween-20) containing 5% bovine serum albumin (BSA, Proliant Biologicals, USA).

Primary antibodies were diluted 1:1000 in TBST with 5% BSA. The following antibodies were used: Tuberin/TSC2 (3612, Cell Signaling Tech., USA), phospho-Tuberin/TSC2 (Ser1387) (5584, Cell Signaling Tech., USA), ULK1 (8054, Cell Signaling Tech., USA), phospho-ULK1 (Ser555) (5869, Cell Signaling Tech., USA), phospho-ULK1 (Ser317) (89267, Cell Signaling Tech., USA), phospho-ULK1 (Ser757) (6888, Cell Signaling Tech., USA), AKT (9272, Cell Signaling Tech., USA), phospho-Akt (Thr308) (9275, Cell Signaling Tech., USA), AMPK (2532, Cell Signaling Tech., USA), phospho-AMPK (T172) (2535, Cell Signaling Tech., USA), 4E-BP1 (9644, Cell Signaling Tech., USA), phospho-4E-BP1 (Thr37/46) (236B4, Cell Signaling Tech., USA), LC3 (4108, Cell Signaling Tech., USA), Becline (Cell Signaling Tech., USA), phospho-Ubiquitin (Ser65) (E2J6T, Cell Signaling Tech., USA), Parkin (#2132, Cell Signaling Tech., USA), anti-Flag (1:3000, F1804, Sigma-Aldrich, USA). α-tubulin (1:3000, ab18251, Abcam, USA), GAPDH (1:3000, ab8245, Abcam, USA) and β-actine (ab8229, Abcam, USA) were used as a loading control.

Secondary HRP-conjugated anti-rabbit (1706515, Biorad, USA) and anti-mouse (7076, Cell signaling, USA) antibodies were used at 1:5000 dilution.

### Cytotoxicity assays

Human neuroblastoma SH-SY5Y cells were seeded at a density of 20,000 cells per 100 μl DMEM/F12 medium per well in 96-well plates. The maximum concentration used for **1e** and **8a-d** was 1000 μM, and the final DMSO concentration did not exceed 0.5% (v/v).

Cell viability was assessed using the MTT assay (**1e**) or resazurin assay (**8a-d**) after 24 h of compound treatment. Cells were incubated with MTT reagent (0.5 g/l; Paneco LLC, Russia) or resazurin (0.15 mg/ml, Thermo Fisher Scientific, USA) for 2 h at 37 °C in a CO₂ incubator. After incubation with the MTT reagent, the medium was removed, and 140 μl of DMSO were added to each well. Plates were incubated for at least 15 min at room temperature on an orbital shaker to solubilize formazan crystals. Absorbance was measured at 545 nm using a VICTOR X5 plate reader (PerkinElmer, USA). After incubation with resazurin, fluorescence was recorded using a 560 nm excitation / 590 nm emission filter set using a VICTOR X5 plate reader (PerkinElmer, USA).

Cytotoxicity data were analyzed using nonlinear regression in GraphPad Prism 9, applying a four-parameter logistic model (lower limit, upper limit, slope, and CC₅₀).

### DNA extraction and RT PCR

Assessment of the mitochondrial-to-nuclear DNA ratio was done as previously described [33]. DNA was extracted from 1 × 10^6^ cells with LumiSpin® UNI DNA Isolation Spin Kit for Any Sample (Lumiprobe, Russia). Quantitative PCR gene amplifications were performed using a SYBR® Green PCR master mix (Ther moFisher Scientific, Waltham, MA, USA) in the CFX384 Touch Real-Time PCR System. qPCR was conducted using primer sets: preGAPDH 5’-CCACCAACTGCTTAGCACC-3’, 5’-CTCCCCACCTTGAAAGGAAAT-3’, and 12S 5’-AAACCCCgATCAACCTCACC-3’, 5’-TTTACgTgggTACTTgCgCT-3’. The amount of mitochondrial DNA was calculated by the 2-ΔΔCT method and normalized to nuclear DNA.

### Lifespan assay on *C. elegans* model

To assess the effects of compounds **1e**, **8c**, and **8d** on nematode lifespan, the wild-type *Caenorhabditis elegan*s N2 Bristol strain (The Swammerdam Institute of Life Sciences, University of Amsterdam, Netherlands) was used. The Escherichia coli OP50 strain (Engelhard Institute of Molecular Biology, Moscow, Russia) was used as a food source.

Lifespan assays were performed using synchronized L4-stage adult nematodes. Synchronization was carried out as described previously [34]. Briefly, 120 µl of synchronized L1 larvae were dispensed into 96-well plates (TPP, Switzerland). After 48 h, 15 µl of 5-fluoro-2′-deoxyuridine (FUDR; Merck, Germany) were added to each well to inhibit progeny production. After a further 24 h, 15 µl of **1e**, **8c**, or **8d** solution were added to obtain the final concentrations of 500, 250, 100, 50, 10, and 1 µg/ml.

Each concentration was tested in six replicates. Lifespan extension efficacy was evaluated relative to untreated controls. Survival data were analyzed using GraphPad Prism (GraphPad Software, LLC, USA) with Kaplan–Meier survival analysis.

## Results

### Primary screening of autophagy activation among benzo[4,5]- and naphtho[2,1:4,5]imidazo[1,2-c]pyrimidinone-based compounds

Primary screening was carried out among the previously described compounds containing a benzo[4,5]- or naphtho[2,1:4,5]imidazo[1,2-c]pyrimidinone scaffold (Fig. 1) [35,36]. In particular, the first subset included 7-*O*-alkylated/non-alkylated benzo[4,5]imidazo[1,2-c]pyrimidinones substituted at the 2-position with 2’-deoxyribose (**1**), ribose (**2**), or left unsubstituted (**3**). In some cases, the sugar moiety was substituted with a *4*,*4’*-dimethoxytrityl (DMTr) moiety at the 5’-hydroxyl group (as in **1b** and **2b**). The compounds were used in phenotypic screening for antiviral activity against a panel of structurally and phylogenetically diverse RNA and DNA viruses, but most, with the exception of **1b** and **2b**, did not exhibit either virus-inhibitory activity or cytotoxicity towards host cells [36]. The second subset consisted of benzo[4,5]- and naphtho[2,1:4,5]imidazo[1,2-c]pyrimidinones **4** and **5**, respectively, bearing structurally diverse substituents at the 2- and 7-(for **4**) or 5-(for **5**) positions. They were probed as putative G4 ligands using a set of biophysical methods and, since G4 ligands are considered as potential antitumor agents [37], their cytotoxicity for tumor and non-tumor cell lines was also evaluated. With the exception of **5**с, all compounds had no significant effect on cellular metabolic activity.

**Fig. 1.**
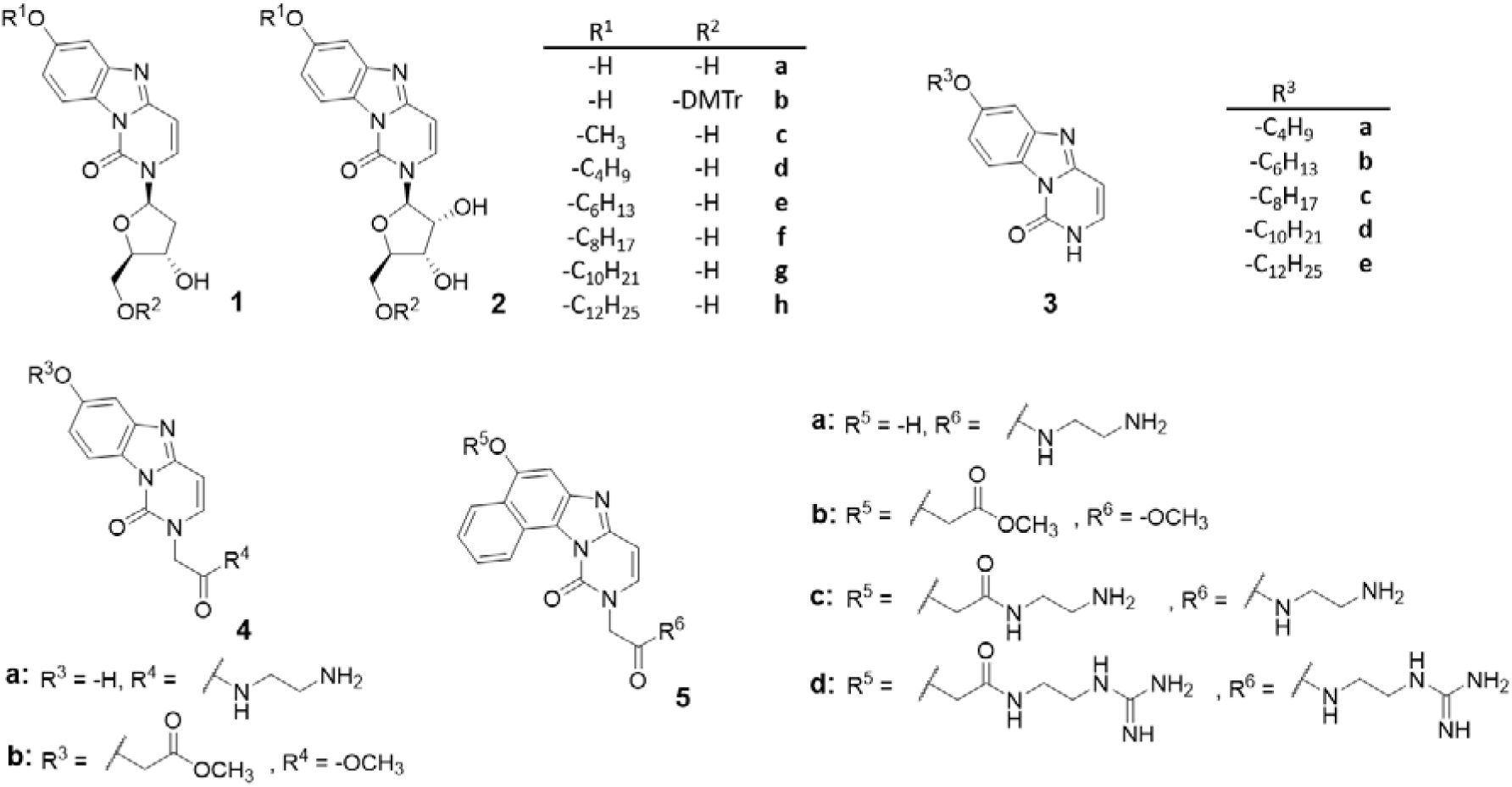
Chemical structures of compounds used in primary screening. Reported procedures were applied for the preparation of **1**-**3** [36] and **4**,**5** [35].

To study the autophagy-activating properties of the compounds, fluorescence screening was performed using a previously obtained human neuroblastoma cell line that constitutively expresses the pMXs GFP-LC3-RFP reporter system [26]. Upon autophagy activation, LC3-GFP fluorescence decreases in response to the acidic environment of autolysosomes, while the background constitutive RFP fluorescence remains unchanged and is used as an internal control. This reporter system enables highly sensitive detection of autophagic flux.

Each compound was tested at three concentrations: 250 μM, 25 μM, and 2.5 μM and autophagy activation was measured after 24 h incubation. Screening of the 27 compounds revealed that 16 of them exhibited autophagy-activating properties at one or more of the tested concentrations (Fig. 2A, Supp. Fig. 1A). Furthermore, two compounds (**4b** and **5d**) exhibited autophagy inhibition at the lowest concentration tested.

**Fig. 2.**
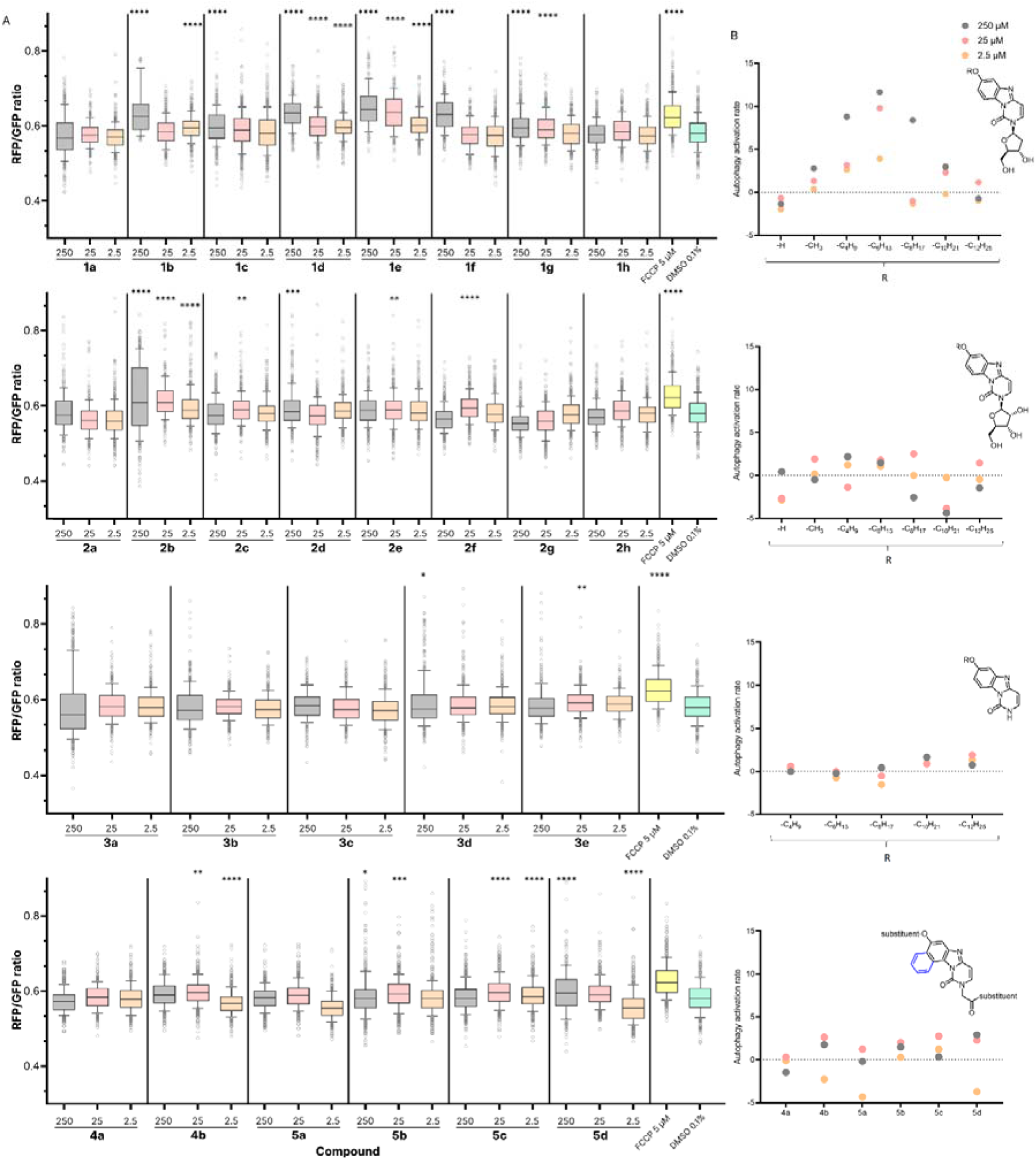
Results of primary screening of autophagy inducers among the reported compounds. A. Changes in the RFP/GFP ratio in the SH-SY5Y pMXs GFP-LC3-RFP cell line when treated with compounds under study. The results of ANOVA tests are indicated on the graph: * – p-value <0.05, ** – p-value <0.01, **** – p-value <0.0001. B. Dependence of the autophagy activation rate on selected substituents in four subgroups of compounds under study.

The greatest number of activators was found among benzo[4,5]imidazo[1,2-c]pyrimidinonyl derivatives **1** of the 2’-deoxy series, and the level of autophagy activation was observed to depend on the length of the aliphatic substituent at the 7-hydroxyl group of the heterocyclic system (Fig. 2B, upper panel). A bell-shaped dependence of activity on the alkyl length was revealed, with compound **1e** substituted with *n*-hexyl having the highest activity.

The results were confirmed by immunoblotting on the key autophagy marker protein, LC3. The accumulation of processed form of this protein with higher electrophoretic mobility indicates autophagy activation in cells (Fig. 3A,B, Supp. Fig. 1B,C). Since immunoblotting is a less sensitive method of detecting autophagy activation, it can only identify the most active inducers [38]. According to the results obtained, the most active autophagy inducers were found among benzo[4,5]imidazo[1,2-c]pyrimidinonyl derivatives **1** (particularly **1e**) of the 2’-deoxy series, which is consistent with the results of fluorescence screening.

**Fig. 3.**
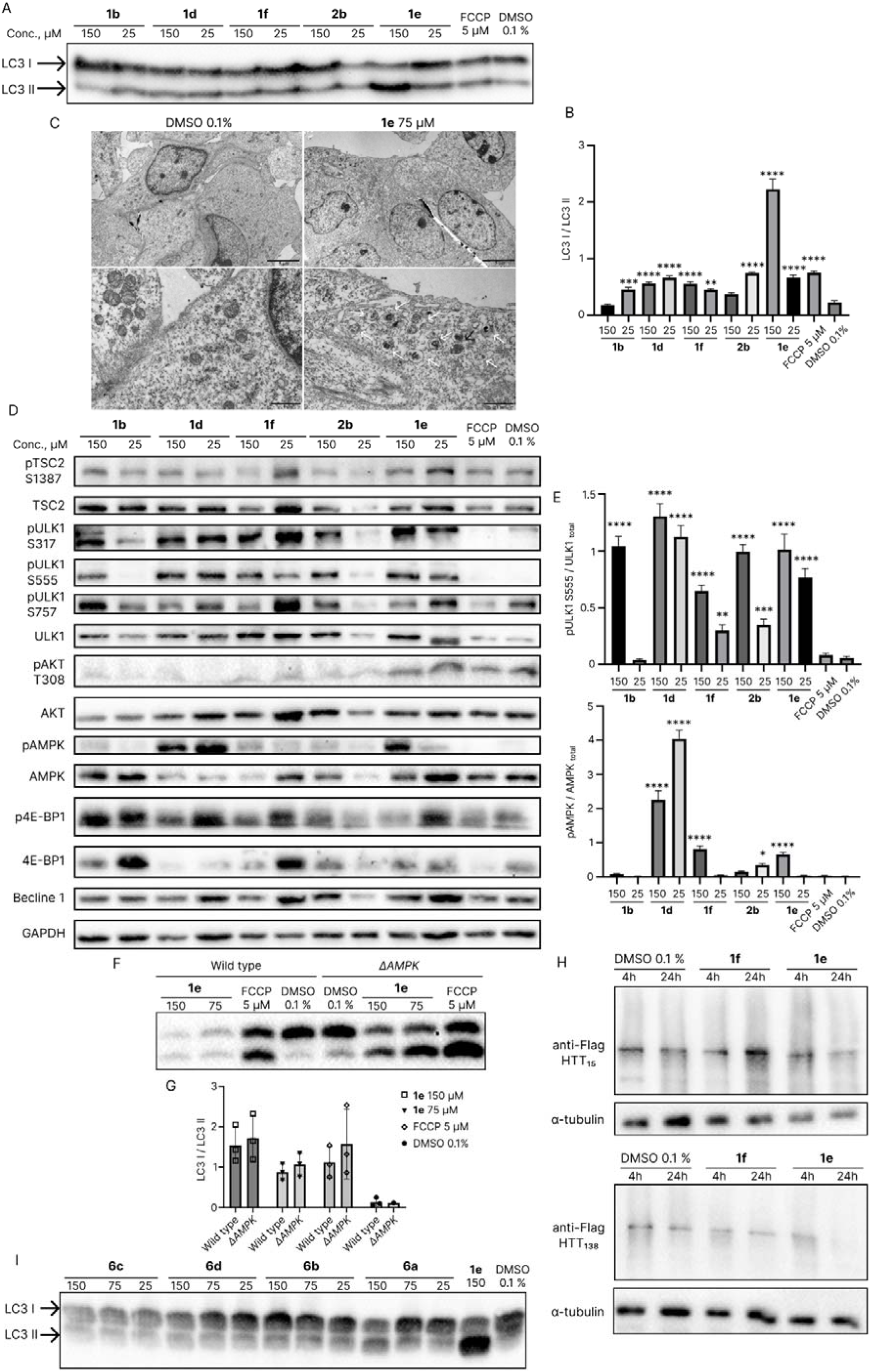
Investigation of the mechanism of autophagy induction by benzo[4,5]imidazo[1,2-c]pyrimidinonyl derivatives. A. Western blotting on the key marker of autophagy (LC3) on SH-SY5Y when treated with derivatives **1b**, **1d-f**, and **2b**. B. Change in the ratio of LC3 forms when treated with **1e**. Data represent mean ± SD (n=3). The results of ANOVA tests are indicated on the graph: ** – p-value <0.01, *** – p-value <0.001, **** – p-value <0.0001. C. Electron microphotographs showing accumulation of autophagosomes after **1e** treatment. Black bars indicate 5 μm (upper panel) and 1 μm (bottom panel). White arrows indicate autophagosomes with folded-membrane structures; black arrows indicate autophagosomes with electron-dense material. D. Western blotting for various protein regulators of the autophagy cascade. E. Changes in phosphorylation status and total amount of main autophagy regulators. Data represent mean ± SD (n = 3). The results of ANOVA tests are indicated on the graph: * – p-value <0.05, ** – p-value <0.01, *** – p-value <0.001, **** – p-value <0.0001. F. Western blotting on the key marker of autophagy (LC3) on SH-SY5Y with inactivated *AMPK* gene and wild type after incubation with **1e**. G. Change in the ratio of LC3 forms when treated with **1e**. Data represent mean ± SD (n=3). H. Western blotting for degradation of normal and mutant form of huntingtin in N2a cell line under treatment with **1f** and **1e**. Induction of both forms of huntingtin was performed 24 h prior the experiment. I. Western blotting on the key marker of autophagy (LC3) on SH-SY5Y when treated with derivatives **6a**-**d** from an additional set.

To further prove autophagy-activating properties of **1e**, we performed TEM analysis (Fig. 3C). Ultrastructural analysis revealed the accumulation of autolysosomal structures surrounded by double membranes, ranging from approximately 200 nm to 500 nm in diameter (Fig. 3C). These vesicles contained predominantly folded membranous structures indicative of organelle degradation, as well as electron-dense material, likely corresponding to lipid- and protein-rich cargo. These observations further illustrated the autophagy-activating properties of lead compound **1e**.

### Benzo[4,5]imidazo[1,2-c]pyrimidinonyl derivatives 1 of the 2’-deoxy series activate autophagy predominantly via the AMPK–ULK1-pathway

Upon closer examination of the selected lead compound **1e**, as well as its closest analogs **1b**, **1d**, **1f**, and **2b**, we found that the treatment of cells with these compounds led to a pronounced increase in AMPK activation (Fig. 3D,E). Enhanced AMPK activity was further supported by increased AMPK-dependent phosphorylation of ULK1 at residues S555 and S317, which are key regulatory sites involved in autophagy initiation. The simultaneous activation of AMPK and phosphorylation of its downstream effector ULK1 indicates that autophagy induction by this compound series proceeds predominantly through the canonical AMPK/ULK1 signaling axis.

In contrast, the AKT/TSC2/mTOR-dependent inhibitory pathway of autophagy regulation remained largely intact. This was evidenced by preserved phosphorylation levels of TSC2 and the mTOR downstream target 4E-BP1. Although activating phosphorylation of AKT was significantly reduced upon incubation with most compounds (**1b, 1d, 1f** and **2d**), this decrease was not accompanied by suppression of downstream mTOR signaling. These observations indicate that the compounds do not induce autophagy through global inhibition of anabolic signaling pathways, but rather selectively engage AMPK-dependent autophagy activation.

Since compound **1e** demonstrated the highest activity across all assays, it was selected for further mechanistic studies. To assess the contribution of AMPK to autophagy induction, we examined the effect of **1e** in a neuroblastoma cell line with inactivation of *AMPK* (Δ*AMPK*). Analysis of LC3 forms ratio revealed that compound **1e** preserved autophagy-activating properties in Δ*AMPK* cells (Fig. 3F,G), indicating that autophagy induction by **1e** leads to AMPK activation in cells.

To evaluate whether the developed autophagy activator might be useful in clinical applications, we next examined the processive clearance of protein aggregates in a neuroblastoma cell model with inducible overexpression of normal (Gln15) and mutant (Gln138) forms of huntingtin [39]. Huntingtin expression was induced 24 h prior to treatment, and protein levels were assessed by immunoblotting at 4- and 24-hour time points following incubation with the test compounds. As a result, compound **1e** was found to preferentially promote degradation of the mutant form of huntingtin (Fig. 3H), which is consistent with enhanced autophagic flux and selective clearance of aggregated substrates, rather than nonspecific protein degradation.

Thus, activation of the AMPK/ULK1 signaling pathway by compound **1e** is accompanied by enhanced functional clearance of aggregation-prone mutant huntingtin, supporting the conclusion that the observed signaling changes result in productive autophagic degradation.

### Variation of the aliphatic substituent at the 7-position of the lead compound does not further enhance the autophagy-activating properties

Since benzo[4,5]imidazo[1,2-c]pyrimidinonyl derivatives **1d** and **1e** containing an *n*-butyl (C_4_) or *n*-hexyl (C_6_) group, respectively, were the most potent autophagy activators among the compounds used in the primary screening (Fig. 1A, Supp. Fig. 1A), an additional set of 2’-deoxyribosyl derivatives bearing an *n*-pentyl (**6a**), *iso*-pentyl (**6b**), phenethyl (**6c**), and phenpropyl (**6d**) moiety was synthesized (Scheme 1). Compound **1b** [36] was subjected to Mitsunobu coupling with *n*-propanol, *iso*-propanol, 2-phenylethan-1-ol, or 3-phenylpropan-1-ol, followed by DMTr removal in mild acidic conditions (aqueous CH_3_COOH, 50 °C) to afford the target derivatives **6a–d**, respectively. However, after studying their autophagy-activating properties, we found that **1e** outperformed the additional members of the series (Fig. 3I).

**Scheme 1.**
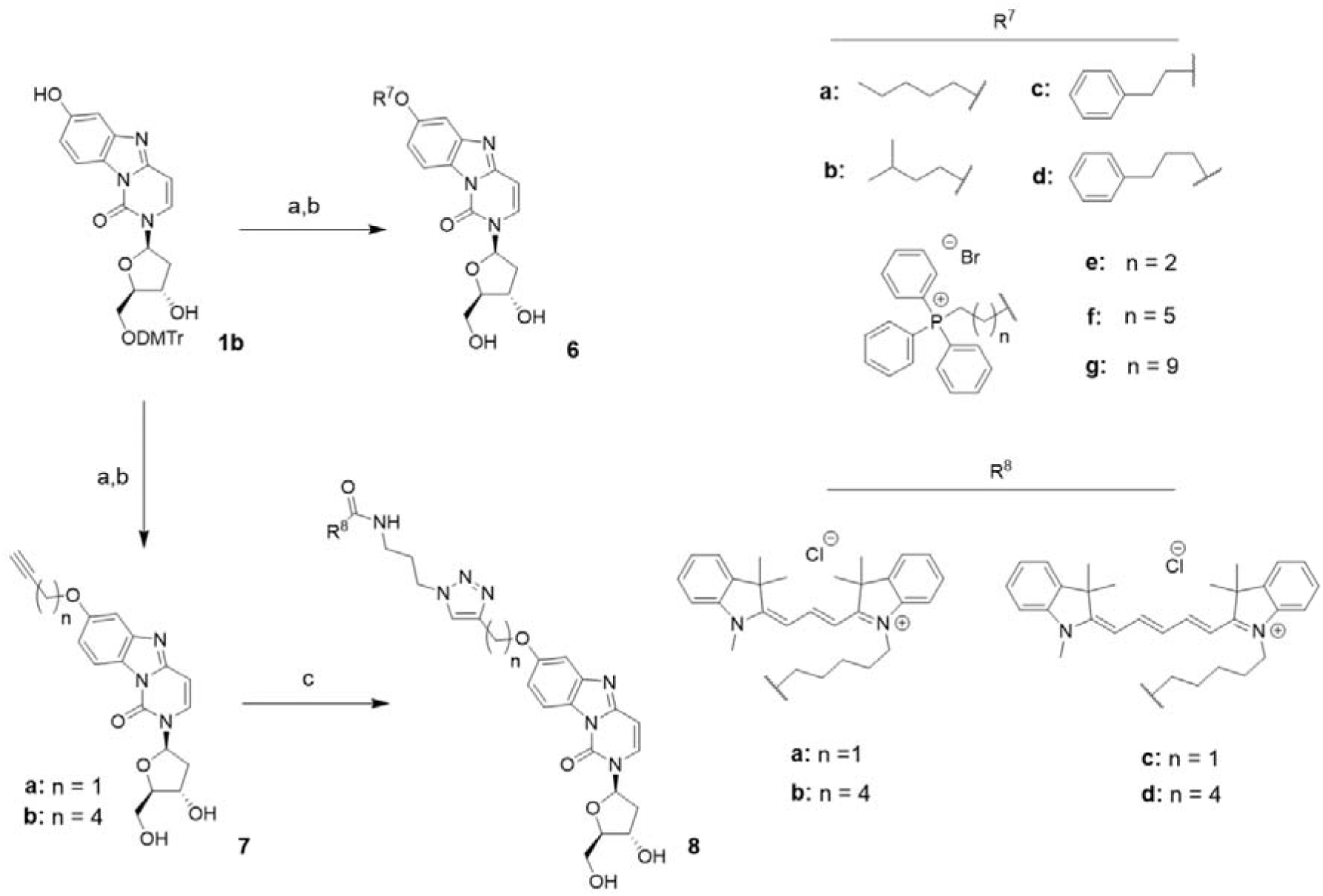
The preparation of additional benzo[4,5]imidazo[1,2-c]pyrimidinonyl derivatives of the 2’-deoxy series. Reagents and conditions: (a) *n*-propanol (for **6a**), *iso*-propanol (for **6b**), 2-phenylethan-1-ol (for **6c**), 3-phenylpropan-1-ol (for **6d**), (3-hydroxypropyl)triphenylphosphonium bromide (for **6e**), (6-hydroxyhexyl)triphenylphosphonium bromide (for **6f**), (10-hydroxydecyl)triphenylphosphonium bromide (for **6g**), prop-2-yn-1-ol (for **7a**) or hex-5-yn-1-ol (for **7b**), PPh_3_, DIAD, CH_2_Cl_2_, 0 °C→rt; (b) CH_3_COOH, H_2_O, 50°C; (c) cyanine3 azide (**8a,b** or cyanine5 azide (**8c,d**), CuSO_4_, TBTA, NaAsc, DMSO, H_2_O, rt.

### The lead compound 1e effectively activates mitophagy in a Parkin-independent manner

Besides general autophagy, cells also undergo more specific processes of selective degradation of damaged organelles. Of particular interest is mitochondrial degradation, or mitophagy. Analysis of mitochondrial turnover revealed that compound **1e** induces mitophagy that does not require functional Parkin.

Upon analyzing the accumulation of phosphorylated ubiquitin in cells, we found that the derivatives are capable of activating mitophagy in addition to general autophagy. This effect was observed for compounds **1e** and **1f**, and to a lesser extent for **1d**. Furthermore, compound **1e**, selected as the most potent autophagy inducer, was also the most potent mitophagy activator (Fig. 4A,B). To assess whether mitochondria are delivered to lysosomes, we used HEK293T cells expressing TOM20MTS-mCherry-EGFP under a doxycycline-inducible promoter. Even at concentrations of 100 μM and 50 μM, compound **1e** significantly promoted mitochondrial entry into lysosomes (Fig. 4C,D, Supp. Fig. 2), whereas compounds **1f** and **1d** exhibited mitophagy-activating properties to a lesser extent.

**Fig. 4.**
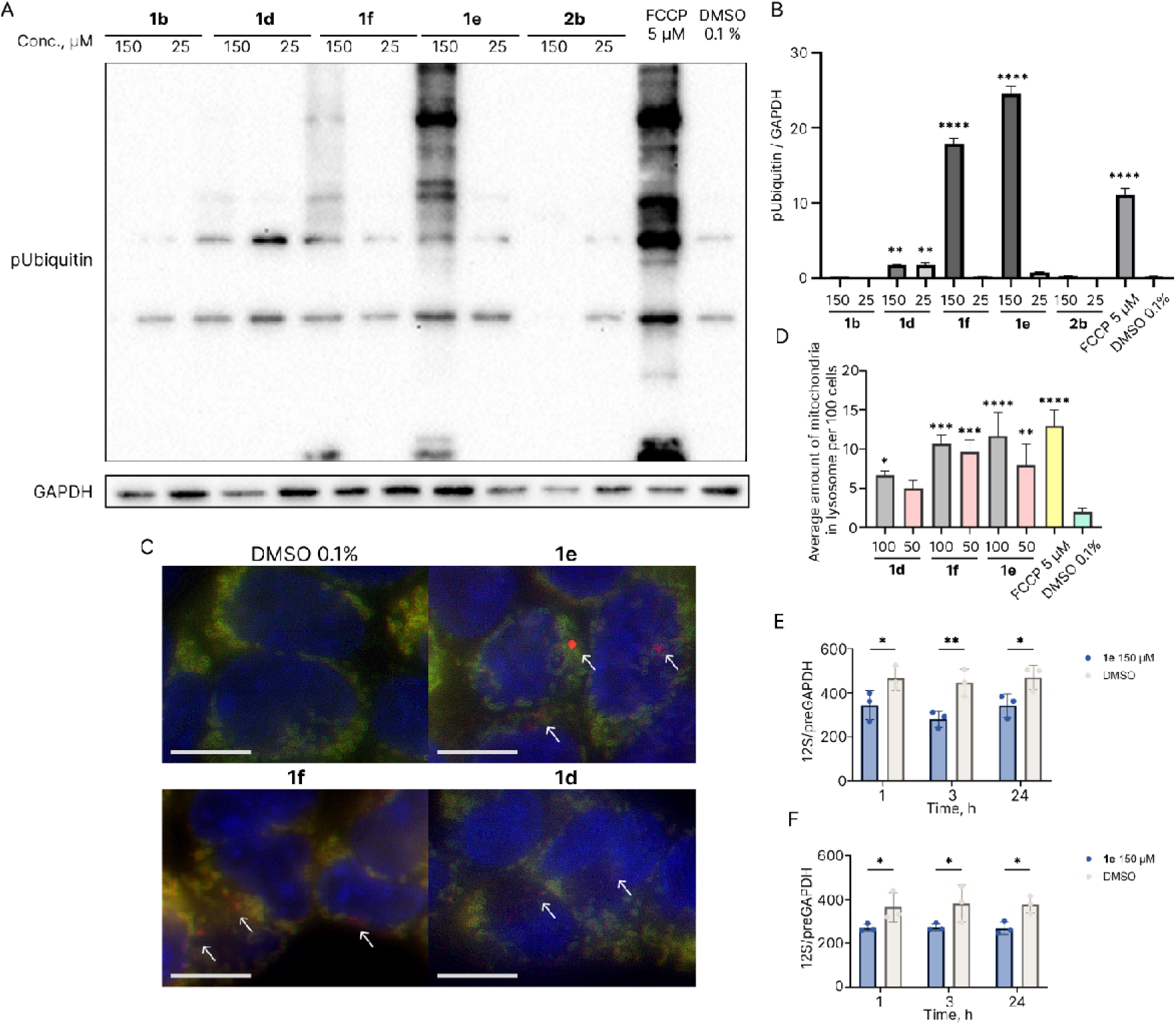
Investigation of the mitophagy activating properties of benzo[4,5]imidazo[1,2-c]pyrimidinonyl derivatives. A. Western blotting for the key marker of mitophagy (p-Ubiquitin) B. Bar graph representing changes in level of p-Ubiquitin normalized on GAPDH in SH-SY5Y cells under treatment with **1b**, **1d**, **1e**, **1f**, **2b.** Data represent mean ± SD (n = 3). The results of ANOVA tests are indicated on the graph: * – p-value <0.05, ** – p-value <0.01, **** – p-value <0.0001. C. Mitophagy induced by 100 μM **1d-f** in wild-type HEK293T expressing TOM20MTS-mCherry-EGFP-Tet-On. White bars indicate 10 μm. White arrows mark mitochondria in lysosomes. Nuclei were stained with Hoechst (blue). Cells were treated with doxycycline (2 ug/mL) 24 h before the experiment. D. Bar graph indicating average amount of mitochondri in lysosome in HEK293T cells expressing TOM20MTS-mCherry-EGFP-Tet-On (n=3). The results of t-test are indicated on the graph: * – p-value < 0.05, ** – p-value < 0.01, *** – p-value < 0.001, **** – p-value < 0.0001. E. Number of mitochondria as addressed by quantitative PCR of mtDNA (*12S* gene) vs. nuclear DNA (*GAPDH* gene) from the wild-type SH-SY5Y cells after treatment for 1, 3 or 24 h with **1e.** Data represents mean ± SD (n=3). The results of t-test are indicated on the graph: * – p-value < 0.05, ** – p-value < 0.01. F. Number of mitochondria as addressed by quantitative PCR of mtDNA (*12S* gene) vs. nuclear DNA (*GAPDH* gene) from the Δ*PRKN* SH-SY5Y cells after treatment for 1, 3, 24 hours with **1e**. Data represents mean ± SD (n=3). The results of t-test are indicated on the graph: * – p-value < 0.05.

The observed mitophagy-activating properties were further supported by ultrastructural analysis using TEM (Fig. 3C). Cells treated with **1e** exhibited an increased number of autolysosomes containing folded membrane structures and fragmented mitochondria. After 24 h of cell incubation with **1e**, predominantly late-stage autolysosomes containing mitochondria can be detected. These can be identified by their double membrane and remnants of the cristae structure (Fig. 4C, marked by white arrows).

Consistent with these findings, the lead compound **1e** not only triggered mitochondrial phospho-ubiquitination and lysosomal delivery, but also resulted in progressive mitochondrial degradation (Fig. 4E), as confirmed by a reduction in mitochondrial DNA content. Although accumulation of phospho-ubiquitin following treatment with **1e** is commonly associated with PINK1/Parkin signaling, mitochondrial degradation was not affected by *PRKN* inactivation (Fig. 4F, Supp. Fig. 3), providing direct evidence that mitophagy induced by **1e** proceeds independently of Parkin.

Together, these data demonstrate that compound **1e** promotes selective mitochondrial turnover characterized by mitochondrial ubiquitination, lysosomal delivery, and loss of mitochondrial mass, and that this process occurs independently of Parkin.

### Conjugation of the lead compound with the Cy5 dye, but not with the Cy3 dye or triphenylphosphonium (TPP^+^) moiety, increases its ability to activate mitophagy

To improve the biological characteristics of the identified lead compound, two approaches based on its conjugation with triphenylphosphonium (TPP^+^) moiety or cyanine dyes through a linker group for specific mitochondrial targeting were employed. The former has long been recognized as an effective means to deliver probes, antioxidants, and pharmacophores to mitochondria [40–42]. The latter uses mitochondria-targeting cyanine dyes consisting of two terminal heterocyclic units linked by a π-conjugated polymethine chain and containing a positively charged nitrogen atom [43,44]. Among them, cyanine3 (Cy3) and cyanine5 (Cy5) are of particular interest, taking into account their comprehensive study and commercial availability [45–47].

To obtain TPP^+^-conjugated derivatives **6** with a linker of varying length (propylene for **6e**, hexylene for **6f**, and decylene for **6g**) connecting the benzo[4,5]imidazo[1,2-c]pyrimidinone 2’-deoxyribosyl moiety and the mitochondria-targeting molecule, reported (hydroxyalkyl)triphenylphosphonium bromides were used in a Mitsunobu coupling reaction (Scheme 1).

Among these compounds, only derivative **6f** exhibited detectable mitophagy-activating properties; however, its activity was substantially lower than that of the compound **1e** (Supp. Fig. 4A). Extending the linker length further revealed that the incorporation of a long decylene spacer (as in **6g**) enhanced general autophagy activation (Supp. Fig. 4A), but was accompanied by increased cytotoxicity (data not shown).

To implement the second approach, conjugates of the lead compound with Cy3 and Cy5 were obtained. For this purpose, derivatives **7a** and **7b** containing a propynyl or hexynyl group, respectively, were synthesized and then used in a copper-catalyzed azide-alkyne cycloaddition (CuAAC) [48] with commercial azido-containing cyanines Cy3 azide and Cy5 azide to afford the target derivatives **8a-d** (Scheme 1).

While the addition of the Cy3 dye significantly impaired the mitophagy-activating properties compared to **1e** (Supp. Fig. 4B), conjugation of a 2-(2′-deoxy-β-D-ribofuranosyl)-benzo[4,5]imidazo[1,2-c]pyrimidinonyl moiety with the Cy5 dye enhanced these properties, especially in the case of **8c** with a shorter linker group (Fig. 5A,B).

**Fig. 5.**
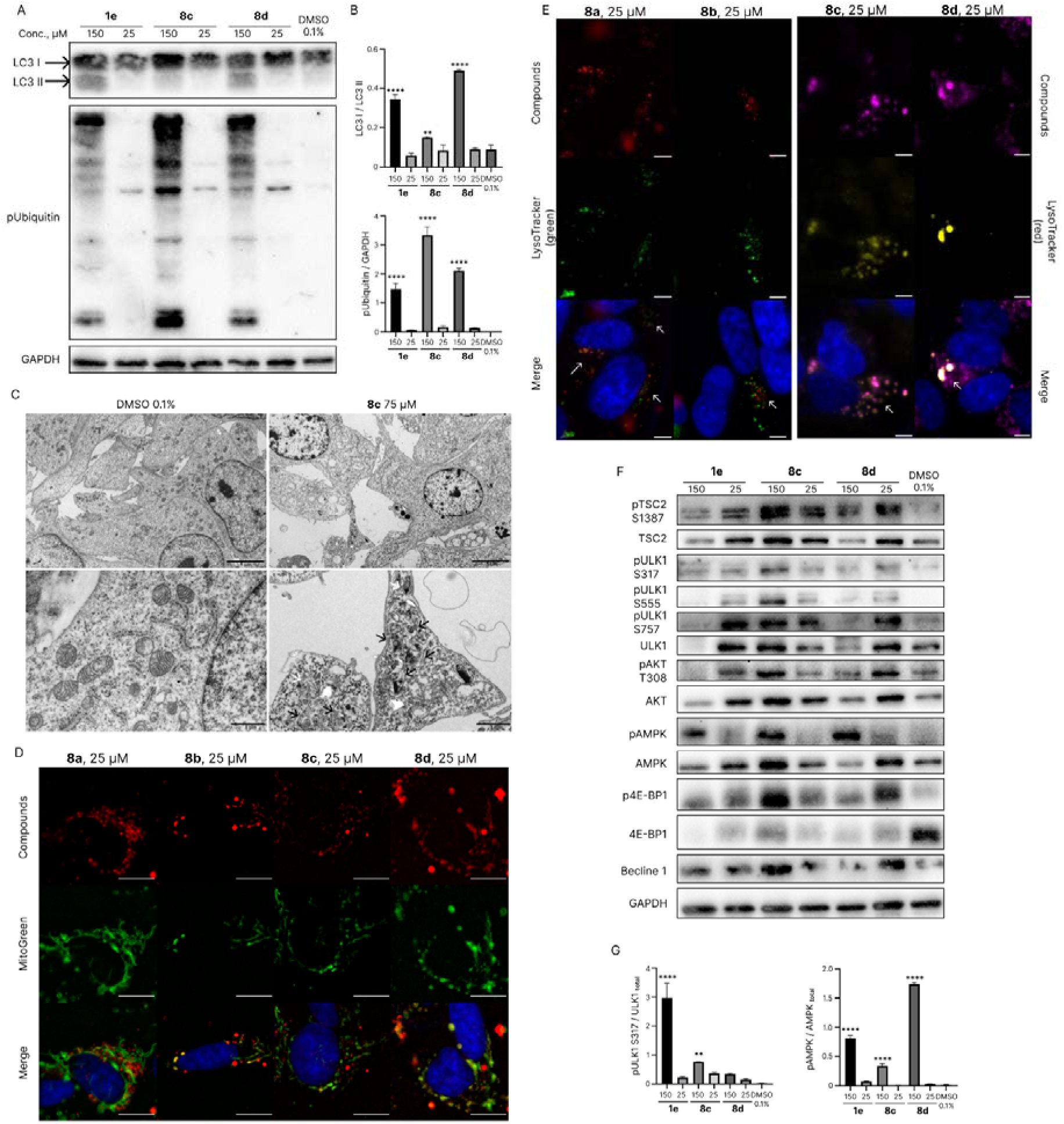
Investigation of the mitophagy activating properties of Cy3/Cy5-bearing conjugates **8a-d.** A. Western blotting for key markers of autophagy (LC3) and mitophagy (p-Ubiquitin). B. Changes in LC3 forms and in level of p-Ubiquitin normalized by GAPDH in SH-SY5Y cells under treatment with **1e**, **8c**, and **8d**. Data represent mean ± SD (n = 3). The results of ANOVA tests are indicated on the graph: * – p-value <0.05, ** – p-value <0.01, **** – p-value <0.0001. C. Electron microphotographs showing accumulation of autophagosomes after **8**с treatment. White arrows indicate fusion of the mito-phagofore with lysosome, black arrows indicate autophagosomes at the final stage of mitophagy. D-E. SH-SY5Y were treated with **8a-d** at a concentration of 25 μM, incubated for 24 h, and then stained with MitoGreen (200 nM, 15 min) (D), or LysoTracker green (150 nM, 15 min) (E, left panel) or LysoTracker red (150 nM, 15 min) (E, right panel). White arrows indicate colocalization of lysosome and added compound. White bars indicate 5 μm. F. Western blotting for various protein regulators of the autophagy cascade. G. Changes in phosphorylation status and total amount of main autophagy regulators. Data represent mean ± SD (n = 3). The results of ANOVA tests are indicated on the graph: ** – p-value <0.01, **** – p-value <0.0001.

The enhanced mitophagy-inducing properties of Cy5 conjugates were further supported by TEM. Neuroblastoma cells treated with compound **8c** displayed a pronounced accumulation of double-membrane autophagic structures containing degrading mitochondria, observed both during fusion of the mitophagosome with lysosomes and at later stages of mitochondrial degradation (Fig. 5C).

Due to the intrinsic fluorescence of cyanine conjugates, we were able to determine their intracellular localization. Compounds **8b**, **8c**, and **8d** exhibited a high degree of co-localization with mitochondria (Pearson coefficients 0.63, 0.58, and 0.77, respectively) (Fig. 5D), whereas compound **8a** showed only weak co-localization (Pearson coefficient 0.44) (Fig. 5D).

Furthermore, like **1e**, cyanine conjugates induced general activation of autophagy in cells, as indicated by an increased pAMPK/AMPK_total_ ratio and increased ULK1 phosphorylation at S317 and S555. This evidence is consistent with these compounds functioning through the same autophagy activation pathway as the lead parent compound **1e**. Interestingly, all studied conjugates exhibited a high degree of co-localization with lysosomes (Pearson coefficients: **8a**=0.57, **8b**=0.52, **8c**=0.81, and **8d**=0.51) (Fig. 5E), which further supports their involvement in autophagy-associated processes. Thus, the obtained data indicate that the synthesized conjugates are accumulated in mitochondria and promote their degradation through mitophagy.

Conjugation of the lead compound with the Cy5 dye selectively enhanced the mitophagy-inducing capacity of **1e**, in contrast to Cy3 or TPP⁺ modifications, without altering AMPK/ULK1 signaling and cytotoxicity levels. Thus, Cy5-mediated targeting emerges as a functionally productive strategy that amplifies mitophagy without altering the core autophagy activation pathway.

### Mitochondrial targeting increases geroprotective properties of autophagy activators at lower concentrations

A toxicity analysis of the discovered effective activators of auto- and mitophagy was conducted to evaluate their potential application in living systems. Compound **1e** exhibited the highest cytotoxicity, with a half-maximal cytotoxic concentration (CC_50_) at 131 μM (Fig. 6A,B). According to data on autophagy and mitophagy activation obtained using fluorescent reporters, significant activity of this compound was observed at a concentration of 25 μM (Fig. 2A). Thus, the difference between the toxic and active concentrations is more than 4-fold.

**Fig. 6.**
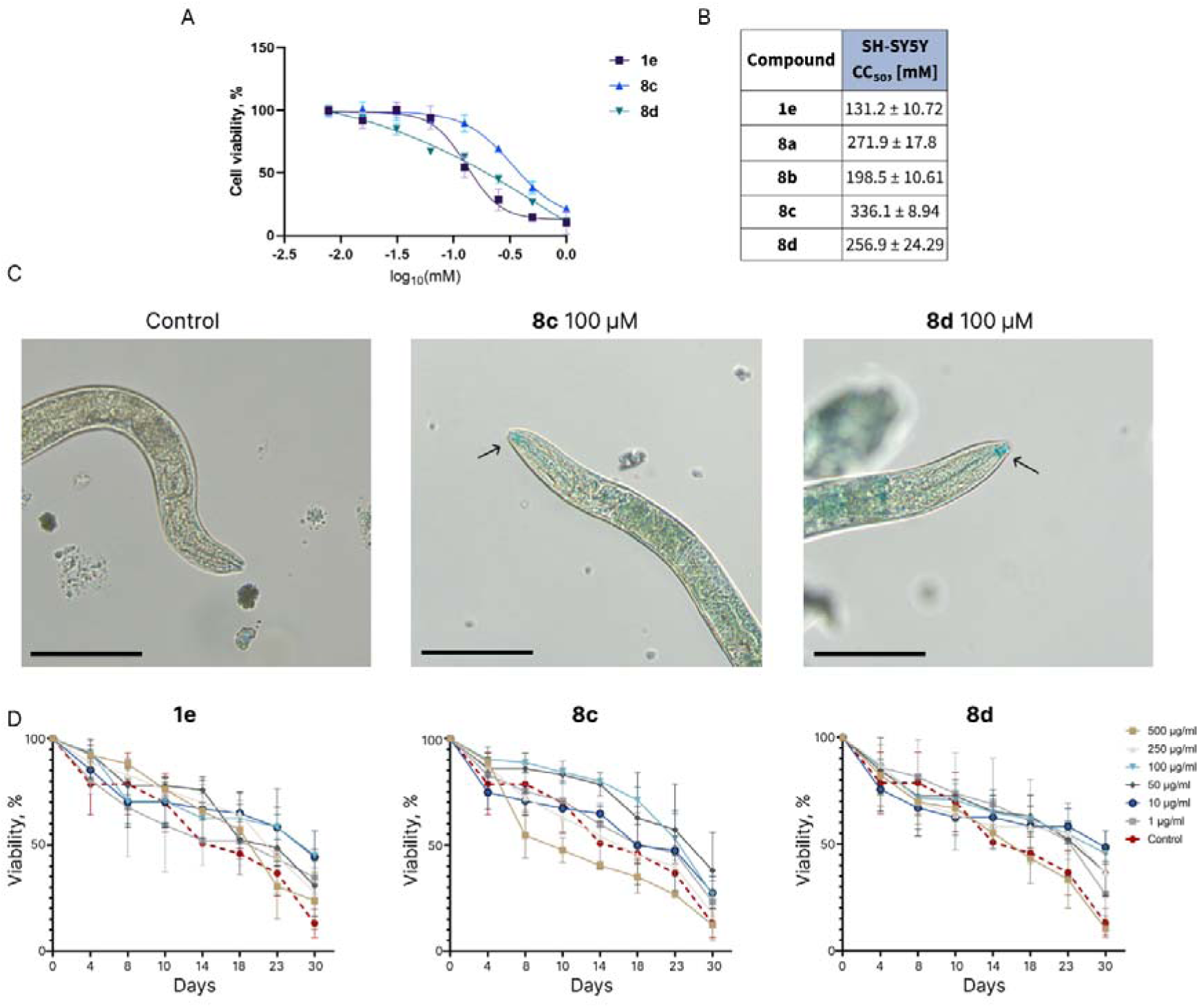
Investigation of the influence of **1e, 8c,** and **8d** on longevity of C*aenorhabditis elegans* A. Viability of human neuroblastoma (SH-SY5Y) cells incubated with **1e**, **8c**, and **8d** for 24 hours. MTT assay, data are the mean ± SD. B. CC_50_ values for SH-SY5Y cell line. C. Microphotographs of C*aenorhabditis elegans* after 17 days of treatment with **8c** and **8d**. Black bars indicate 100 μm. D. *Caenorhabditis elegans* lifespan under the treatment with **1e**, **8c**, and **8d**. Results are expressed as the average ± standard deviation (SEM) from 3 independent replicates.

Conjugation with cyanine dyes significantly reduced the toxicity of the compounds, with the CC_50_ falling within the 200–340 μM range, thus, for compounds **8a–d**, the difference between the active and toxic concentrations is 8–12-fold (Fig. 6A,B, Supp. Fig. 5 A,B).

As mitophagy activators are considered to be promising geroprotective molecules [18], we evaluated the impact of lead compounds **1e**, **8c**, and **8d** on nematode lifespan. Once the nematodes had reached adulthood, they were incubated in a medium containing various concentrations of the test substances: 1, 10, 50, 100, 250, and 500 μg/ml. The number of surviving nematodes was then evaluated over a period of 30 days.

When compounds **8c** and **8d** were incubated with the nematodes, accumulation of the compounds was observed in the peripharyngeal nerve ring (Fig. 6C). This indicates that the nematodes were actively absorbing the compounds, as well as that the compounds have penetrated neuronal cells.

Compound **1e** significantly extended the lifespan of *C. elegans* at all tested concentrations (Fig. 6D, Supplementary Table 1). The greatest effect—a 26.4% increase in lifespan—was observed at a concentration of 500 μg/ml. Compounds **8c** and **8d** also had a positive impact on nematode lifespan; however, their maximal effects were achieved at lower concentrations than that of **1e**. For compound **8d**, the optimal concentration was 100 μg/ml, at which lifespan was significantly extended by 30.2%. Compound **8c** was most effective at an even lower concentration of 50 μg/ml, resulting in a maximal lifespan extension of 32.1%. Notably, at concentrations above these optimal levels, the efficacy of compounds **8c** and **8d** decreased, which may be attributed to reduced compound solubility.

The lifespan extension observed in *C. elegans* following treatment with **1e** and Cy5-bearing conjugates **8c** and **8d** is consistent with their mitophagy-activating properties and reduced cytotoxicity. Notably, these conjugates achieved maximal geroprotective effects at lower concentrations than the parent compound, suggesting that improved mitochondrial targeting enhances biological efficiency *in vivo*.

## Discussion

In this study, we identified a novel class of small-molecule autophagy and mitophagy inducers based on a benzo[4,5]imidazo[1,2-c]pyrimidinone scaffold. Among the 27 compounds evaluated, 7-O-alkylated benzo[4,5]imidazo[1,2-c]pyrimidinonyl derivatives **1** of the 2’-deoxy series exhibited the most pronounced activity, with autophagy induction showing a strong dependence on the length of the alkyl substituent. A clear bell-shaped structure–activity relationship (SAR) was observed, with maximal activity achieved for the *n*-hexyl derivative **1e**.

This narrow and well-defined dependence on lipophilic tail length distinguishes this molecular series from many previously described phenotypic autophagy modulators, whose activity often lacks clear SAR trends.

Mechanistically, we demonstrate that the lead compound **1e** activates autophagy predominantly through the AMPK/ULK1-pathway. Treatment with **1e** resulted in robust AMPK activation and increased phosphorylation of ULK1 at S317 and S555, while the inhibitory AKT/TSC2/mTOR pathway remained largely intact. These findings indicate that autophagy induction occurs primarily through energy-stress–associated signaling rather than global suppression of anabolic pathways. Importantly, although AMPK activation accompanied autophagy induction, residual activity of **1e** in AMPK-deficient cells suggests that AMPK is not its exclusive target. This observation points to the involvement of additional upstream signals, potentially linked to mitochondrial functional perturbation or membrane-associated stress, which converge on the autophagy machinery.

In addition to general autophagy, we identified that **1e** exhibited strong mitophagy-inducing properties. Multiple independent assays demonstrated enhanced mitochondrial phospho-ubiquitination, increased mitochondrial delivery to lysosomes, and progressive mitochondrial degradation. Ultrastructural analysis further confirmed the accumulation of autophagosomes and autolysosomes containing fragmented mitochondria. Notably, although phospho-ubiquitin accumulation is classically associated with PINK1/Parkin-mediated mitophagy, mitochondrial clearance induced by **1e** proceeded efficiently in PRKN-deficient cells. This finding indicates that mitophagy activation by this compound is largely Parkin-independent, consistent with emerging models in which ubiquitin signaling cooperates with AMPK–ULK1 activation [5,49] and receptor-mediated pathways to promote mitochondrial turnover [50–52].

The ability of **1e** to selectively enhance degradation of mutant huntingtin aggregates further underscores the potential clinical relevance of the induced autophagic flux. Preferential clearance of the aggregation-prone protein relative to its soluble counterpart suggests that this compound may enhance the capacity of the autophagy system to process aberrant or damaged substrates, a property of particular importance for neurodegenerative disease contexts. Together with the observed neuronal accumulation of cyanine-conjugated derivatives in *C. elegans*, these findings highlight the neuroprotective potential of this scaffold.

To improve mitochondrial targeting and pharmacological properties, we explored conjugation of the identified lead compound with established mitochondria-directing moieties. Triphenylphosphonium conjugation yielded limited enhancement of mitophagy and was associated with increased cytotoxicity in the case of a long decylene linker (as in **6g**), consistent with excessive mitochondrial membrane accumulation. In contrast, conjugation with cyanine dyes—particularly Cy5—resulted in a marked enhancement of mitophagy-activating properties while simultaneously reducing cellular toxicity. Cy5-bearing conjugates displayed efficient mitochondrial localization, strong colocalization with lysosomes, and preserved activation of the AMPK/ULK1 pathway, indicating that mitochondrial targeting augments but does not alter the fundamental mechanism of action.

The functional significance of enhanced mitophagy activation was further supported by *in vivo* experiments in *Caenorhabditis elegans*. Both **1e** and its Cy5 conjugates significantly extended nematode lifespan, with the latter achieving maximal effects at substantially lower concentrations. The most potent conjugate, **8c**, increased lifespan by more than 30%, an effect comparable to or exceeding that reported for recently identified autophagy-based candidates in geroprotectors such as AA-20 [53], proAX [54], ribavirin [55], neoagarotetraose [56], and neomangiferin [57], thus suggesting therapeutic potential of the identified autophagy inducers.

### Conclusion

In this study, we identify a 2-(2′-deoxy-β-D-ribofuranosyl)-benzo[4,5]imidazo[1,2-c]pyrimidinone scaffold as an effective modulator of autophagy, with activity strongly dependent on the substituent at the 7-hydroxyl group. The lead compound enhances autophagy and mitophagy predominantly through activation of the AMPK–ULK1 pathway in a Parkin-independent manner, while promoting autophagosome formation and the clearance of aggregation-prone mutant huntingtin.

Conjugation of the lead compound with the mitochondria-targeting Cy5 dye further potentiates mitophagy induction, likely via preferential mitochondrial accumulation, while reducing cytotoxicity and significantly extending *C. elegans* lifespan at lower concentrations compared to the unconjugated compound. These findings validate mitochondria-targeted conjugation as an effective strategy to improve the biological performance of autophagy modulators.

Overall, our results define a promising chemical scaffold for the development of novel auto- and mitophagy activators and provide *in vivo* evidence supporting their potential relevance for modulating proteostasis and mitochondrial quality control, with possible implications for geroprotective interventions.

## Acknowledgements

We are grateful to the Moscow State University Development Program for providing access to the CelenaX High Content Imaging System. We thank Subdiffractional microscopy and spectroscopy core facility at Moscow state university for granting access to the equipment. We thank Alexander Korshun for the fruitful discussion.

## Data Availability Statement

The original contributions presented in the study are included in the article/Supplementary Material; further inquiries can be directed to the corresponding author.

## Institutional Review Board Statement

All studies conformed to the Guide for the Care and Use of Laboratory Animals, published by the United States National Institutes of Health (Publication No. 85-23, revised 1996) and approved by the Moscow Institute of Physics and Technology Life Science Center Provisional Animal Care and Research Procedures Committee, Protocol #A2-2012-09-02

