## supplementary file for "Cyanine dye conjugates of a 2’-deoxycytidine-based auto- and mitophagy activator extend *Caenorhabditis elegan*s lifespan"

[1]Belozersky Institute of Physico-Chemical Biology, Lomonosov Moscow State University, Moscow, Russia
[2]Chemical Department, Lomonosov Moscow State University, Moscow, Russia
[3]Shemyakin-Ovchinnikov Institute of Bioorganic Chemistry, Russian Academy of Sciences, Moscow, Russia
[4]Faculty of Fundamental Medicine, M.V. Lomonosov Moscow State University, Moscow, Russia
[5]Lomonosov Institute of Fine Chemical Technologies, MIREA—Russian Technological University, 119571 Moscow, Russia
[6]Frumkin Institute of Physical Chemistry and Electrochemistry, Russian Academy of Science, 119071 Moscow, Russia
[7]Peoples' friendship university of Russia named after Patrice Lumumba, Moscow, Russia
[8] Educational Resource Center for Cellular Technologies, RUDN University, 117198 Moscow, Russia
[9] Moscow Center for Advanced Studies, Moscow, Russia
†Ekaterina A. Guseva and Polina N. Kamzeeva have contributed equally to this work.
Full list of author information is available at the end of the article

### Table of Contents

**Chemistry** ........ **S2**

**Supplementary Figure 1** ........ **S7**

**Supplementary Figure 2** ........ **S8**

**Supplementary Figure 3** ........ **S9**

**Supplementary Figure 4** ........ **S9**

**Supplementary Figure 5** ........ **S9**

**Supplementary Table 1** ........ **S10**

**NMR spectra** ........ **S11**

**HPLC data** ........ **S36**

**References** ........ **S47**

### Chemistry

All reagents were commercially available unless otherwise mentioned and used without further purification. All solvents were purchased from commercial sources. Thin-layer chromatography (TLC) was performed on plates (Merck) precoated with silica gel (60 µm, F254) and visualized using UV light (254 and 365 nm). Column chromatography (CC) was performed on silica gel (0.040–0.063 mm, Merck, Germany). $^{1}$H, $^{13}$C, and $^{31}$P NMR spectra were recorded on a Bruker Avance III 600 spectrometer (Germany) at 600, 150, and 243 MHz, respectively. Chemical shifts are reported in δ (ppm) units using residual $^{1}$H signal from deuterated DMSO as reference. Multiplicity is reported using the following abbreviations: s (singlet), d (doublet), t (triplet), m (multiplet), br (broad). Coupling constants (J) are given in Hz. ESI HR mass spectra were acquired on a Thermo Scientific LTQ Orbitrap hybrid instrument (Thermo Electron Corp., Germany) in continuous-flow direct sample infusion (positive ion mode).

Purification of **8** was performed on an Interchim Puriflash 4250 preparative chromatograph (France) using a VDSpher 100 C18-E 250x20 mm, 10 µm column. MeOH and aqueous 0.2% trifluoroacetic acid were used as eluents. UV-vis detection was achieved at 205, 254, 350, 550 (**8a,b**), and 600 (**8c,d**) nm. A linear gradient from 5 to 95% of organic solvent in 10 min, followed by 4 min of 95% MeOH at a 20 mL/min flow rate, was used. Retention time ($R_t$) was 11.2 (**8a**), 11.4 (**8b**), 11.4 (**8c**), and 12.2 (**8d**) min.

Analytical HPLC analysis was performed on a Thermo Ultimate 3000 HPLC biocompatible system (USA) with DAD equipped with a Phenomenex Luna 3 µm C18(2) 4.6 x 75 mm column. For all samples except **7e**, the eluents were A - mQ water (Millipore), B - acetonitrile (HPLC grade, Fisher scientific). For compound **7e**, 0.1% (v/v) trifluoroacetic acid (HPLC grade, Fisher Scientific) was added to acetonitrile. During analysis, the solvent composition was linearly ramped from 5% B to 95% B in 5 minutes, held at 95% B for 2 minutes, and then returned to initial conditions (5% B) in 5 minutes at a flow rate of 1.2 ml/min. Before injection, the column was equilibrated for 2 minutes at initial conditions. The injection volume was 20 µL. The UV chromatogram was recorded at 210 nm. Retention time ($R_t$) is provided in the characteristics of the compounds.

Cyanine3 azide and cyanine5 azide were purchased from Lumiprobe (Russia). Compound **6**, (3-hydroxypropyl)triphenylphosphonium bromide, (6-hydroxyhexyl)triphenylphosphonium bromide, and (10-hydroxydecyl)triphenylphosphonium bromide were synthesized according to the literature [1–4].

General procedure for the preparation of 7-*O*-alkylated benzo[4,5]imidazo[1,2-c]pyrimidin-1(2H)-onyl derivatives of the 2'-deoxy series

To a stirred solution of **1b** (0.12 mmol), $PPh_3$ (2 eq.) and the corresponding alcohol (1.3 eq.) in dry $CH_2Cl_2$ (6 mL), DIAD (2 eq.) was added at 0°C. The reaction mixture was stirred at room temperature overnight and concentrated *in vacuo*. The residue was dissolved in a mixture of $CH_3COOH/H_2O$ (2 mL, 1:9, v:v) and kept at 50 °C for 8 h. After concentration *in vacuo* and co-evaporation with toluene (3 x 10 mL), the residue was purified using column chromatography on silica gel (0→6% $CH_3OH$ in $CH_2Cl_2$ for **6a-d** and **7a**,**b**, and 0→50% $CH_3OH$ in $CH_2Cl_2$ for **6e-g**), yielding **6a–g** and **7a,b**.

*2-((2R,4S,5R)-4-hydroxy-5-(hydroxymethyl)tetrahydrofuran-2-yl)-7-(pentyloxy)benzo[4,5]imidazo[1,2-c]pyrimidin-1(2H)-one* **6a**

Starting from *n*-propanol, this derivative was prepared as a brownish foam with a yield of 90%. $R_t$ = 3.87 min. $^{1}$H NMR (600 MHz, DMSO-$d_6$): 7.86 (d, J = 8.0 Hz, 1H), 7.85 (d, J = 2.5 Hz, 1H), 7.62 (d, J = 8.8 Hz, 1H), 7.08 (dd, J = 8.8 Hz, J = 2.5 Hz, 1H), 6.70 (d, J = 8.0 Hz, 1H), 6.46 (t, J = 6.7Hz, 1H), 5.30 (d, J = 4.2 Hz, 1H), 5.07 (t, J = 5.2 Hz, 1H), 4.33-4.30 (m, 1H), 4.07-4.01 (m, 2H), 3.89-3.86 (m, 1H), 3.68-3.59 (m, 2H), 2.28-2.20 (m, 2H), 1.80-1.74 (m, 2H), 1.47-1.41 (m, 2H), 1.40-1.33 (m, 2H), 0.91 (t, J = 7.3 Hz, 3H). $^{13}$C NMR (150 MHz, DMSO-$d_6$): 155.2, 147.3,

146.7, 138.1, 131.2, 130.3, 119.1, 114.5, 99.0, 98.0, 87.6, 84.8, 70.2, 68.0, 61.1, 40.0, 28.2, 27.6, 21.7, 13.7. HRMS (ESI) m/z: calcd for $C_{20}H_{26}N_3O_5^+$ [M+H]$^+$: 388.1867; found 388.1878.

*2-((2R,4S,5R)-4-hydroxy-5-(hydroxymethyl)tetrahydrofuran-2-yl)-7-(isopentyloxy)benzo[4,5]imidazo[1,2-c]pyrimidin-1(2H)-one* **6b**

Starting from *iso*-propanol, this derivative was prepared as a beige foam with a yield of 88%. $R_t$ = 3.82 min. $^1$H NMR (600 MHz, DMSO-$d_6$): 7.86 (d, J = 8.1 Hz, 1H), 7.84 (d, J = 2.1 Hz, 1H), 7.61 (d, J = 8.8 Hz, 1H), 7.07 (dd, J = 8.1 Hz, J = 2.1 Hz, 1H), 6.69 (d, J = 8.0 Hz, 1H), 6.50 (t, J = 6.7Hz, 1H), 5.31 (d, J = 3.7 Hz, 1H), 5.08 (t, J = 5.2 Hz, 1H), 4.34-4.30 (m, 1H), 4.08-4.04 (m, 2H), 3.90-3.87 (m, 1H), 3.70-3.60 (m, 2H), 2.28-2.20 (m, 2H), 1.86-1.78 (m, 1H), 1.68-1.63 (m, 2H), 0.95 (d, J = 6.7 Hz, 6H). $^{13}$C NMR (150 MHz, DMSO-$d_6$): 155.2, 147.3, 146.6, 138.1, 131.2, 130.3, 119.1, 114.5, 99.1, 98.0, 87.6, 84.8, 70.2, 66.4, 61.1, 40.1, 37.3, 24.5, 22.3 (2C). HRMS (ESI) m/z: calcd for $C_{20}H_{26}N_3O_5^+$ [M+H]$^+$: 388.1867; found 388.1880.

*2-((2R,4S,5R)-4-hydroxy-5-(hydroxymethyl)tetrahydrofuran-2-yl)-7-phenethoxybenzo[4,5]imidazo[1,2-c]pyrimidin-1(2H)-one* **6c**

Starting from 2-phenylethan-1-ol, this derivative was prepared as a beige foam with a yield of 88%. $R_t$ = 3.71 min. $^1$H NMR (600 MHz, DMSO-$d_6$): 7.87 (d, J = 8.1 Hz, 1H), 7.85 (d, J = 2.5 Hz, 1H), 7.62 (d, J = 8.8 Hz, 1H), 7.37-7.30 (m, 4H), 7.25-7.21 (m, 1H), 7.08 (dd, J = 8.1 Hz, J = 2.5 Hz, 1H), 6.69 (d, J = 8.0 Hz, 1H), 6.45 (t, J = 6.7Hz, 1H), 5.31 (d, J = 4.2 Hz, 1H), 5.08 (t, J = 5.2 Hz, 1H), 4.34-4.30 (m, 1H), 4.27 (dt, J = 0.8 Hz, J = 6.8 Hz, 2H), 3.89-3.87 (m, 1H), 3.68-3.60 (m, 2H), 3.09 (t, J = 6.8 Hz, 2H), 2.27-2.20 (m, 2H). $^{13}$C NMR (150 MHz, DMSO-$d_6$): 154.9, 147.4, 146.6, 138.2, 138.2, 131.3, 130.3, 128.8 (2C), 128.2 (2C), 126.1, 119.1, 114.4, 99.2, 98.0, 87.6, 84.8, 70.2, 68.8, 61.1, 40.1, 34.8. HRMS (ESI) m/z: calcd for $C_{23}H_{24}N_3O_5^+$ [M+H]$^+$: 422.1710; found 422.1711.

*2-((2R,4S,5R)-4-hydroxy-5-(hydroxymethyl)tetrahydrofuran-2-yl)-7-(3-phenylpropoxy)benzo[4,5]imidazo[1,2-c]pyrimidin-1(2H)-one* **6d**

Starting from 3-phenylpropan-1-ol, this derivative was prepared as a brownish foam with a yield of 89%. $R_t$ = 3.95 min. $^1$H NMR (600 MHz, DMSO-$d_6$): 7.87 (d, J = 8.1 Hz, 1H), 7.84 (d, J = 2.5 Hz, 1H), 7.63 (d, J = 8.8 Hz, 1H), 7.30-7.23 (m, 4H), 7.20-7.16 (m, 1H), 7.10 (dd, J = 8.1 Hz, J = 2.5 Hz, 1H), 6.70 (d, J = 8.0 Hz, 1H), 6.46 (t, J = 6.7Hz, 1H), 5.30 (d, J = 4.2 Hz, 1H), 5.07 (t, J = 5.2 Hz, 1H), 4.33-4.30 (m, 1H), 4.07-4.01 (m, 2H), 3.89-3.87 (m, 1H), 3.68-3.59 (m, 2H), 2.81-2.77 (m, 2H), 2.28-2.20 (m, 2H), 2.11-2.05 (m, 2H). $^{13}$C NMR (150 MHz, DMSO-$d_6$): 155.1, 147.4, 146.6, 141.2, 138.2, 131.2, 130.3, 128.2 (4C), 125.7, 119.1, 114.5, 99.1, 98.0, 87.6, 84.8, 70.2, 67.3, 61.1, 40.0, 31.3, 30.2. HRMS (ESI) m/z: calcd for $C_{24}H_{26}N_3O_5^+$ [M+H]$^+$: 436.1867; found 436.1867.

*(3-((2-((2R,4S,5R)-4-hydroxy-5-(hydroxymethyl)tetrahydrofuran-2-yl)-1-oxo-1,2-dihydrobenzo[4,5]imidazo[1,2-c]pyrimidin-7-yl)oxy)propyl)triphenylphosphonium bromide* **6e**

Starting from (3-hydroxypropyl)triphenylphosphonium bromide, this derivative was prepared as a brownish foam with a yield of 71%. $R_t$ = 2.91 min. $^{31}$P NMR (243 MHz, DMSO-$d_6$): δ 24.6. $^1$H NMR (600 MHz, DMSO-$d_6$): δ 7.93-7.83 (m, 11H), 7.80-7.74 (m, 6H), 7.64 (d, J = 8.8 Hz, 1H), 7.10 (dd, J = 8.8 Hz, J = 2.3 Hz, 1H), 6.71 (d, J = 8.0 Hz, 1H), 6.46 (t, J = 6.8 Hz, 1H), 5.41 (d, J = 4.1 Hz, 1H), 5.18 (t, J = 5.0 Hz, 1H), 4.36-4.32 (m, 1H), 4.20 (t, J = 5.7 Hz, 2H), 3.91-3.87 (m, 1H), 3.85-3.78 (m, 2H), 3.68-3.59 (m, 2H), 2.28-2.20 (m, 2H), 2.09-2.02 (m, 2H). $^{13}$C NMR (151 MHz, DMSO-$d_6$): δ 154.5, 147.5, 146.6, 138.4, 134.8 (3C), 133.5 (d, $^3J_{P,C}$ = 10.0 Hz, 6C), 131.5, 130.2, 130.2 (d, $^2J_{P,C}$ = 12.5 Hz, 6C), 119.1, 118.3 (d, $^2J_{P,C}$ = 86.1 Hz, 3H), 114.4, 99.4, 98.0, 87.7, 84.9, 70.2, 67.3 (d, $^3J_{P,C}$ = 18.0 Hz, 1C), 61.1, 40.0, 22.1 (d, $^3J_{P,C}$ = 2.0 Hz, 1C), 17.6 (d, $^1J_{P,C}$ = 52.7 Hz, 1C). HRMS (ESI) m/z: calcd for $C_{36}H_{35}N_3O_5P^+$ [M-Br$^-$]$^+$: 620.2309; found 620.2331.

*(6-((2-((2R,4S,5R)-4-hydroxy-5-(hydroxymethyl)tetrahydrofuran-2-yl)-1-oxo-1,2-dihydrobenzo[4,5]imidazo[1,2-c]pyrimidin-7-yl)oxy)hexyl)triphenylphosphonium bromide* **6f**

Starting from (6-hydroxyhexyl)triphenylphosphonium bromide, this derivative was prepared as a brownish foam with a yield of 76%. $R_t$ = 3.13 min. $^{31}$P NMR (243 MHz, DMSO-$d_6$): δ 24.2. $^1$H NMR (600 MHz, DMSO-$d_6$): δ 7.91-7.86 (m, 4H), 7.84-7.74 (m, 13H), 7.62 (d, J = 8.8 Hz, 1H), 7.05 (dd, J = 8.8 Hz, J = 2.5 Hz, 1H), 6.70 (d, J = 8.0 Hz, 1H), 6.45 (t, J = 6.8 Hz, 1H), 5.33 (br s, 1H), 5.09 (br s, 1H), 4.35-4.31 (m, 1H), 4.00 (t, J = 6.5 Hz, 2H), 3.90-3.87 (m, 1H), 3.67-3.55 (m, 4H), 2.27-2.20 (m, 2H), 1.74-1.67 (m, 2H), 1.62-1.51 (m, 4H), 1.51-1.44 (m, 2H). $^{13}$C NMR (151 MHz, DMSO-$d_6$): δ 155.1, 147.3, 146.6, 138.1, 134.7 (3C), 133.5 (d, $^3J_{P,C}$ = 9.9 Hz, 6C), 131.2, 130.3, 130.1 (d, $^2J_{P,C}$ = 12.3 Hz, 6C), 119.1, 118.4 (d, $^2J_{P,C}$ = 85.6 Hz, 3H), 114.5, 99.0, 98.0, 87.6, 84.8, 70.2, 67.8, 61.1, 40.0, 29.4 (d, $^2J_{P,C}$ = 16.6 Hz, 1C), 28.1, 24.5, 21.6 (d, $^3J_{P,C}$ = 3.3 Hz, 1C), 20.1 (d, $^1J_{P,C}$ = 50.0 Hz, 1C). HRMS (ESI) m/z: calcd for $C_{39}H_{41}N_3O_5P^+$ [M-Br$^-$]$^+$: 662.2778; found 662.2796.

*(10-((2-((2R,4S,5R)-4-hydroxy-5-(hydroxymethyl)tetrahydrofuran-2-yl)-1-oxo-1,2-dihydrobenzo[4,5]imidazo[1,2-c]pyrimidin-7-yl)oxy)decyl)triphenylphosphonium bromide* **6g**

Starting from (10-hydroxydecyl)triphenylphosphonium bromide, this derivative was prepared as a brownish foam with a yield of 78%. $R_t$ = 2.03 min. $^{31}$P NMR (243 MHz, DMSO-$d_6$): δ 24.1. $^1$H NMR (600 MHz, DMSO-$d_6$): δ 7.91-7.86 (m, 3H), 7.84 (d, J = 2.5 Hz, 1H), 7.83-7.7.4 (m, 13H), 7.62 (d, J = 8.8 Hz, 1H), 7.07 (d, J = 8.8 Hz, J = 2.5 Hz, 1H), 6.70 (d, J = 8.0 Hz, 1H), 6.46 (t, J = 6.7 Hz, 1H), 5.31 (br s, 1H), 5.07 (br s, 1H), 4.34-4.30 (m, 1H), 4.03 (t, J = 6.4 Hz, 2H), 3.90-3.86 (m, 1H), 3.65 (dd, J = 11.7 Hz, J = 3.5 Hz, 1H), 3.61 (dd, J = 11.7 Hz, J = 3.6 Hz, 1H), 3.58-3.52 (m, 2H), 2.27-2.19 (m, 2H), 1.78-1.71 (m, 2H), 1.57-1.48 (m, 2H), 1.48-1.39 (m, 4H), 1.33-1.21 (m, 8H). $^{13}$C NMR (151 MHz, DMSO-$d_6$): δ 155.8, 147.9, 147.3, 138.7, 135.4 (d, $^4J_{P,C}$ = 1.6 Hz, 3C), 134.1 (d, $^3J_{P,C}$ = 10.0 Hz, 6C),131.8, 130.9, 130.7 (d, $^2J_{P,C}$ = 12.4 Hz, 6C), 119.7, 119.1 (d, $^2J_{P,C}$ = 85.6 Hz, 3H), 115.1, 99.7, 98.6, 88.2, 85.4, 70.8, 68.6, 61.7, 40.6, 30.3 (d, $^2J_{P,C}$ = 16.4 Hz, 1C), 29.2, 29.1 (2C), 29.0, 28.4, 25.9, 22.2 (d, $^3J_{P,C}$ = 4.0 Hz, 1C), 20.7 (d, $^1J_{P,C}$ = 49.9 Hz, 1C). HRMS (ESI) m/z: calcd for $C_{43}H_{49}N_3O_5P^+$ [M-Br$^-$]$^+$: 718.3404; found 718.3420.

*2-((2R,4S,5R)-4-hydroxy-5-(hydroxymethyl)tetrahydrofuran-2-yl)-7-(prop-2-yn-1-yloxy)benzo[4,5]imidazo[1,2-c]pyrimidin-1(2H)-one* **7a**

Starting from prop-2-yn-1-ol, this derivative was prepared as a brownish foam with a yield of 72%. $^1$H NMR (600 MHz, DMSO-$d_6$): 7.97 (d, J = 2.4 Hz, 1H), 7.89 (d, J = 8.0 Hz, 1H), 7.66 (d, J = 8.8 Hz, 1H), 7.14 (dd, J = 8.8 Hz, J = 2.4 Hz, 1H), 6.71 (d, J = 8.0 Hz, 1H), 6.47 (t, J = 6.7 Hz, 1H), 5.31 (br s, 1H), 5.08 (br s, 1H), 4.88 (d, J = 2.1 Hz, 2H), 4.34-4.30 (m, 1H), 3.90-3.86 (m, 1H), 3.69-3.60 (m, 2H), 3.57 (t, J = 2.1 Hz, 1H), 2.30-2.20 (m, 2H). $^{13}$C NMR (150 MHz, DMSO-$d_6$): 153.6, 147.7, 146.6, 138.7, 131.4, 130.1, 119.1, 114.8, 100.1, 97.9, 87.6, 84.8, 79.1, 78.2, 70.1, 61.1, 56.1, 40.1. HRMS (ESI) m/z: calcd for $C_{18}H_{18}N_3O_5^+$ [M+H]$^+$: 356.1241; found 356.1241.

*7-(hex-5-yn-1-yloxy)-2-((2R,4S,5R)-4-hydroxy-5-(hydroxymethyl)tetrahydrofuran-2-yl)benzo[4,5]imidazo[1,2-c]pyrimidin-1(2H)-one* **7b**

Starting from hex-5-yn-1-ol, this derivative was prepared as a brownish foam with a yield of 72%. $^1$H NMR (600 MHz, DMSO-$d_6$): 7.86 (d, J = 8.1 Hz, 1H), 7.84 (d, J = 2.5 Hz, 1H), 7.62 (d, J = 8.8 Hz, 1H), 7.08 (dd, J = 8.8 Hz, J = 2.5 Hz, 1H), 6.69 (d, J = 8.0 Hz, 1H), 6.50 (t, J = 6.7 Hz, 1H), 5.31 (d, J = 4.2 Hz, 1H), 5.08 (t, J = 5.2 Hz, 1H), 4.34-4.30 (m, 1H), 4.09-4.03 (m, 2H), 3.90-3.87 (m, 1H), 3.68-3.60 (m, 2H), 2.76 (t, J = 2.6 Hz, 1H), 2.39-2.20 (m, 4H), 1.88-1.82 (m, 2H), 1.68-1.61 (m, 2H). $^{13}$C NMR (150 MHz, DMSO-$d_6$): 155.1, 147.4, 146.6, 138.2, 131.2, 130.3, 119.1, 114.5, 99.1, 98.0, 87.6, 84.8, 84.1, 71.2, 70.2, 67.5, 61.1, 40.1, 27.7, 24.5, 17.3. HRMS (ESI) m/z: calcd for $C_{21}H_{24}N_3O_5^+$ [M+H]$^+$: 398.1710; found 398.1710.

General procedure for the preparation of cyanine-conjugated benzo[4,5]imidazo[1,2-c]pyrimidin-1(2H)-onyl derivatives **8**

To a stirred solution of the corresponding cyanine azide (0.017 mmol), compound **7** (1.2 eq.), and sodium ascorbate (10 eq.) in 65% aqueous DSMO (1.5 mL), a solution of 100 mM $CuSO_4$/TBTA (0.1 eq.) in 50% aqueous DMSO was added and stirred for 8 hours at room temperature. After an addition of an aqueous solution of EDTA (0.5 M, 1.3 eq.), the resulting mixture was lyophilized and purified by preparative HPLC, affording the target compounds as a 2,2,2-trifluoroacetate salt.

*1-(6-((3-(4-(((2-((2R,4S,5R)-4-hydroxy-5-(hydroxymethyl)tetrahydrofuran-2-yl)-1-oxo-1,2-dihydrobenzo[4,5]imidazo[1,2-c]pyrimidin-7-yl)oxy)methyl)-1H-1,2,3-triazol-1-yl)propyl)amino)-6-oxohexyl)-3,3-dimethyl-2-((E)-3-((E)-1,3,3-trimethylindolin-2-ylidene)prop-1-en-1-yl)-3H-indol-1-ium 2,2,2-trifluoroacetate* **8a**

Starting from **7a** and cyanine3, this derivative was prepared as a crimson amorphous solid with a yield of 49%. $R_t$ = 3.36 min. $^1$H NMR (600 MHz, DMSO-$d_6$): δ 8.33 (t, J = 13.5 Hz, 1H), 8.26 (s, 1H), 8.01-7.94 (m, 2H), 7.90 (d, J = 7.9 Hz, 1H), 7.68-7.57 (m, 3H), 7.47-7.36 (m, 4H), 7.32-7.23 (m, 2H), 7.17 (dd, J = 8.5 Hz, J = 2.5 Hz, 1H), 6.70 (d, J = 7.8 Hz, 1H), 6.51-6.42 (m, 3H), 5.22 (s, 2H), 4.39-4.31 (m, 3H), 4.10 (t, J = 6.7 Hz, 2H), 3.89-3.84 (m, 1H), 3.68-3.56 (m, 5H), 3.06-3.00 (m, 2H), 2.29-2.19 (m, 2H), 2.12-2.06 (m, 2H), 1.95-1.86 (m, 2H), 1.76-1.68 (m, 2H), 1.67 (s, 12H), 1.61-1.54 (m, 2H), 1.42-1.35 (m, 2H). HRMS (ESI) m/z: calcd for $C_{51}H_{60}N_9O_6^+$ [M-$CF_3COO^-$]$^+$: 894.4661; found 894.4675.

*1-(6-((3-(4-(4-((2-((2R,4S,5R)-4-hydroxy-5-(hydroxymethyl)tetrahydrofuran-2-yl)-1-oxo-1,2-dihydrobenzo[4,5]imidazo[1,2-c]pyrimidin-7-yl)oxy)butyl)-1H-1,2,3-triazol-1-yl)propyl)amino)-6-oxohexyl)-3,3-dimethyl-2-((E)-3-((E)-1,3,3-trimethylindolin-2-ylidene)prop-1-en-1-yl)-3H-indol-1-ium 2,2,2-trifluoroacetate* **8b**

Starting from **7b** and cyanine3, this derivative was prepared as a crimson amorphous solid with a yield of 51%. $R_t$ = 3.43 min. $^1$H NMR (600 MHz, DMSO-$d_6$): δ 8.33 (t, J = 13.5 Hz, 1H), 7.98-7.93 (m, 1H), 7.91-7.87 (m 2H), 7.84 (d, J = 2.3 Hz, 1H), 7.63-7.59 (m, 3H), 7.46-7.38 (m, 4H), 7.32-7.24 (m, 2H), 7.08 (dd, J = 8.8 Hz, J = 2.3 Hz, 1H), 6.69 (d, J = 8.1 Hz, 1H), 6.49-6.42 (m, 3H), 4.36-4.31 (m, 1H), 4.28 (t, J = 7.0 Hz, 2H), 4.09 (t, J = 7.5 Hz, 2H), 4.06 (t, J = 5.5 Hz, 2H), 3.88-3.84 (m, 1H), 3.66-3.60 (m, 5H), 3.03-2.98 (m, 2H), 2.68 (t, J = 7.0 Hz, 2H), 2.29-2.19 (m, 2H), 2.08 (t, J = 7.1 Hz, 2H), 1.92-1.85 (m, 2H), 1.84-1.74 (m, 4H), 1.75-1.69 (m, 2H), 1.67 (s, 12H), 1.60-1.54 (m, 2H), 1.42-1.35 (m, 2H). HRMS (ESI) m/z: calcd for $C_{54}H_{66}N_9O_6^+$ [M-$CF_3COO^-$]$^+$: 936.5131; found 936.5169.

*1-(6-((3-(4-(((2-((2R,4S,5R)-4-hydroxy-5-(hydroxymethyl)tetrahydrofuran-2-yl)-1-oxo-1,2-dihydrobenzo[4,5]imidazo[1,2-c]pyrimidin-7-yl)oxy)methyl)-1H-1,2,3-triazol-1-yl)propyl)amino)-6-oxohexyl)-3,3-dimethyl-2-((1E,3E)-5-((E)-1,3,3-trimethylindolin-2-ylidene)penta-1,3-dien-1-yl)-3H-indol-1-ium 2,2,2-trifluoroacetate* **8c**

Starting from **7a** and cyanine5, this derivative was prepared as a blue amorphous solid with a yield of 62%. $R_t$ = 3.54 min. $^1$H NMR (600 MHz, DMSO-$d_6$): δ 8.31 (t, J = 13.0 Hz, 2H), 8.25 (s, 1H), 7.98 (d, J = 2.4 Hz, 1H), 7.91-7.88 (m, 1H), 7.89 (d, J = 8.1 Hz, 1H), 7.64 (d, J = 8.7 Hz, 1H), 7.61-7.57 (m, 2H), 7.41-7.35 (m, 4H), 7.26-7.20 (m, 2H), 7.17 (dd, J = 8.9 Hz, J = 2.4 Hz, 1H), 6.70 (d, J = 8.1 Hz, 1H), 6.54 (t, J = 12.3 Hz, 1H), 6.46 (t, J = 6.7 Hz, 1H), 6.29 (d, J = 13.5 Hz, 1H), 6.23 (d, J = 13.6 Hz, 1H), 5.22 (s, 2H), 4.36 (t, J = 6.9 Hz, 2H), 4.34-4.31 (m, 1H), 4.08 (t, J = 7.1 Hz, 2H), 3.89-3.86 (m, 1H), 3.65 (dd, J = 11.8 Hz, J = 3.6 Hz, 1H), 3.62 (dd, J = 11.8 Hz, J = 3.4 Hz, 1H), 3.58 (s, 3H), 3.05-3.00 (m 2H), 2.27-2.20 (m, 2H), 2.07 (t, J = 7.2 Hz, 2H), 1.95-1.89 (m, 2H), 1.71-1.66 (m, 2H), 1.67 (s, 12H), 1.58-1.52 (m, 2H), 1.37-1.32 (m, 2H). HRMS (ESI) m/z: calcd for $C_{53}H_{62}N_9O_6^+$ [M-$CF_3COO^-$]$^+$: 920.4818; found 920.4830.

*1-(6-((3-(4-(4-((2-((2R,4S,5R)-4-hydroxy-5-(hydroxymethyl)tetrahydrofuran-2-yl)-1-oxo-1,2-dihydrobenzo[4,5]imidazo[1,2-c]pyrimidin-7-yl)oxy)butyl)-1H-1,2,3-triazol-1-yl)propyl)amino)-6-oxohexyl)-3,3-dimethyl-2-((1E,3E)-5-((E)-1,3,3-trimethylindolin-2-ylidene)penta-1,3-dien-1-yl)-3H-indol-1-ium 2,2,2-trifluoroacetate* **8d**

Starting from **7b** and cyanine5, this derivative was prepared as a blue amorphous solid with a yield of 48%. $R_t$ = 3.60 min. $^1$H NMR (600 MHz, DMSO-$d_6$): δ 8.31 (t, J = 12.9 Hz, 2H), 7.90-7.86 (m, 3H), 7.84 (d, J = 2.3 Hz, 1H), 7.63-7.57 (m, 3H), 7.41-7.34 (m, 4H), 7.26-7.20 (m, 2H), 7.07 (dd, J = 8.8 Hz, J = 2.3 Hz, 1H), 6.69 (d, J = 8.0 Hz, 1H), 6.54 (t, J = 12.2 Hz, 1H), 6.45 (t, J = 6.7 Hz, 1H), 6.29 (d, J = 13.5 Hz, 1H), 6.24 (d, J = 14.0 Hz, 1H), 4.35-4.30 (m, 1H), 4.28 (t, J = 6.9 Hz, 2H), 4.10-4.03 (m, 4H), 3.89-3.86 (m, 1H), 3.65 (dd, J = 12.0 Hz, J = 3.5 Hz, 1H), 3.62 (dd, J = 12.0 Hz, J = 3.6 Hz, 1H), 3.58 (s, 3H), 3.03-2.98 (m, 2H), 2.69 (t, J = 7.0 Hz, 2H), 2.26-2.20 (m, 2H), 2.06 (t, J = 7.0 Hz, 2H), 1.92-1.86 (m, 2H), 1.83-1.75 (m, 6H), 1.67 (s, 12H), 1.58-1.52 (m, 2H), 1.38-1.32 (m, 2H). HRMS (ESI) m/z: calcd for $C_{56}H_{68}N_9O_6^+$ $[M-CF_3COO^-]^+$: 962.5287; found 962.5314.

A

| Compound | 250 µM | 25 µM | 2.5 µM |
|---|---|---|---|
| 1a | -1.35 | -0.68 | -1.96 |
| 1b | 9.75 | 0.65 | 2.20 |
| 1c | 2.78 | 1.33 | 0.38 |
| 1d | 8.79 | 3.17 | 2.64 |
| 1e | 11.66 | 9.78 | 3.92 |
| 1f | 8.40 | -0.98 | -1.33 |
| 1g | 2.96 | 2.30 | -0.19 |
| 1h | -0.72 | 1.15 | -0.97 |
| 2a | 0.44 | -2.68 | -2.84 |
| 2b | 6.28 | 5.70 | 2.66 |
| 2c | -0.48 | 1.91 | 0.17 |
| 2d | 2.18 | -1.39 | 1.23 |
| 2e | 1.47 | 1.79 | 1.08 |
| 2f | -2.56 | 2.52 | 0.02 |
| 2g | -4.38 | -3.87 | -0.23 |
| 2h | -1.47 | 1.45 | -0.45 |
| 3a | -0.04 | 0.58 | 0.36 |
| 3b | -0.24 | -0.03 | -0.75 |
| 3c | 0.40 | -0.55 | -1.54 |
| 3d | 1.63 | 0.84 | 0.92 |
| 3e | 0.73 | 1.89 | 1.33 |
| 4a | -1.47 | 0.31 | -0.12 |
| 4b | 1.76 | 2.62 | -2.26 |
| 5a | -0.20 | 1.24 | -4.35 |
| 5b | 1.48 | 2.02 | 0.28 |
| 5c | 0.33 | 2.75 | 1.23 |
| 5d | 2.91 | 2.26 | -3.73 |

B

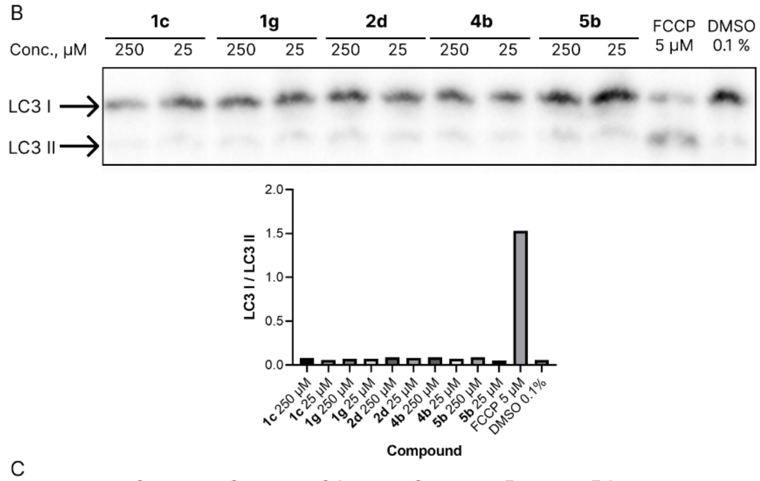

C

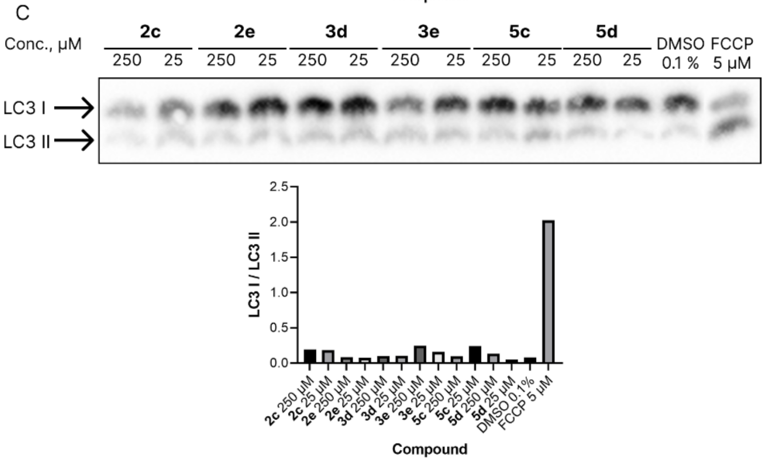

**Supplementary Fig. 1** A. Change in the RFP/GFP ratio relative to DMSO. Positive values indicate an increase in the level of autophagy, while negative values indicate a decrease. B. Western blotting on the key marker of autophagy (LC3) on SH-SY5Y when treated with **1c**, **1g**, **2d**, **4b**, **5b**, and bar graph indicating changes in LC3 forms ratio. C. Western blotting on the key marker of autophagy (LC3) on SH-SY5Y when treated with **2c, 2e, 3d, 3e, 5c, 5d**, and bar graph indicating changes in LC3 forms ratio.

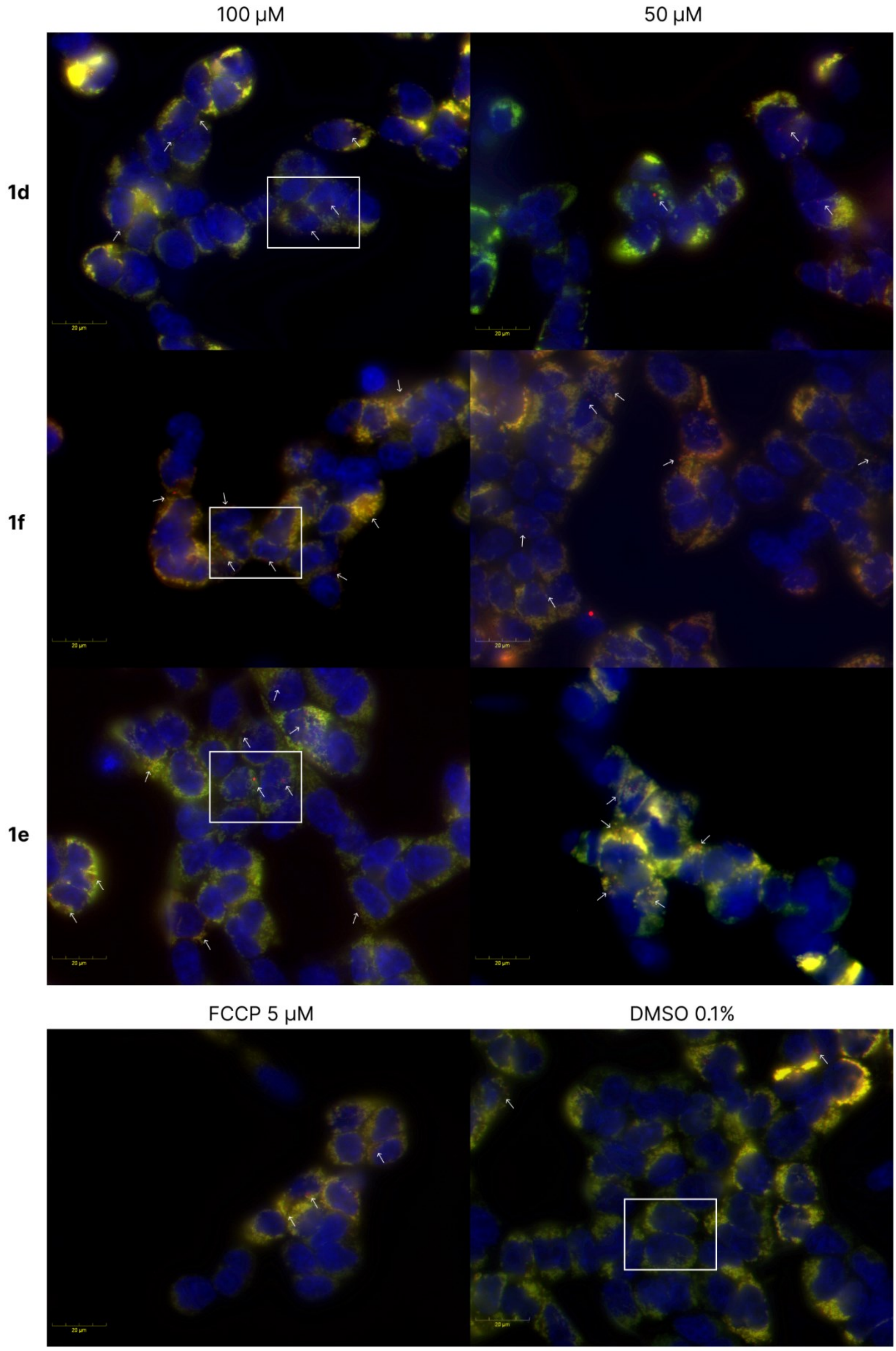

**Supplementary Fig. 2** Mitophagy induced by **1d-f** in wild-type HEK293T expressing TOM20MTS-mCherry-EGFP-Tet-On. Yellow bars indicate 20 μm. White arrows mark mitochondria in lysosomes. Nuclei were stained with Hoechst (blue). Cells were treated with doxycycline (2 ug/mL) 24 h before the experiment. White frames indicate fields, which are shown at higher magnification on Fig. 4D.

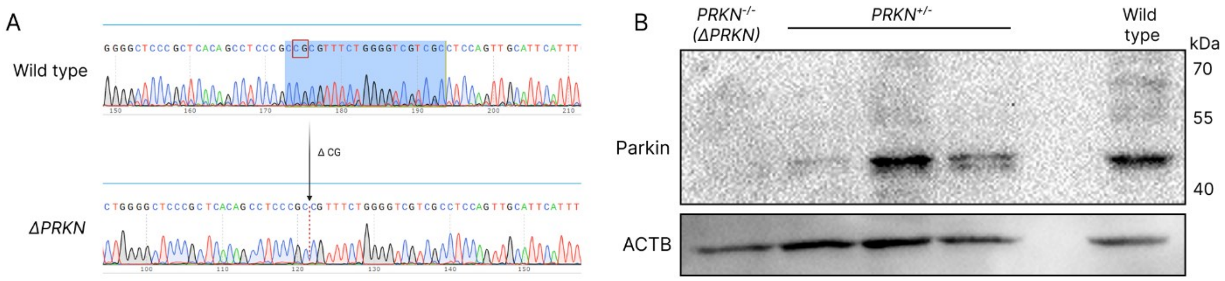

**Supplementary Fig. 3** Generation of *ΔPRKN* SH-SY5Y cell line. A. Sequences of the *PRKN* genome loci of generated *ΔPRKN* and wild-type SH-SY5Y cell lines. B. Western blotting with anti-Parkin antibodies of generated *ΔPRKN* (*PRKN*$^{-/-}$), *PRK*$^{+/-}$, and wild-type SH-SY5Y cell lines.

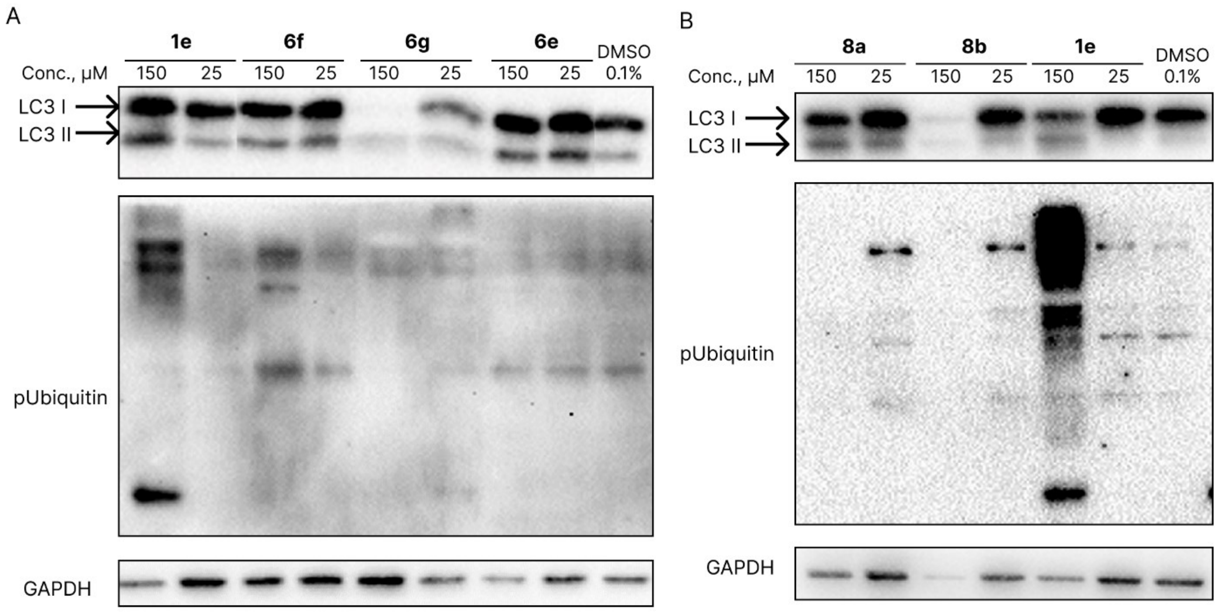

**Supplementary Fig. 4** Western blotting for the key markers of autophagy (LC3) and mitophagy (p-Ubiquitin) in SH-SY5Y cells under treatment with **8a** (A) and **8b** (B)**.** Data represent mean ± SD (n = 3). The results of ANOVA tests are indicated on the graph: * – p-value <0.05, ** – p-value <0.01, **** – p-value <0.0001.

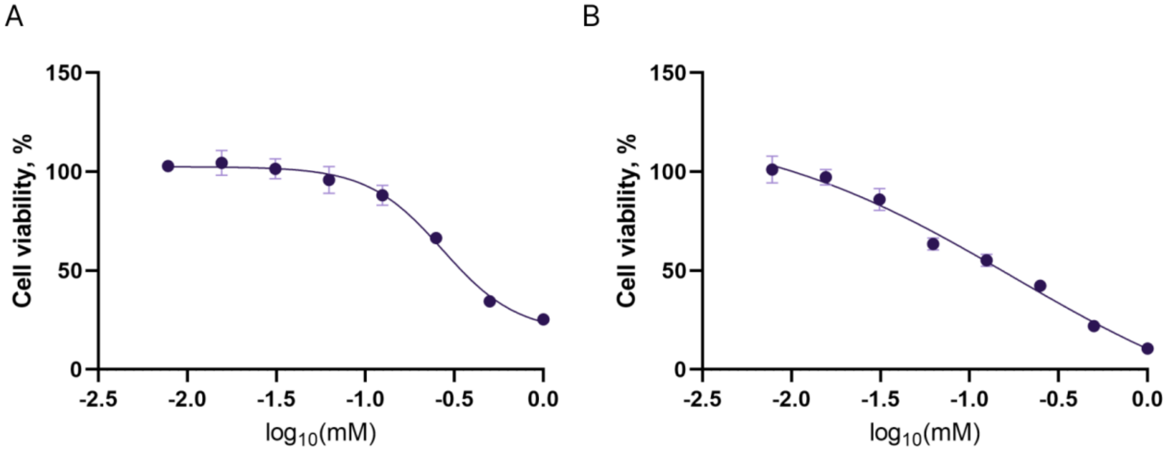

**Supplementary Fig. 5** Viability of human neuroblastoma (SH-SY5Y) cells incubated with **8a** (A) and **8b** (B) for 24 h. MTT assay, data are the mean ± SD.

**Supplementary Table 1** Survival of nematodes upon treatment with various concentrations of **1e**, **8c**, and **8d**.

| | Mean lifespan ± SE (days) | Extention rate (%) | N | p-value |
|---|---|---|---|---|
| DMSO 0.1% | 17.0 ± 3.4 | | 207 | |
| Concentration | **1e** | | | |
| 1 μg/ml | 20.5 ± 3.2 | 20.5 | 104 | 0.0399 |
| 10 μg/ml | 21.4 ± 2.2 | 25.6 | 123 | 0.0252 |
| 50 μg/ml | 21.0 ± 2.0 | 23.6 | 89 | 0.0269 |
| 100 μg/ml | 21.2 ± 1.9 | 24.7 | 94 | 0.0141 |
| 250 μg/ml | 21.2 ± 1.4 | 24.5 | 76 | 0.0227 |
| 500 μg/ml | 21.5 ± 1.8 | 26.4 | 123 | 0.0064 |
| Concentration | **8c** | | | |
| 1 μg/ml | 19.1 ± 1.9 | 12.2 | 112 | 0.4151 |
| 10 μg/ml | 20.8 ± 1.0 | 22.3 | 97 | 0.0256 |
| 50 μg/ml | 22.5 ± 2.3 | 32.1 | 102 | 0.003 |
| 100 μg/ml | 22.0 ± 1.9 | 29.3 | 107 | 0.0039 |
| 250 μg/ml | 19.1 ± 1.4 | 12.2 | 101 | 0.2997 |
| 500 μg/ml | 16.5 ± 0.9 | -2.7 | 105 | 0.374 |
| Concentration | **8d** | | | |
| 1 μg/ml | 21.0 ± 3.4 | 23.7 | 82 | 0.0337 |
| 10 μg/ml | 20.8 ± 0.9 | 22.6 | 81 | 0.0292 |
| 50 μg/ml | 20.7 ± 1.8 | 21.6 | 98 | 0.0381 |
| 100 μg/ml | 22.1 ± 1.6 | 30.2 | 101 | 0.0013 |
| 250 μg/ml | 19.8 ± 2.6 | 16.4 | 102 | 0.1906 |
| 500 μg/ml | 19.0 ± 2.1 | 11.8 | 96 | 0.4011 |

### NMR spectra

$^{1}$H NMR spectrum of **6a**

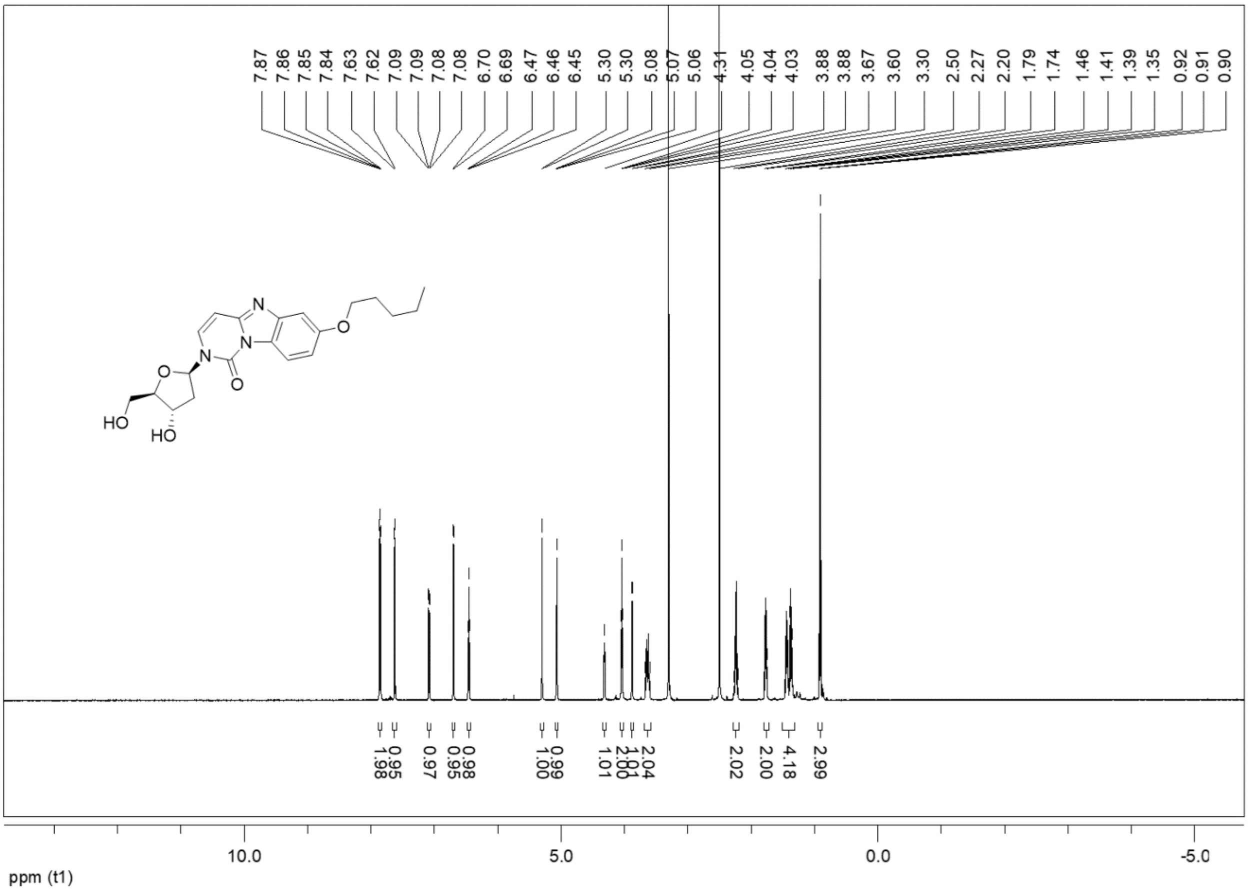

$^{13}$C NMR spectrum of **6a**

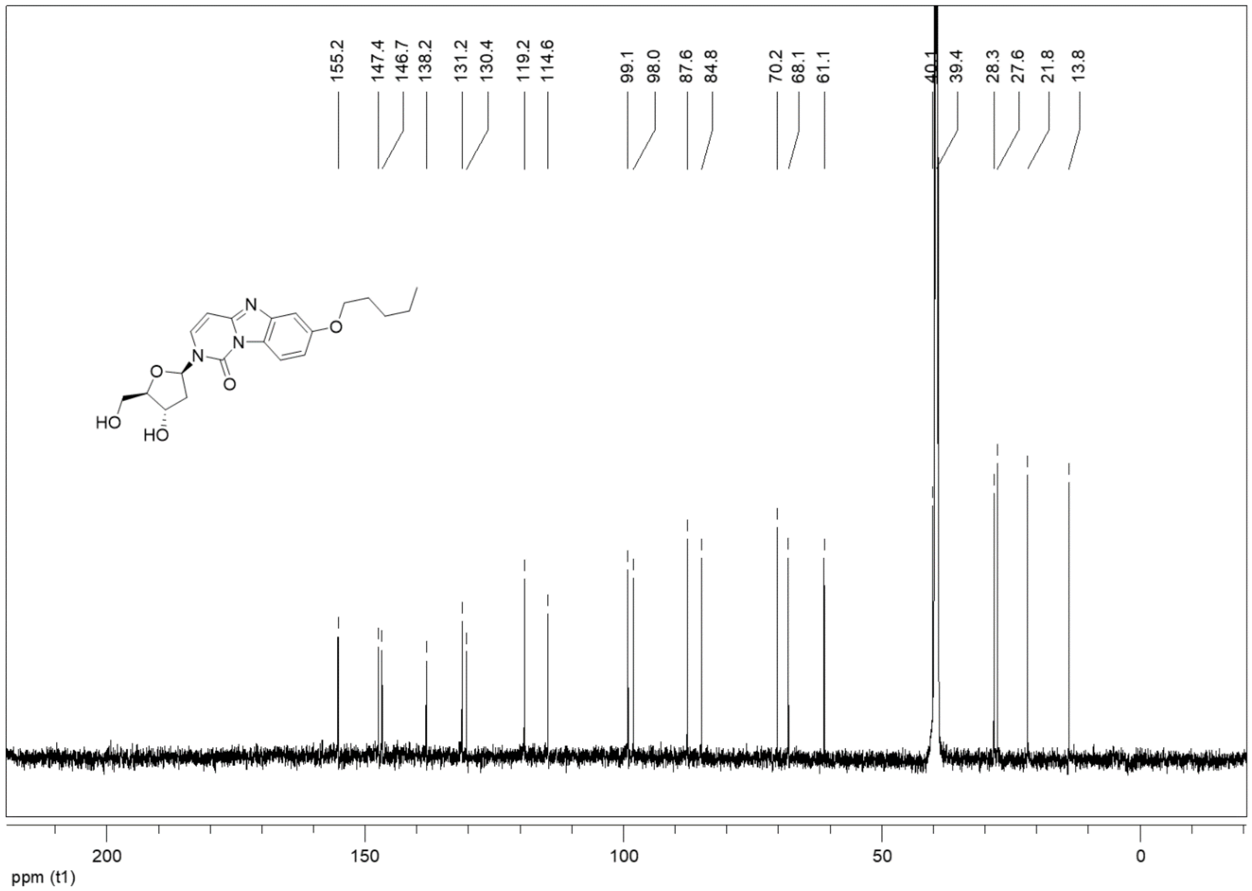

$^1$H NMR spectrum of **6b**

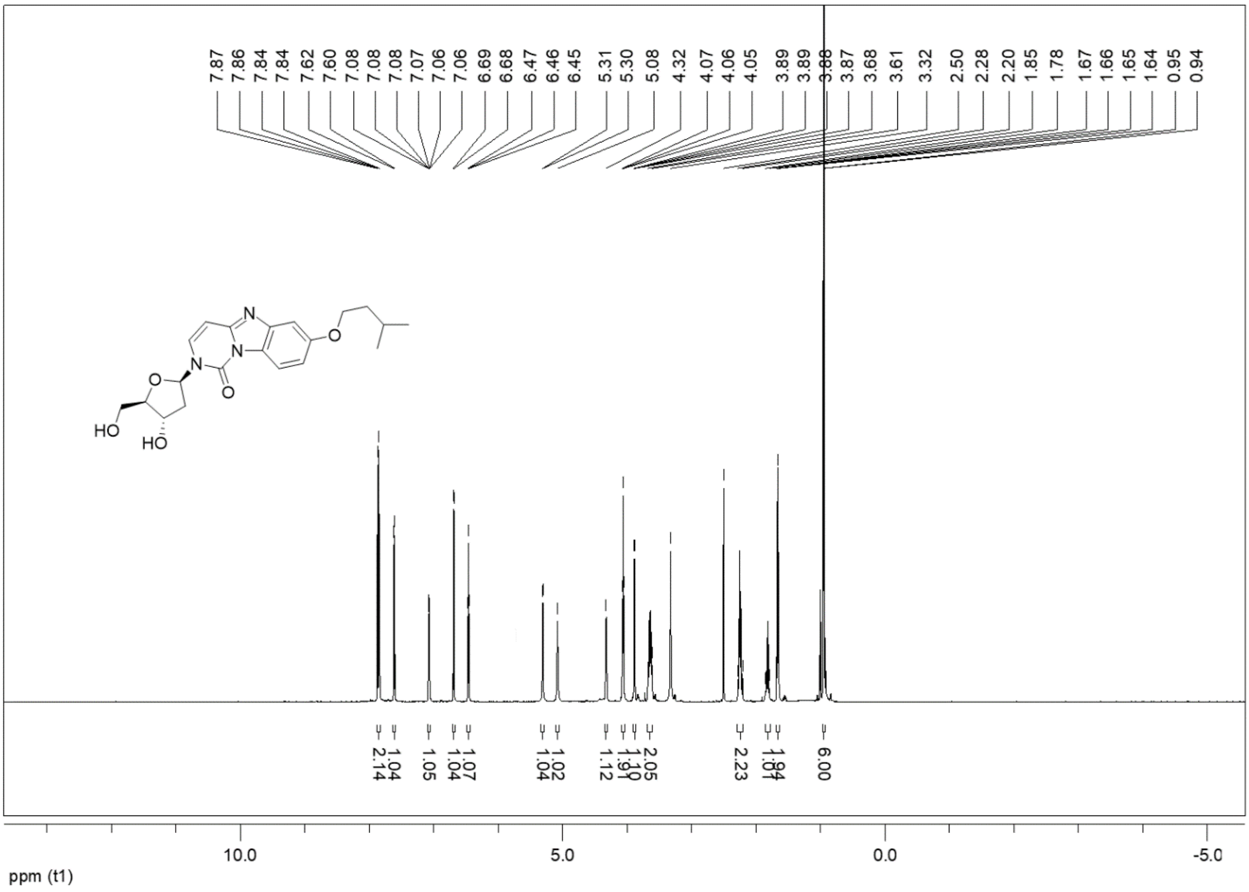

$^{13}$C NMR spectrum of **6b**

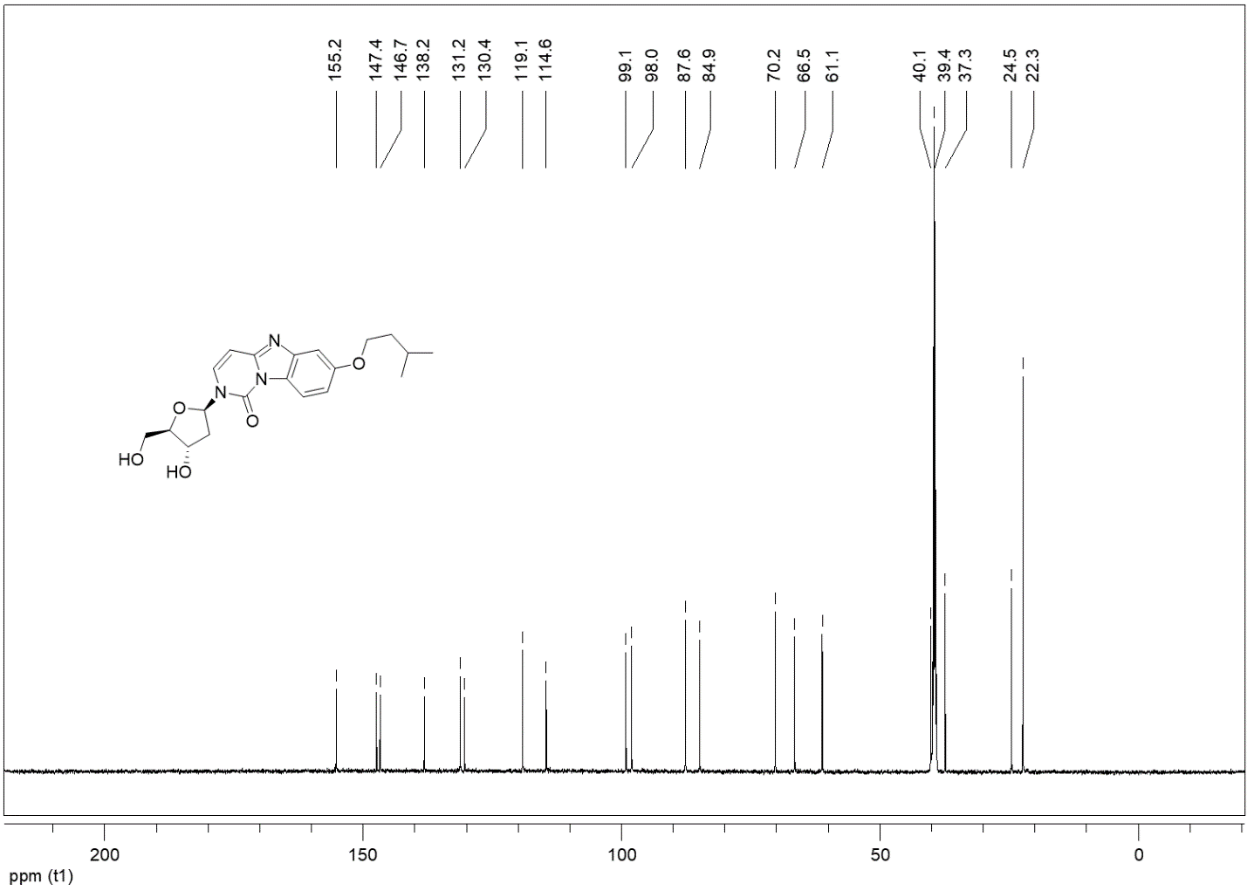

$^{1}$H NMR spectrum of **6c**

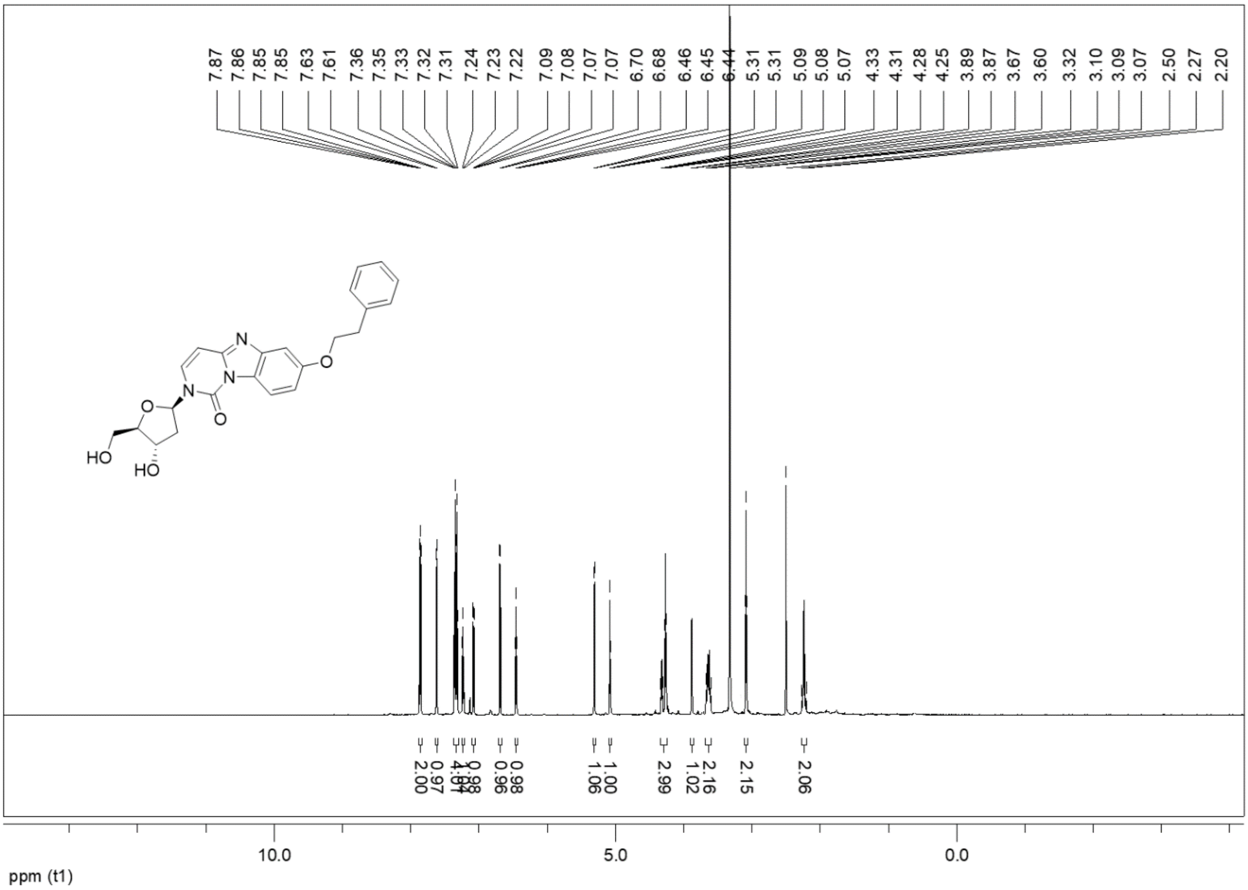

<sup></sup>$^{13}$C NMR spectrum of **6c**

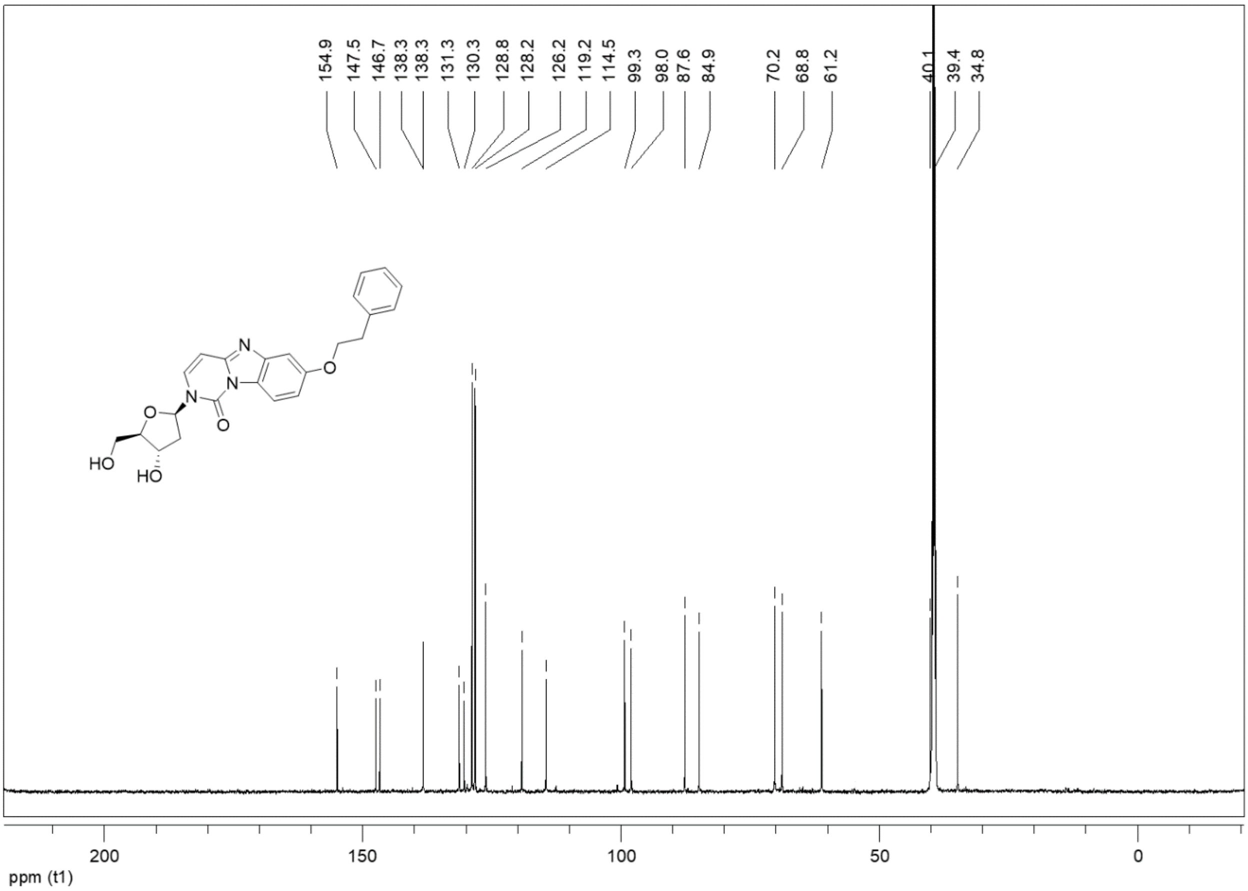

[1]H NMR spectrum of **6d**

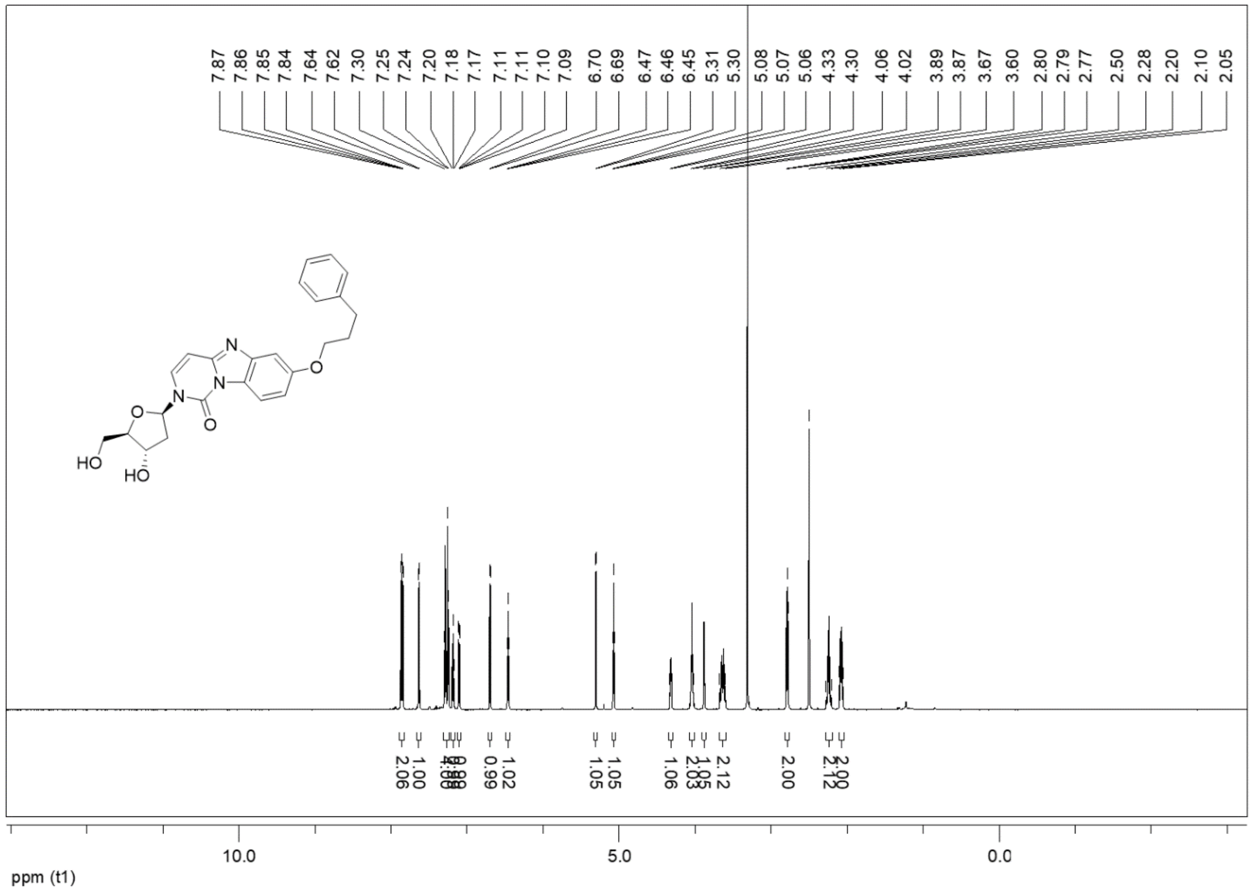

$^{13}C$ NMR spectrum of **6d**

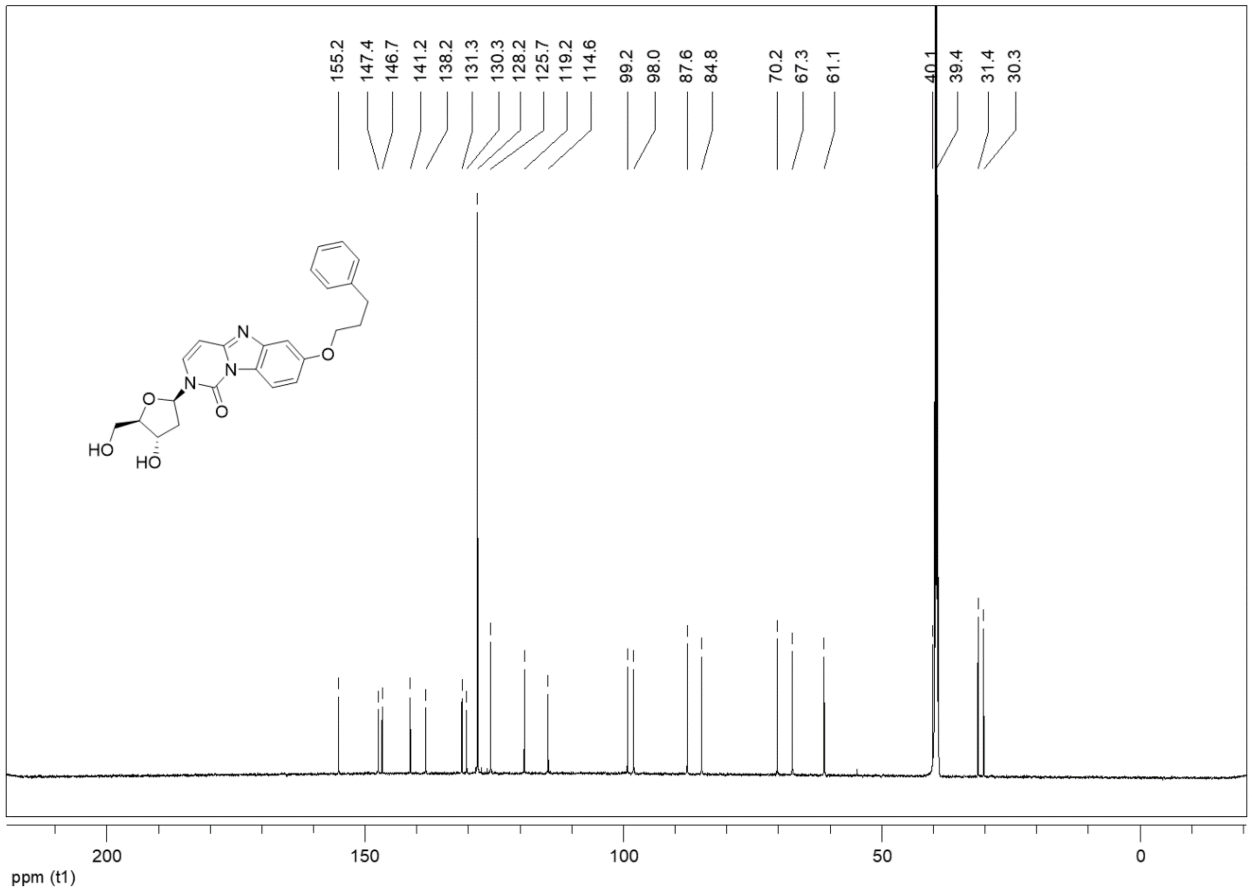

[1]H NMR spectrum of **6e**

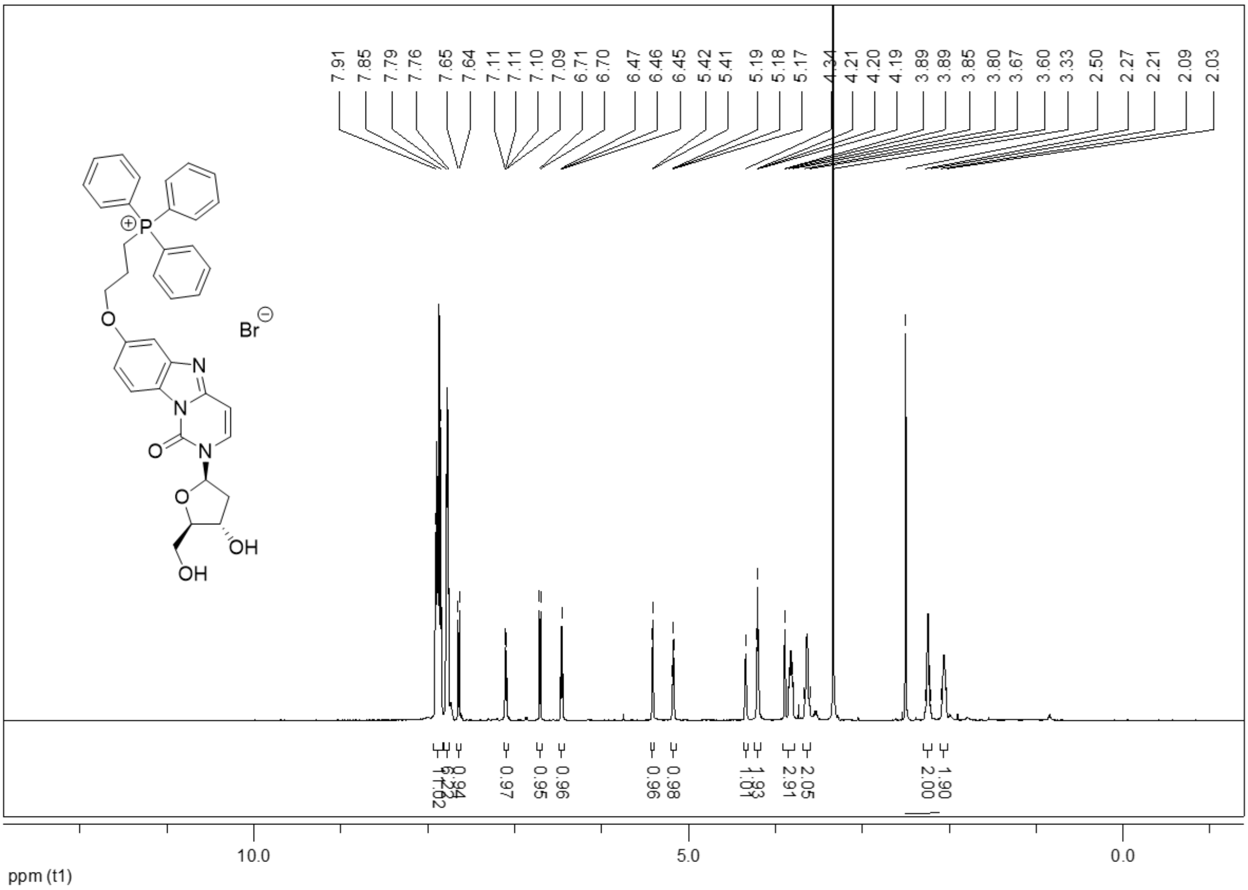

$^{13}$C NMR spectrum of **6e**

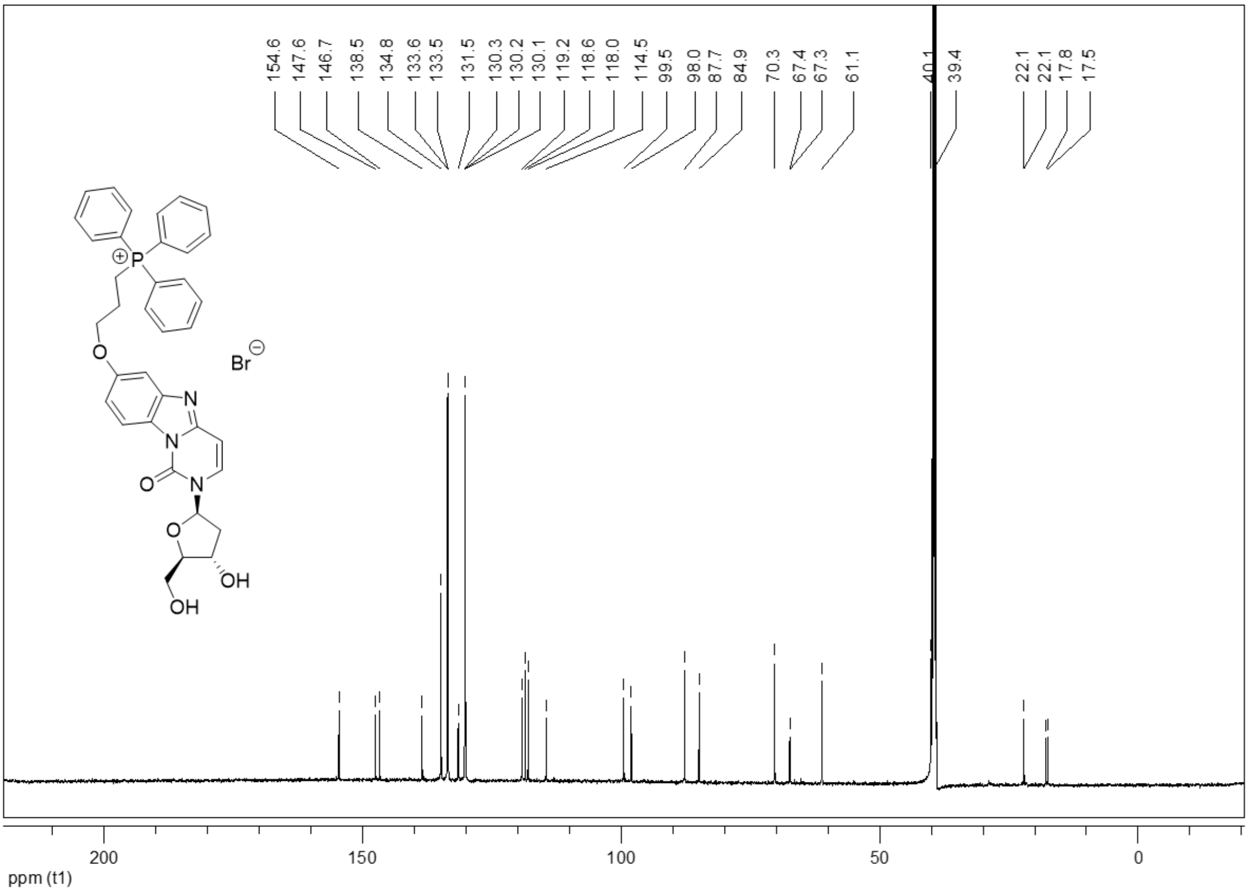

$^{31}$P NMR spectrum of **6e**

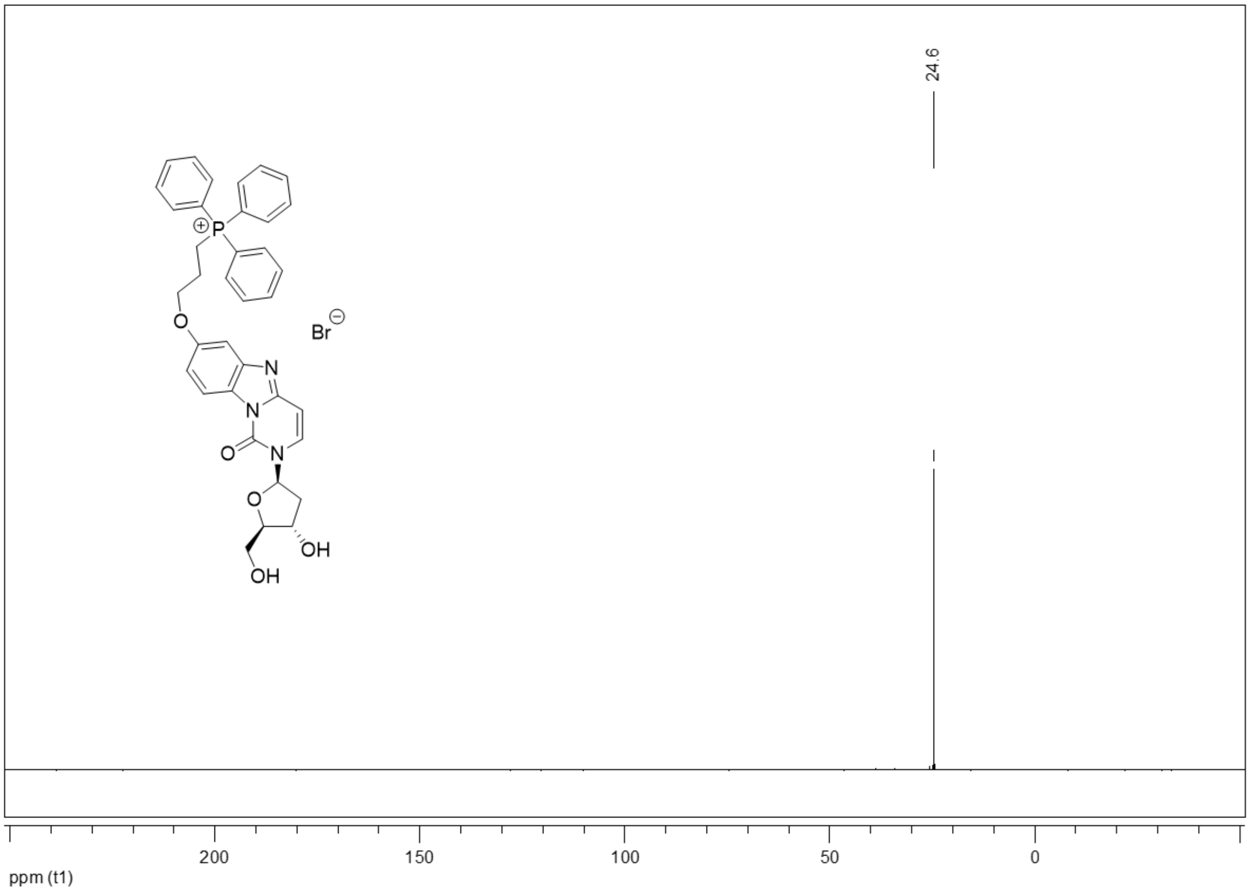

[1]H NMR spectrum of **6f**

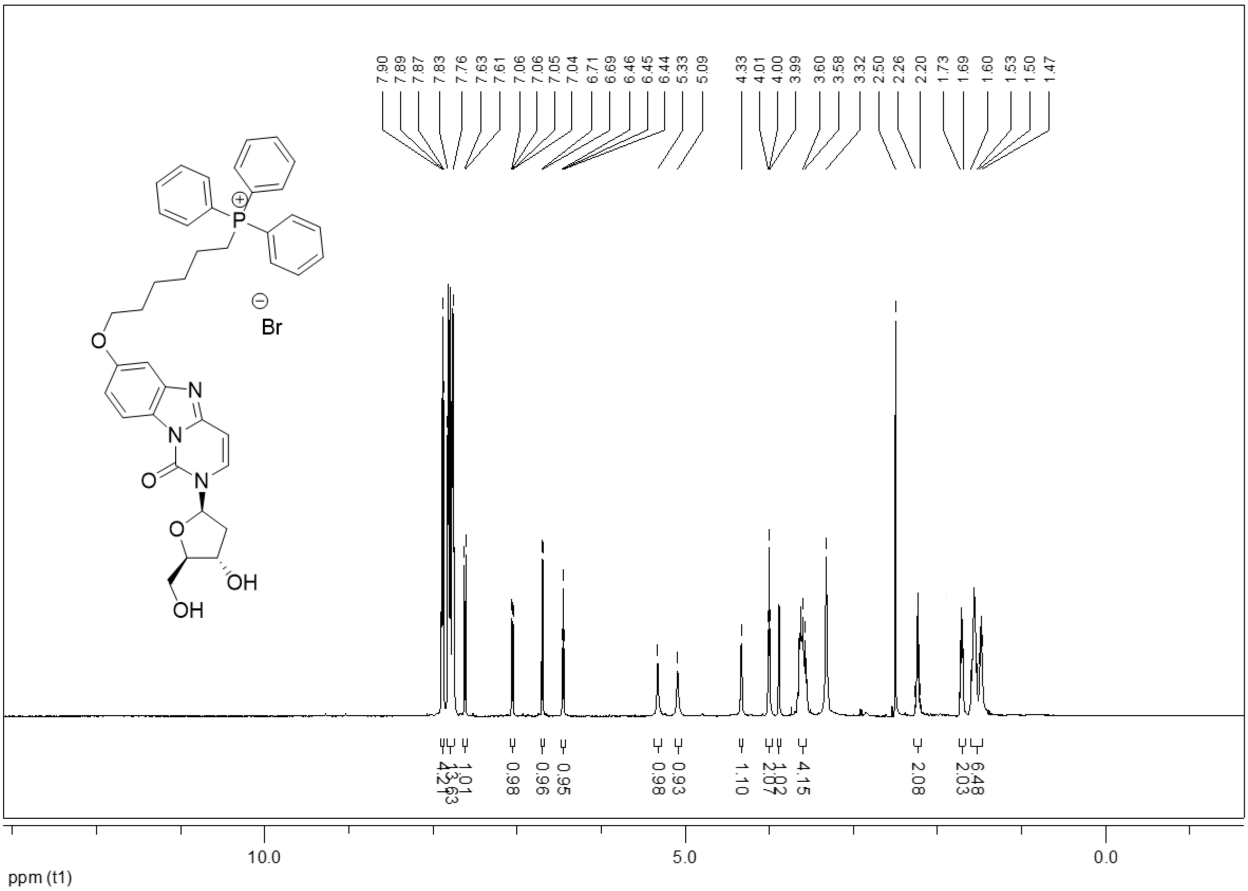

$^{13}C$ NMR spectrum of **6f**

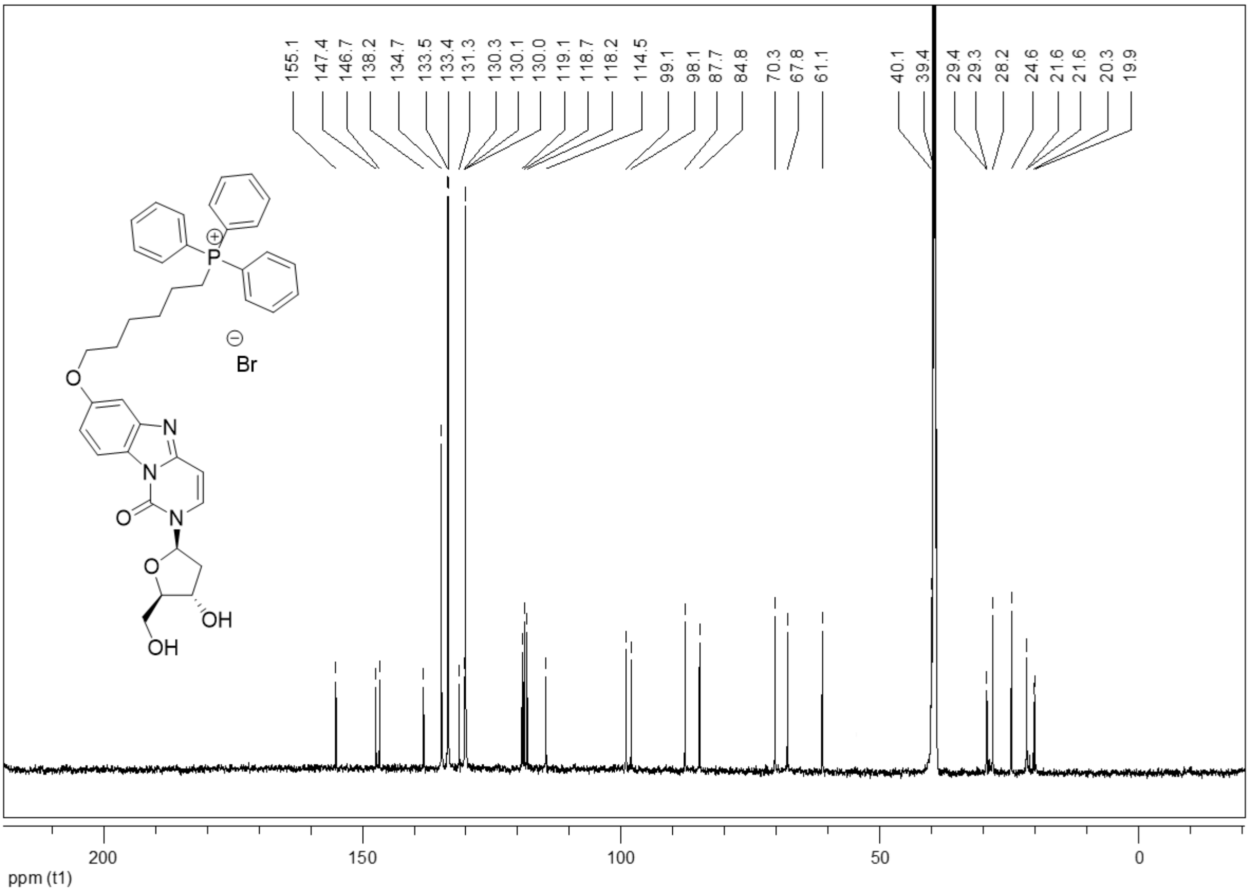

$^{31}$P NMR spectrum of **6f**

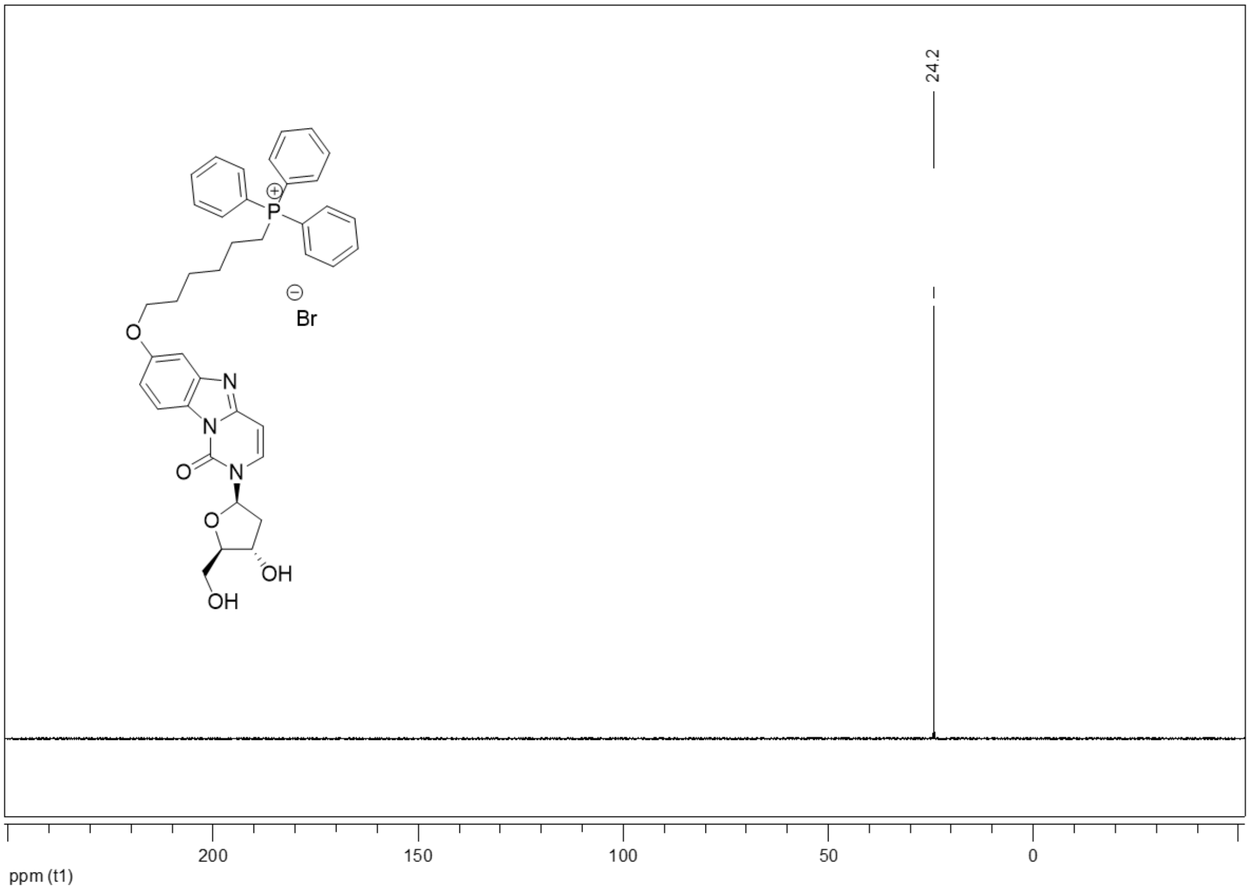

[1]H NMR spectrum of **6g**

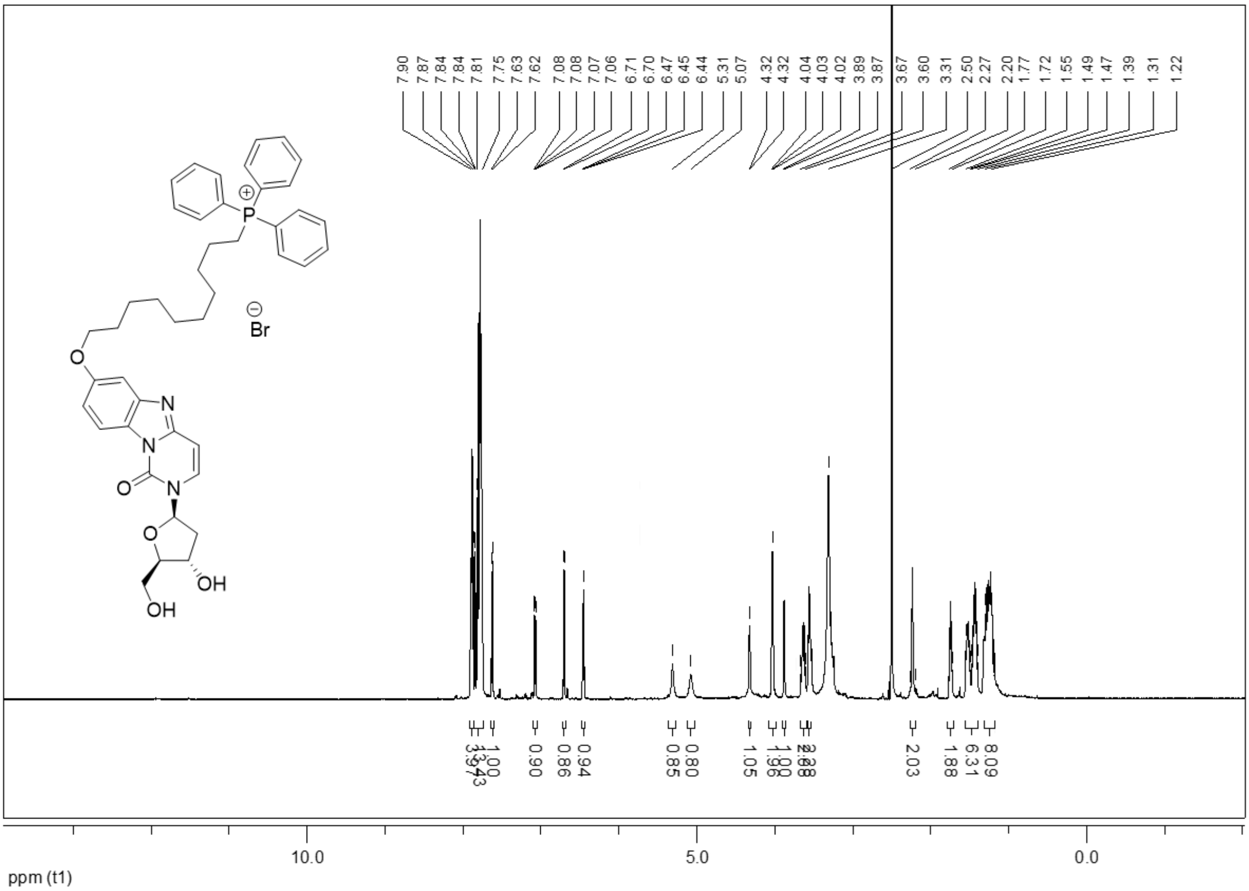

[13]C NMR spectrum of **6g**

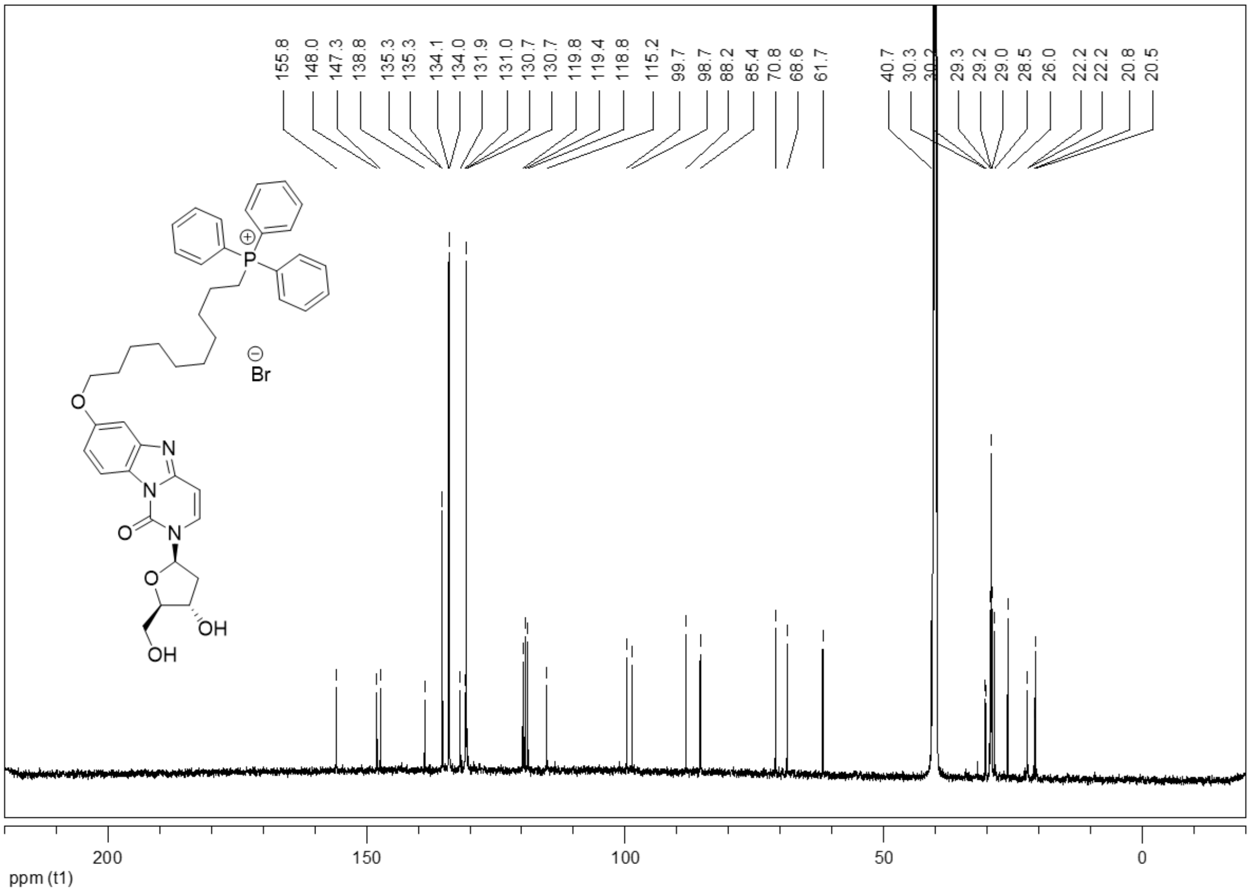

[31]P NMR spectrum of **6g**

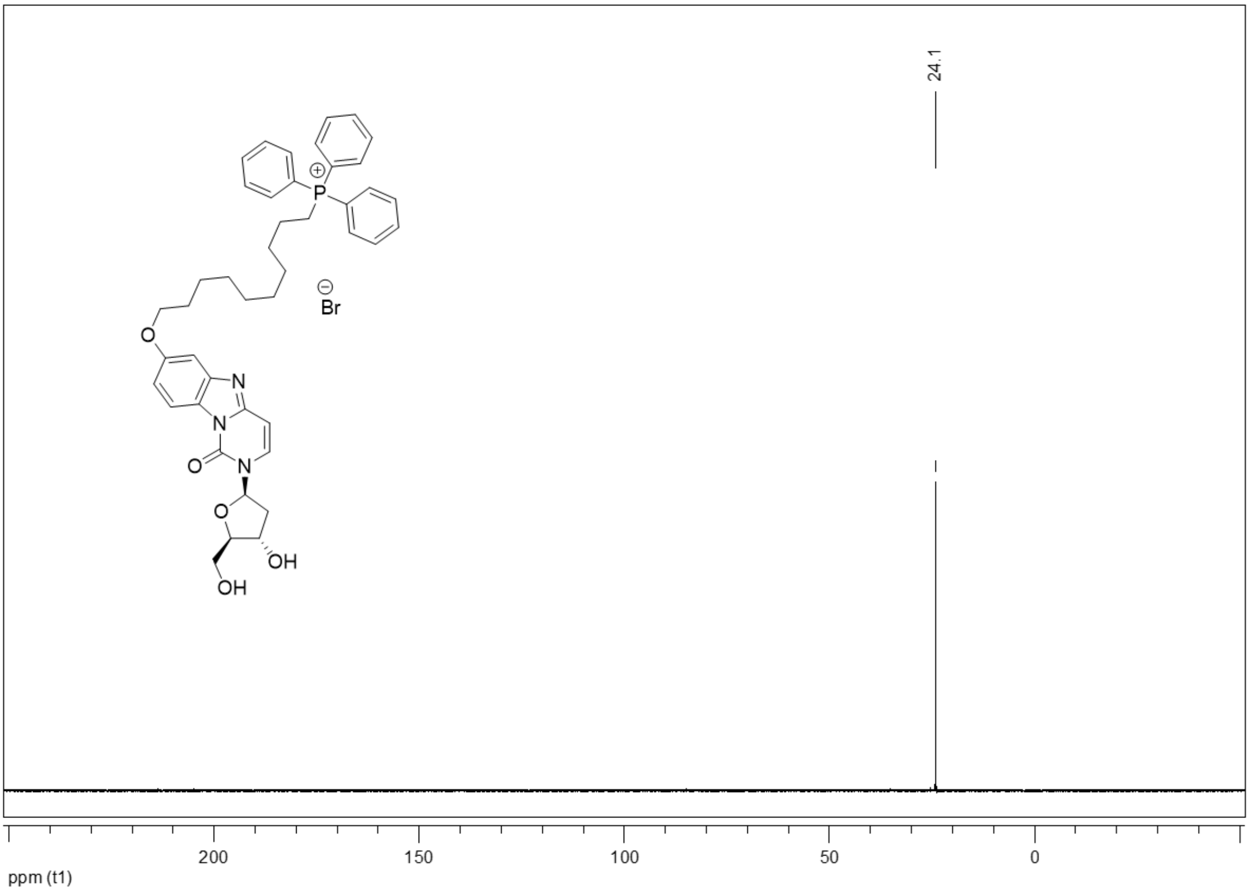

<sup></sup>

[1]H NMR spectrum of **7a**

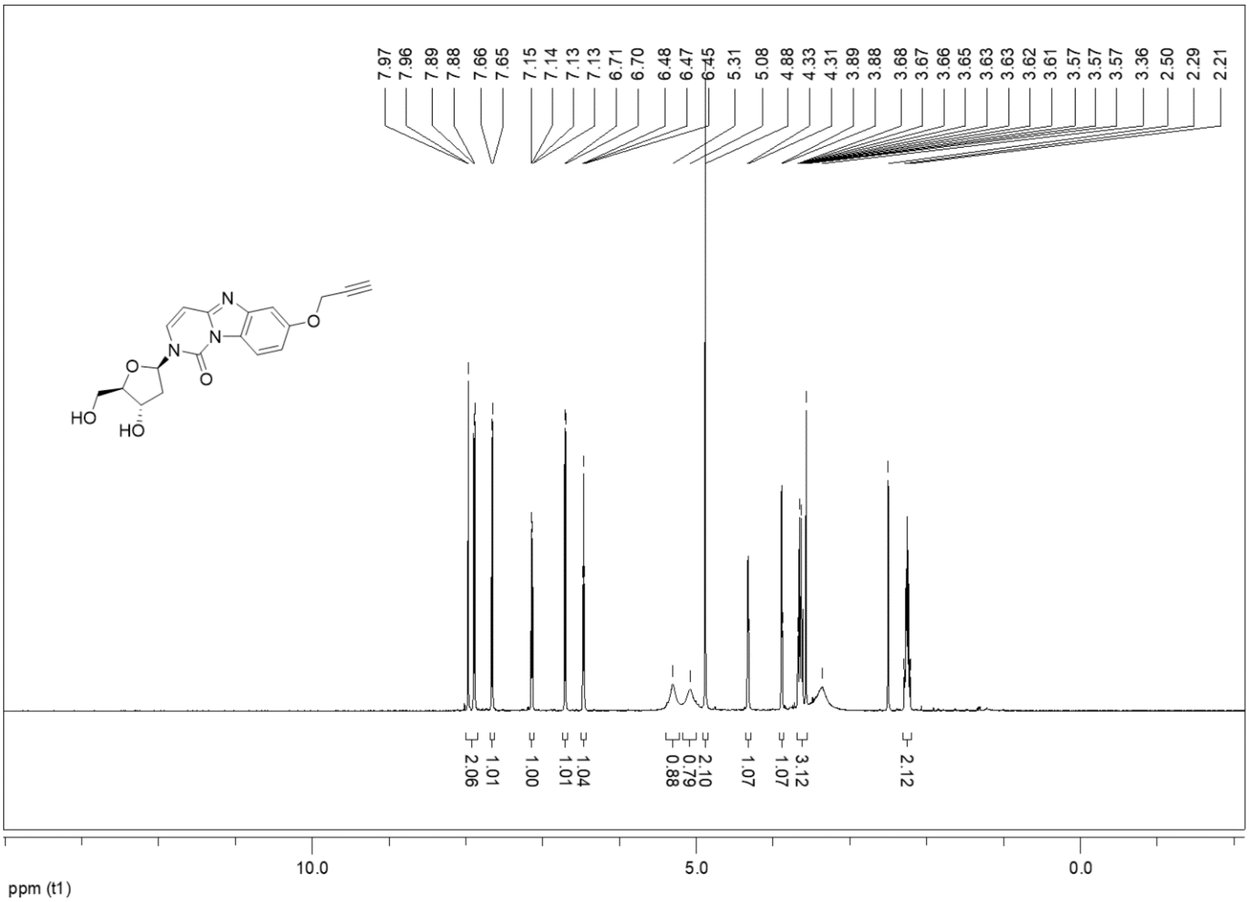

$^{13}$C NMR spectrum of **7a**

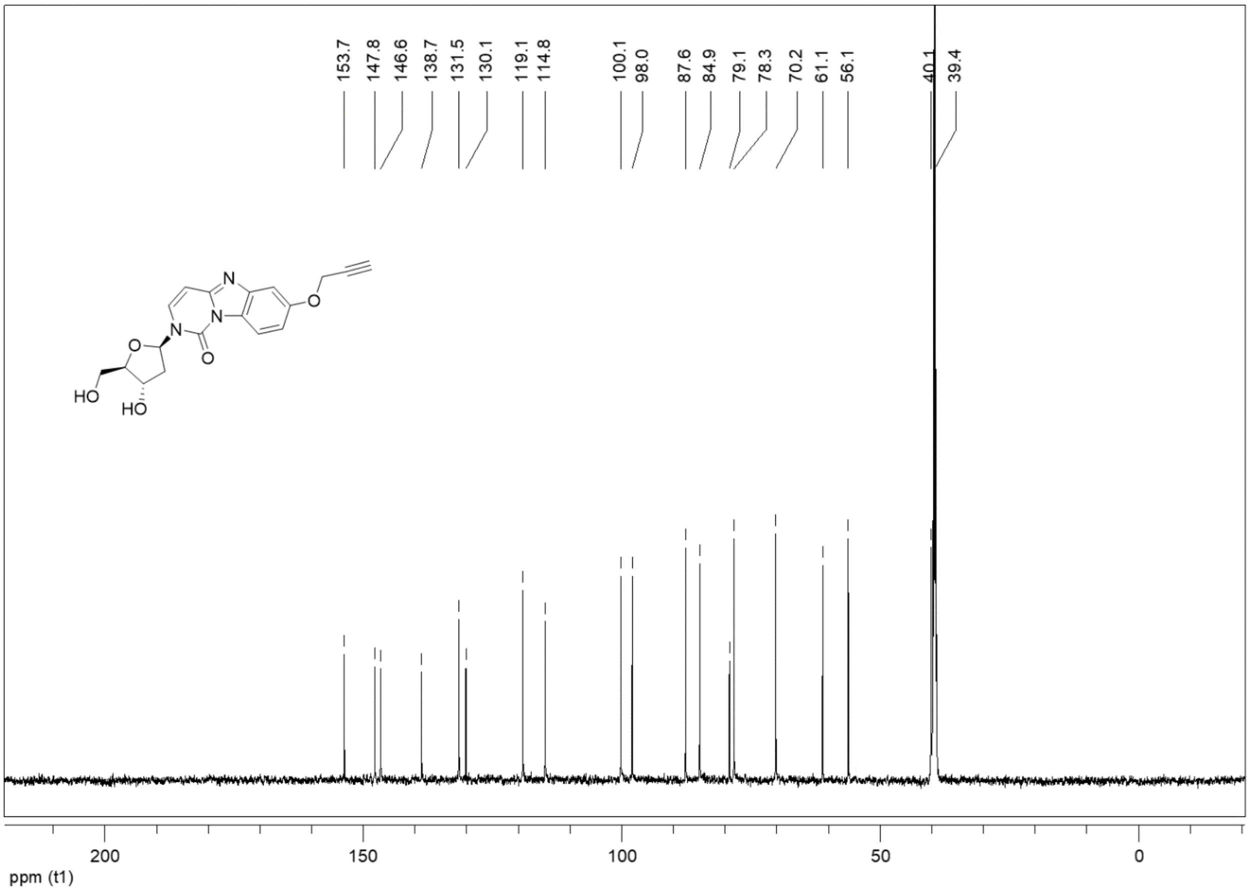

[1]H NMR spectrum of **7b**

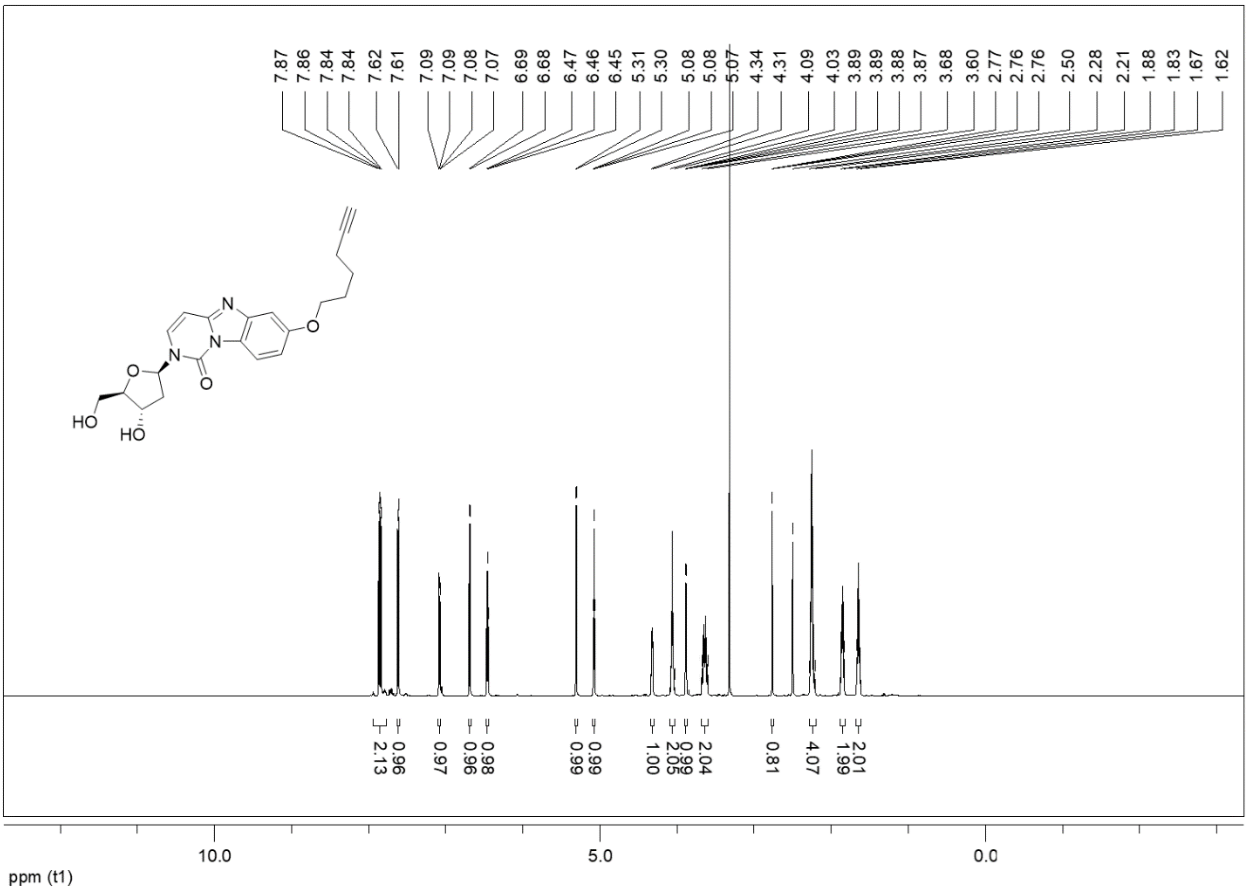

$^{13}$C NMR spectrum of **7b**

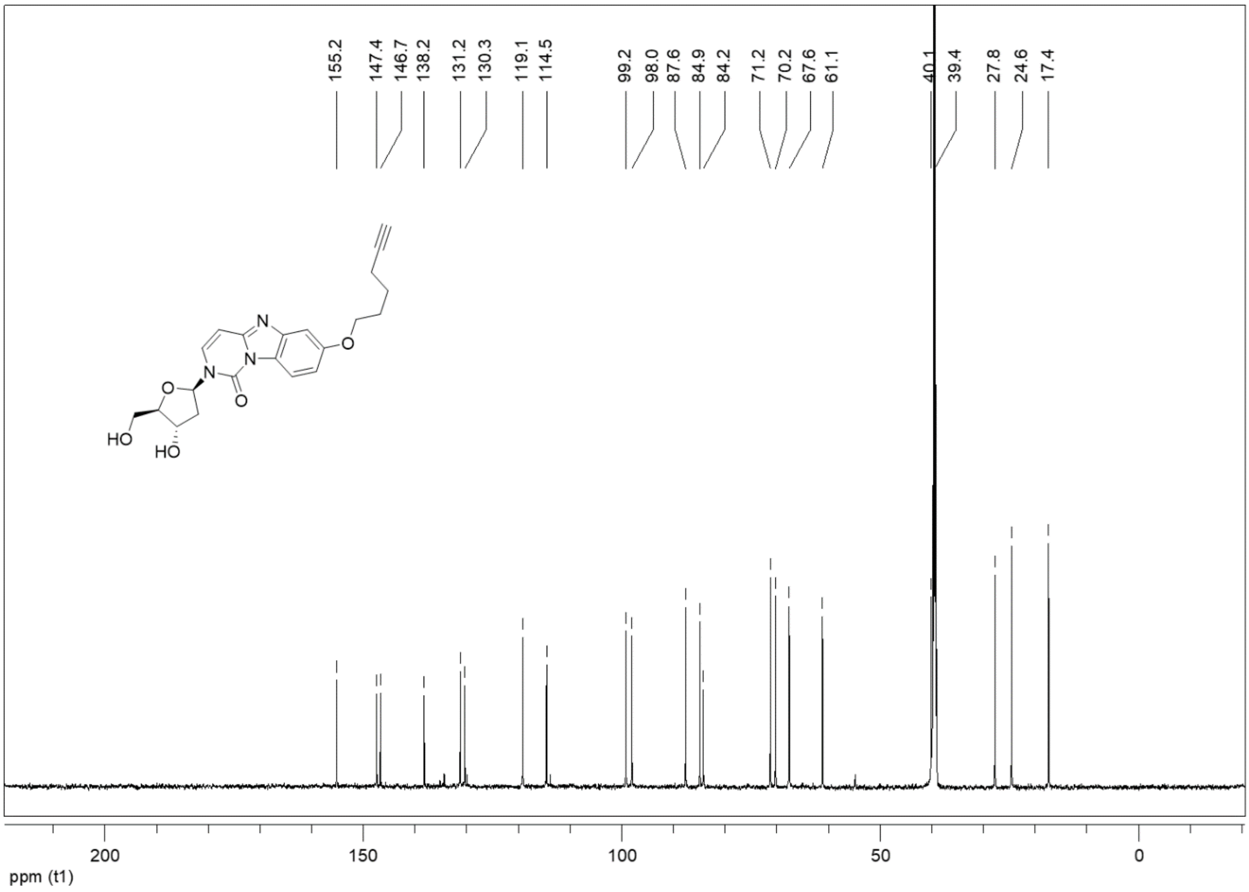

$^{1}H$ NMR spectrum of **8a**

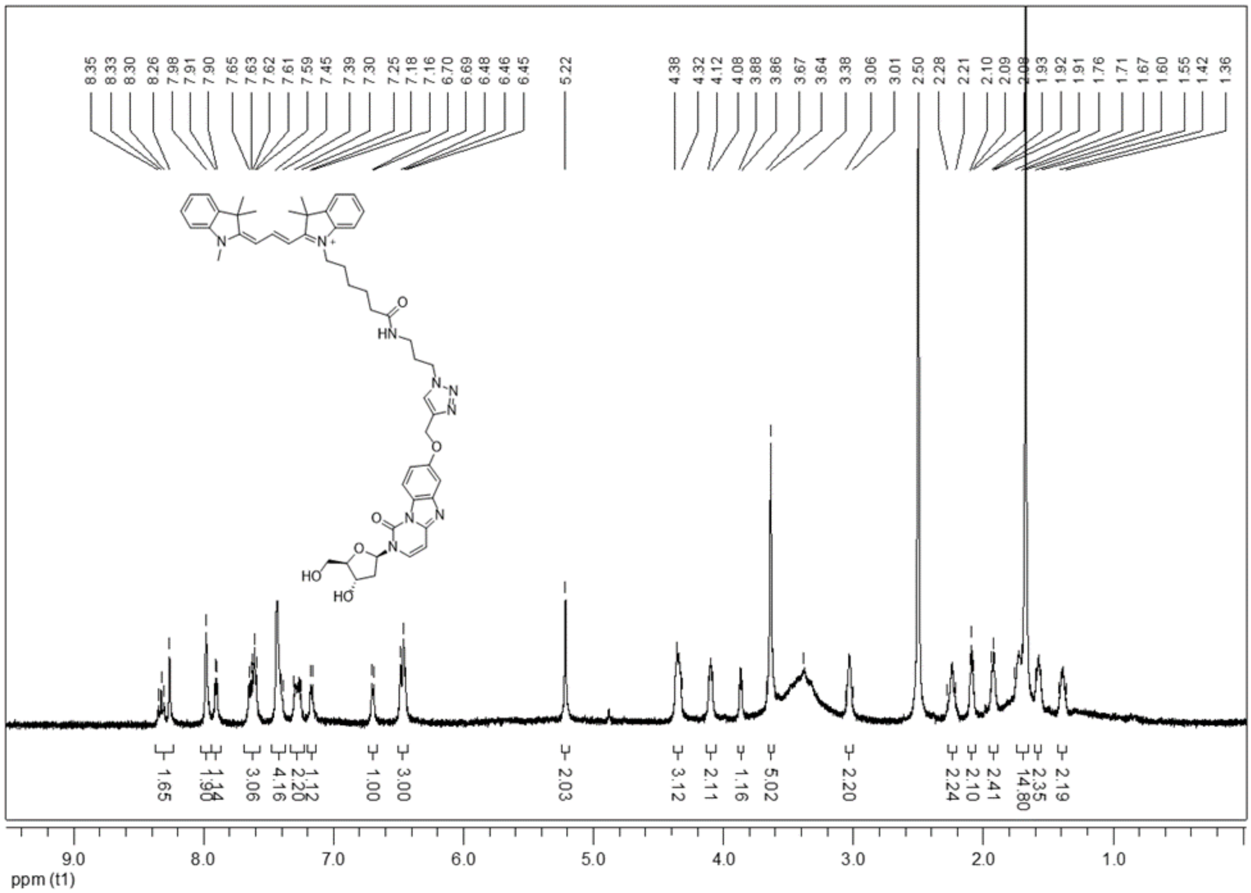

<sup></sup>[1]H NMR spectrum of **8b**

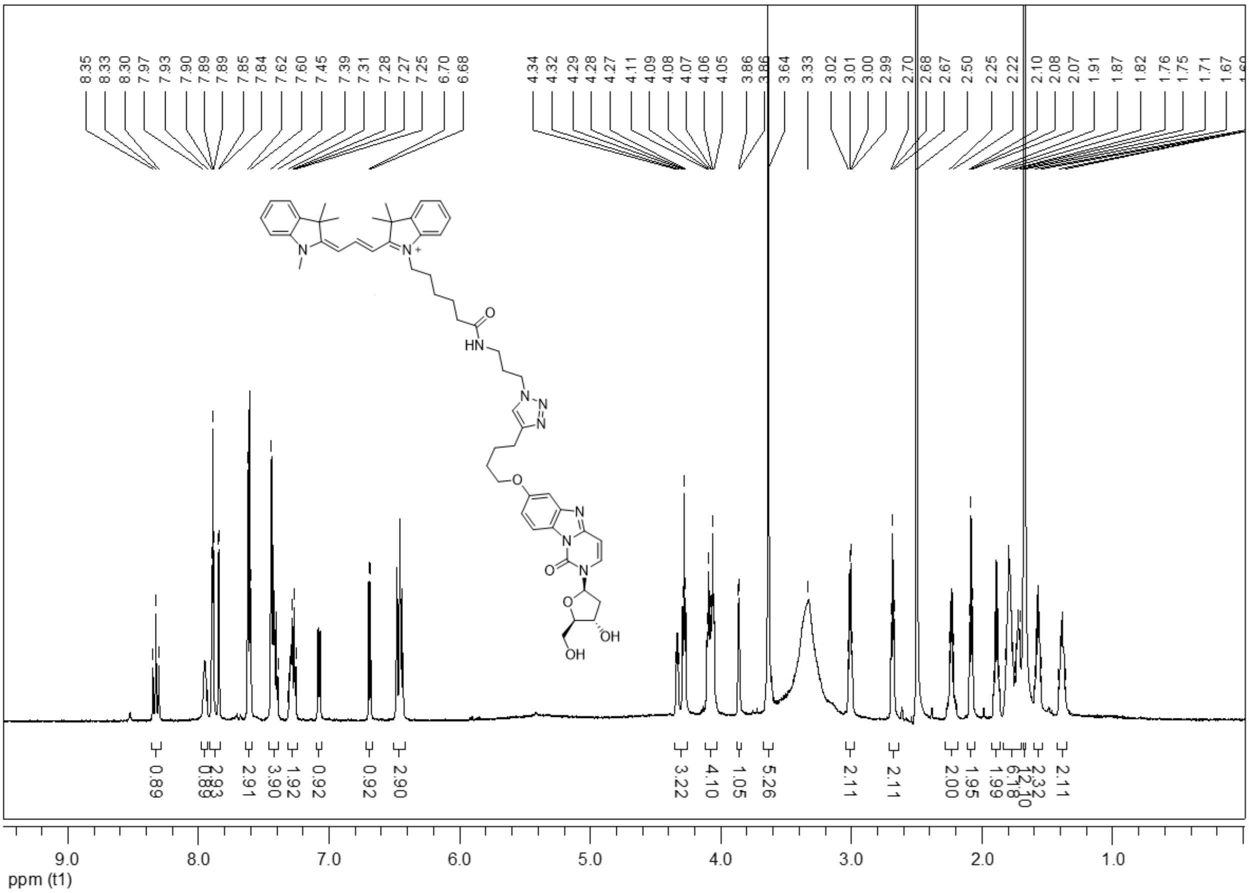

¹H NMR spectrum of **8c**

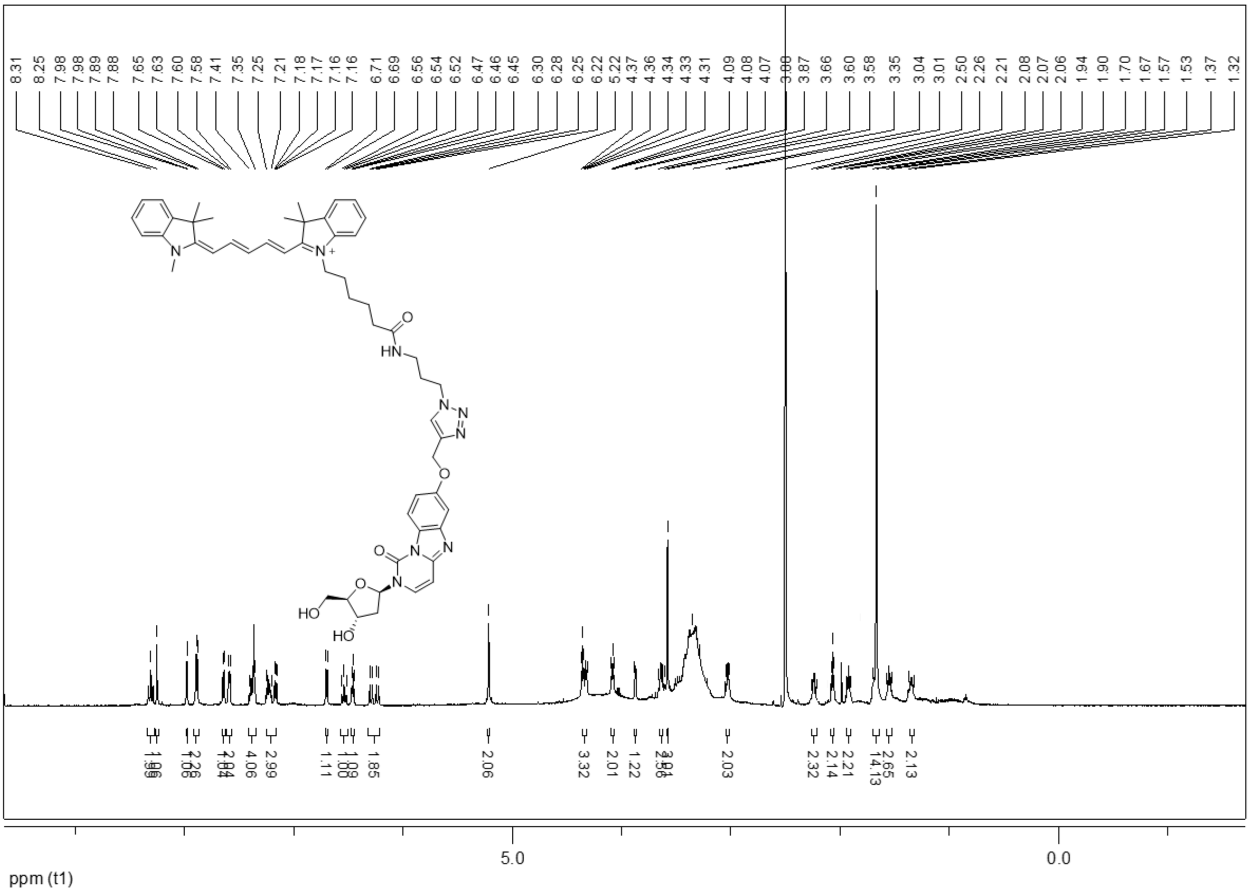

$^{1}H$ NMR spectrum of **8d**

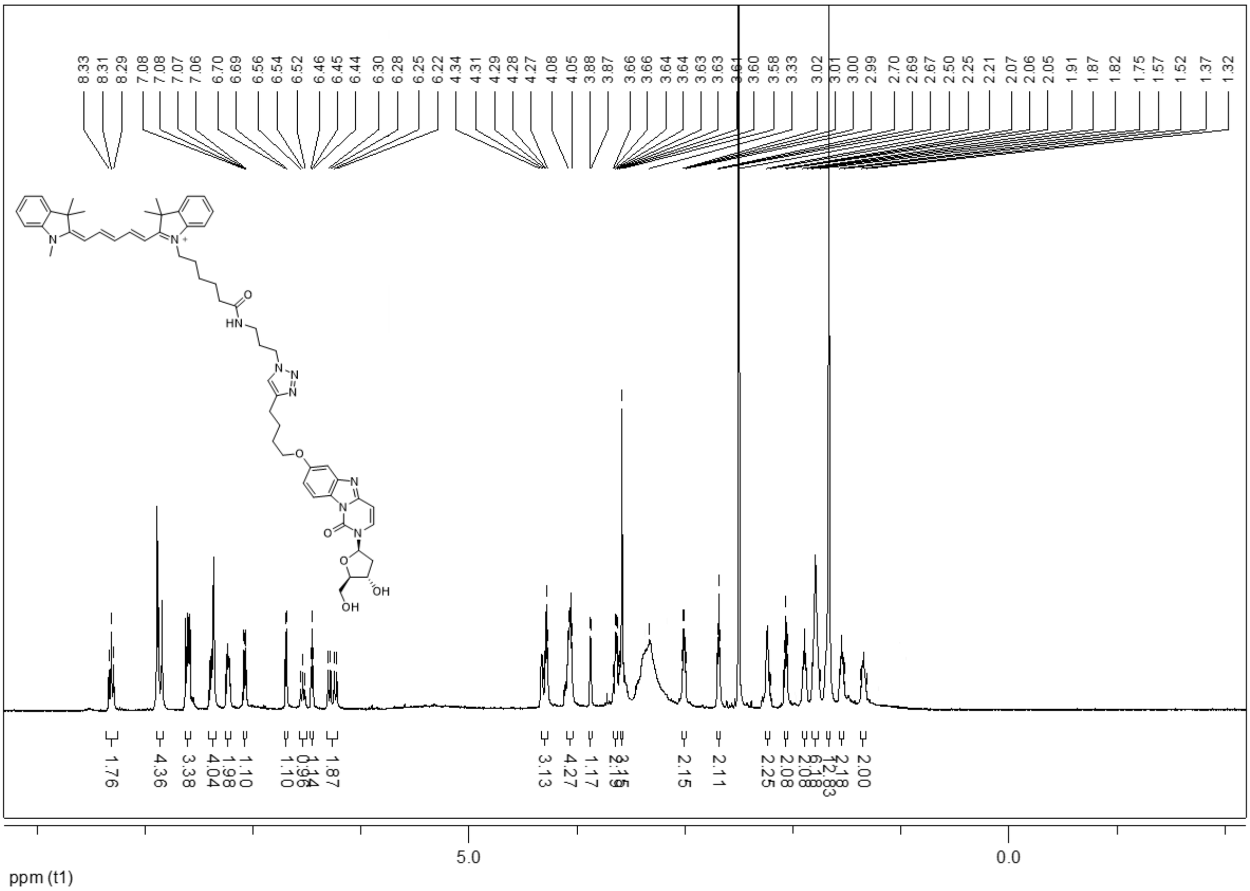

### HPLC data

*2-((2R,4S,5R)-4-hydroxy-5-(hydroxymethyl)tetrahydrofuran-2-yl)-7-(pentyloxy)benzo[4,5]imidazo[1,2-c]pyrimidin-1(2H)-one* **6a**

**283 AR975**

| | | | |
|---|---|---|---|
| *Sample Name:* | **AR975** | *Injection Volume:* | **20.0** |
| *Vial Number:* | **RG9** | *Channel:* | **UV_VIS_1** |
| *Sample Type:* | **unknown** | *Wavelength:* | **210.0** |
| *Control Program:* | **NewObj_short** | *Bandwidth:* | **4** |
| *Quantif. Method:* | **default** | *Dilution Factor:* | **1.0000** |
| *Recording Time:* | **20.9.2025 0:34** | *Sample Weight:* | **1.0000** |
| *Run Time (min):* | **8.00** | *Sample Amount:* | **1.0000** |

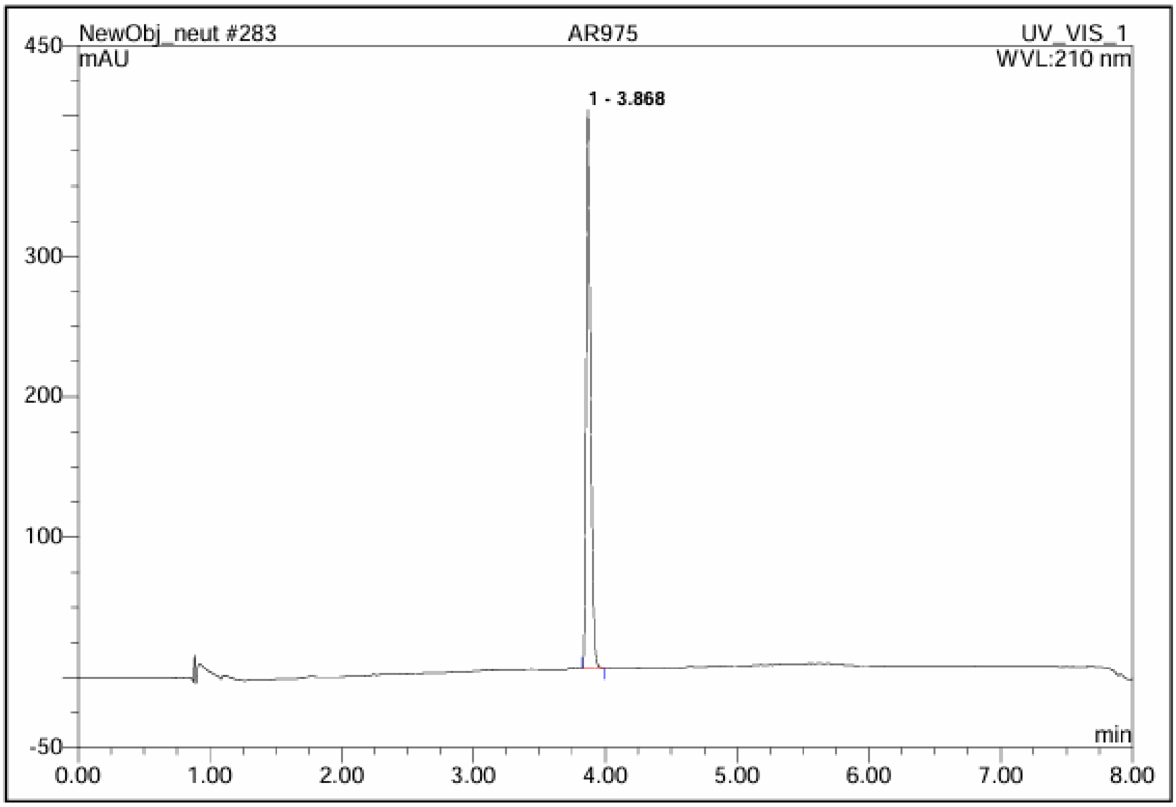

| No. | Ret.Time min | Peak Name | Height mAU | Area mAU*min | Rel.Area % | Amount | Type |
|---|---|---|---|---|---|---|---|
| 1 | 3.87 | n.a. | 397.830 | 15.081 | 100.00 | n.a. | BMB |
| **Total:** | | | 397.830 | 15.081 | 100.00 | 0.000 | |

*2-((2R,4S,5R)-4-hydroxy-5-(hydroxymethyl)tetrahydrofuran-2-yl)-7-(isopentyloxy)benzo[4,5]imidazo[1,2-c]pyrimidin-1(2H)-one* **6b**

**287 AR974**

| | | | |
|---|---|---|---|
| *Sample Name:* | **AR974** | *Injection Volume:* | **20.0** |
| *Vial Number:* | **RH6** | *Channel:* | **UV_VIS_1** |
| *Sample Type:* | **unknown** | *Wavelength:* | **210.0** |
| *Control Program:* | **NewObj_short** | *Bandwidth:* | **4** |
| *Quantif. Method:* | **default** | *Dilution Factor:* | **1.0000** |
| *Recording Time:* | **20.9.2025 1:23** | *Sample Weight:* | **1.0000** |
| *Run Time (min):* | **8.00** | *Sample Amount:* | **1.0000** |

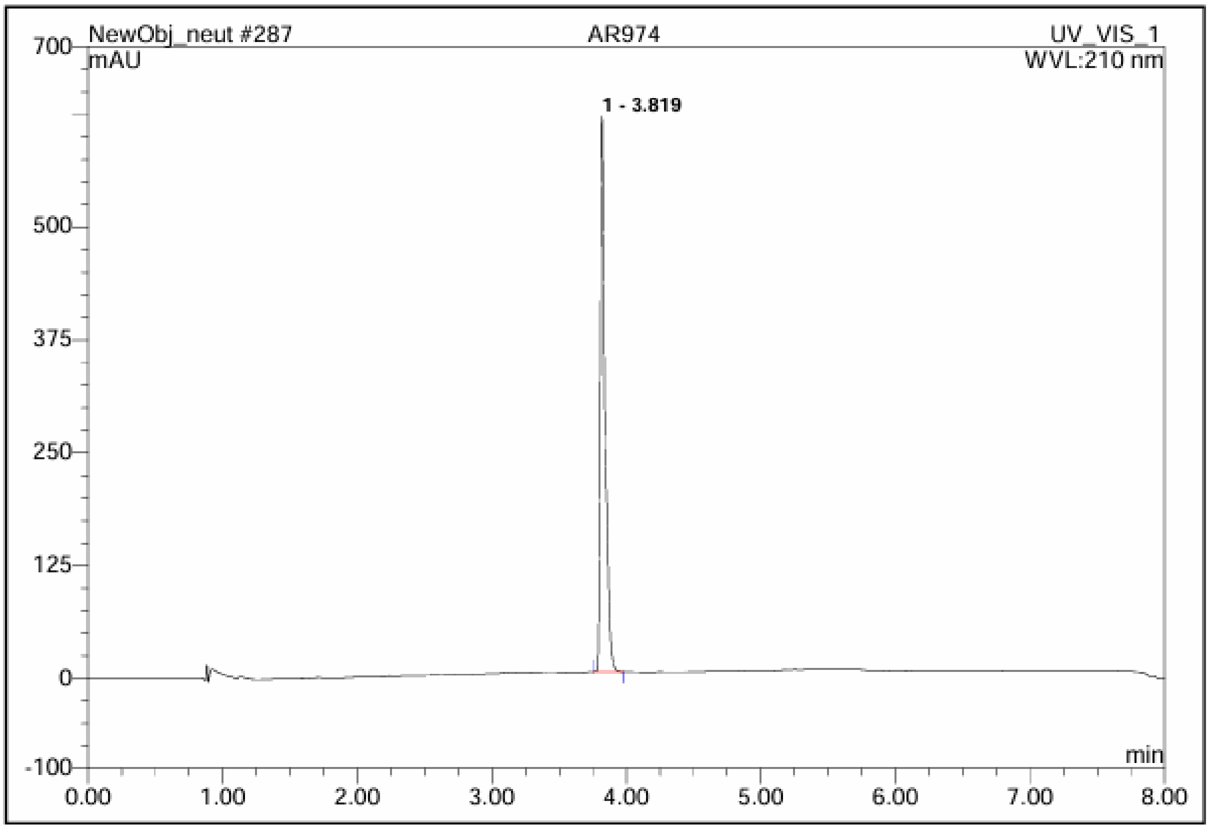

| No. | Ret.Time min | Peak Name | Height mAU | Area mAU*min | Rel.Area % | Amount | Type |
|---|---|---|---|---|---|---|---|
| 1 | 3.82 | n.a. | 615.340 | 24.924 | 100.00 | n.a. | BMB |
| **Total:** | | | 615.340 | 24.924 | 100.00 | 0.000 | |

*2-((2R,4S,5R)-4-hydroxy-5-(hydroxymethyl)tetrahydrofuran-2-yl)-7-phenethoxybenzo[4,5]imidazo[1,2-c]pyrimidin-1(2H)-one* **6c**

## 285 AR971

| | | | |
|---|---|---|---|
| *Sample Name:* | **AR971** | *Injection Volume:* | **20.0** |
| *Vial Number:* | **RC7** | *Channel:* | **UV_VIS_1** |
| *Sample Type:* | **unknown** | *Wavelength:* | **210.0** |
| *Control Program:* | **NewObj_short** | *Bandwidth:* | **4** |
| *Quantif. Method:* | **default** | *Dilution Factor:* | **1.0000** |
| *Recording Time:* | **20.9.2025 0:59** | *Sample Weight:* | **1.0000** |
| *Run Time (min):* | **8.00** | *Sample Amount:* | **1.0000** |

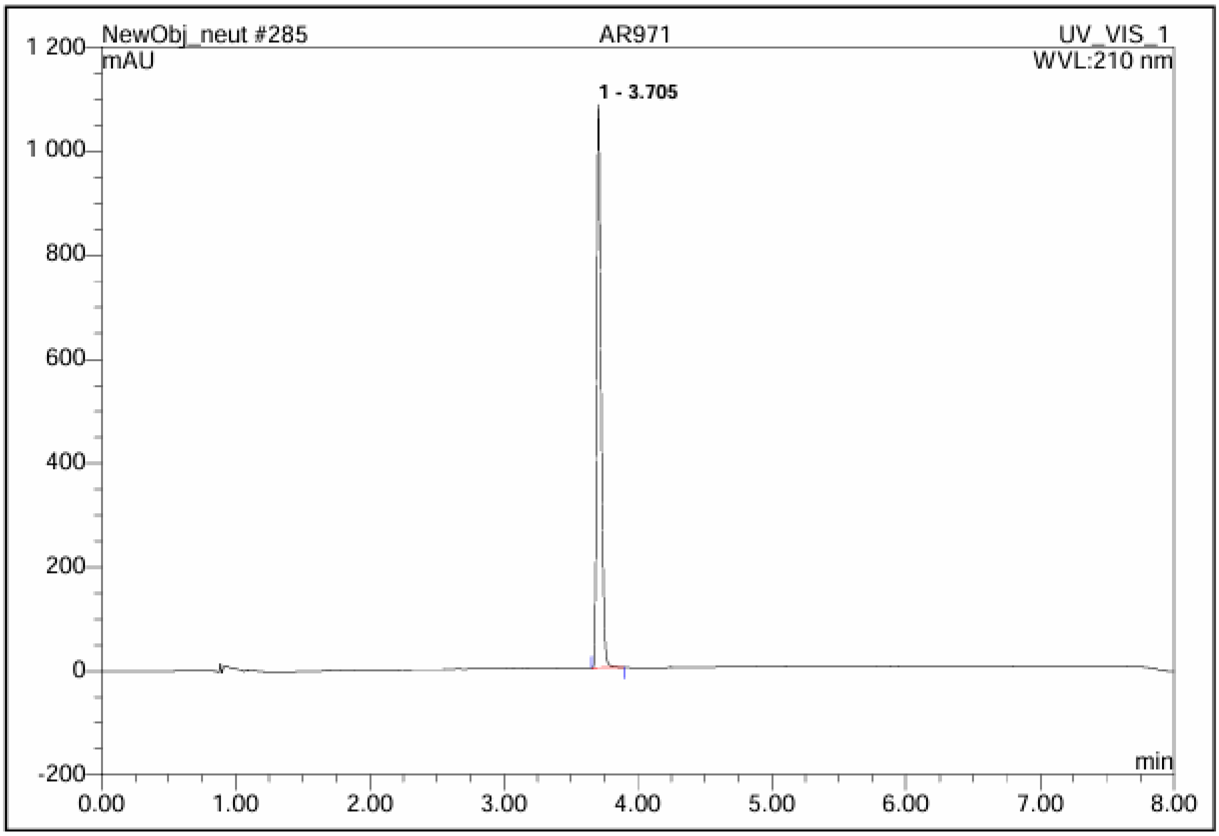

| No. | Ret.Time min | Peak Name | Height mAU | Area mAU*min | Rel.Area % | Amount | Type |
|---|---|---|---|---|---|---|---|
| 1 | 3.71 | n.a. | 1082.852 | 40.016 | 100.00 | n.a. | BMB |
| **Total:** | | | 1082.852 | 40.016 | 100.00 | 0.000 | |

*2-((2R,4S,5R)-4-hydroxy-5-(hydroxymethyl)tetrahydrofuran-2-yl)-7-(3-phenylpropoxy)benzo[4,5]imidazo[1,2-c]pyrimidin-1(2H)-one* **6d**

**280 AR972**

| | | | |
|---|---|---|---|
| *Sample Name:* | **AR972** | *Injection Volume:* | **20.0** |
| *Vial Number:* | **RD5** | *Channel:* | **UV_VIS_1** |
| *Sample Type:* | **unknown** | *Wavelength:* | **210.0** |
| *Control Program:* | **NewObj_short** | *Bandwidth:* | **4** |
| *Quantif. Method:* | **default** | *Dilution Factor:* | **1.0000** |
| *Recording Time:* | **19.9.2025 22:59** | *Sample Weight:* | **1.0000** |
| *Run Time (min):* | **8.00** | *Sample Amount:* | **1.0000** |

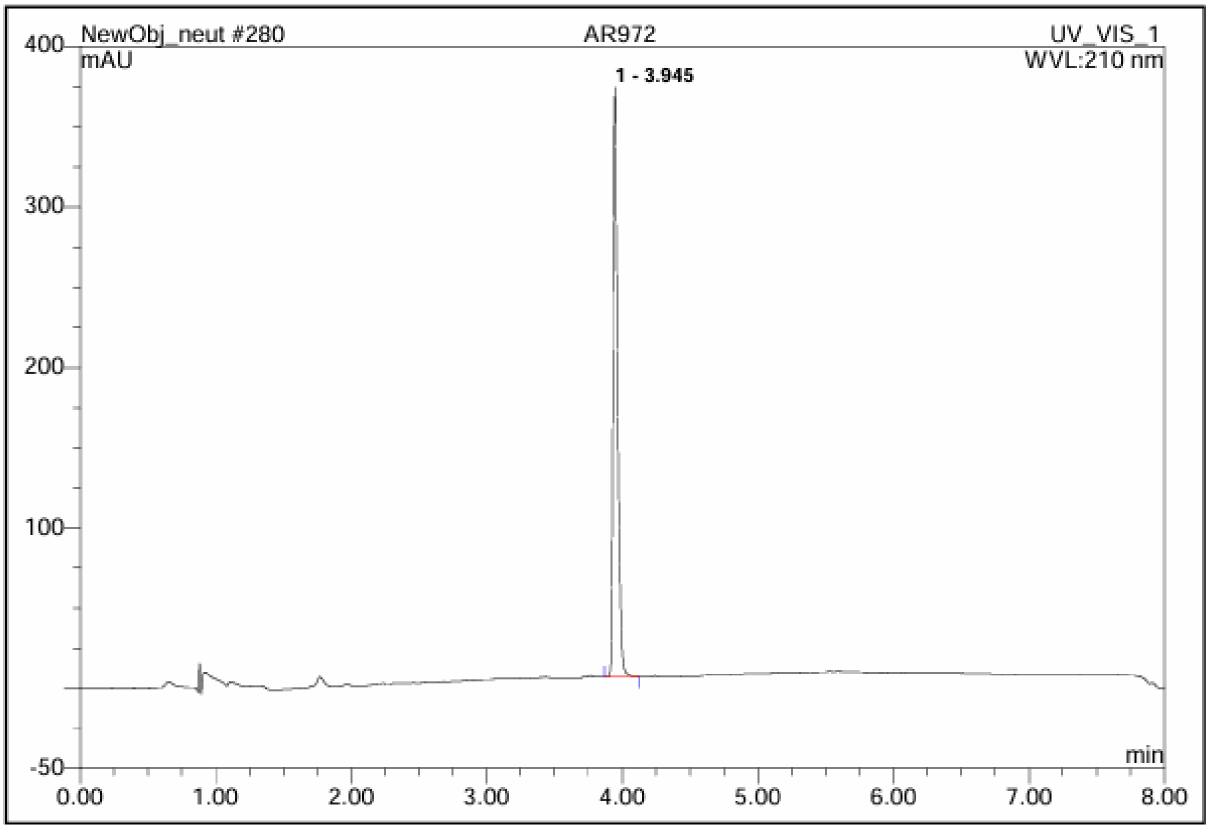

| No. | Ret.Time min | Peak Name | Height mAU | Area mAU*min | Rel.Area % | Amount | Type |
|---|---|---|---|---|---|---|---|
| 1 | 3.95 | n.a. | 367.794 | 14.087 | 100.00 | n.a. | BMB |
| **Total:** | | | 367.794 | 14.087 | 100.00 | 0.000 | |

*(3-((2-((2R,4S,5R)-4-hydroxy-5-(hydroxymethyl)tetrahydrofuran-2-yl)-1-oxo-1,2-dihydrobenzo[4,5]imidazo[1,2-c]pyrimidin-7-yl)oxy)propyl)triphenylphosphonium bromide* **6e**

**72 AR947**

| | | | |
|---|---|---|---|
| *Sample Name:* | **AR947** | *Injection Volume:* | **20.0** |
| *Vial Number:* | **RF4** | *Channel:* | **UV_VIS_1** |
| *Sample Type:* | **unknown** | *Wavelength:* | **210.0** |
| *Control Program:* | **NewObj_short** | *Bandwidth:* | **4** |
| *Quantif. Method:* | **default** | *Dilution Factor:* | **1.0000** |
| *Recording Time:* | **20.9.2025 2:03** | *Sample Weight:* | **1.0000** |
| *Run Time (min):* | **8.00** | *Sample Amount:* | **1.0000** |

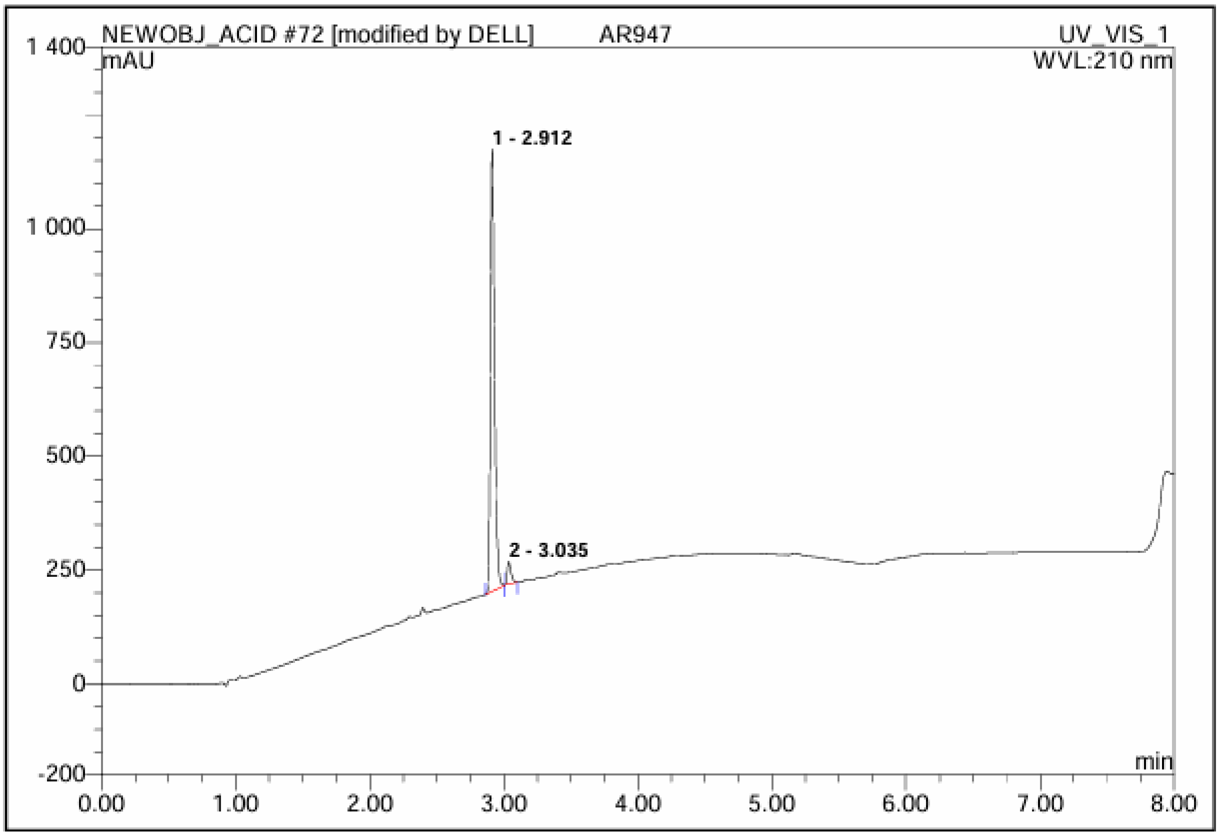

| No. | Ret.Time min | Peak Name | Height mAU | Area mAU*min | Rel.Area % | Amount | Type |
|---|---|---|---|---|---|---|---|
| 1 | 2.91 | n.a. | 972.637 | 34.824 | 95.56 | n.a. | BMB |
| 2 | 3.04 | n.a. | 48.471 | 1.618 | 4.44 | n.a. | BMB* |
| **Total:** | | | 1021.108 | 36.442 | 100.00 | 0.000 | |

*(6-((2-((2R,4S,5R)-4-hydroxy-5-(hydroxymethyl)tetrahydrofuran-2-yl)-1-oxo-1,2-dihydrobenzo[4,5]imidazo[1,2-c]pyrimidin-7-yl)oxy)hexyl)triphenylphosphonium bromide* **6f**

**84 AR944**

| | | | |
|---|---|---|---|
| *Sample Name:* | **AR944** | *Injection Volume:* | **5.0** |
| *Vial Number:* | **BC4** | *Channel:* | **UV_VIS_1** |
| *Sample Type:* | **unknown** | *Wavelength:* | **210.0** |
| *Control Program:* | **NewObj_short** | *Bandwidth:* | **4** |
| *Quantif. Method:* | **default** | *Dilution Factor:* | **1.0000** |
| *Recording Time:* | **4.10.2025 3:04** | *Sample Weight:* | **1.0000** |
| *Run Time (min):* | **8.00** | *Sample Amount:* | **1.0000** |

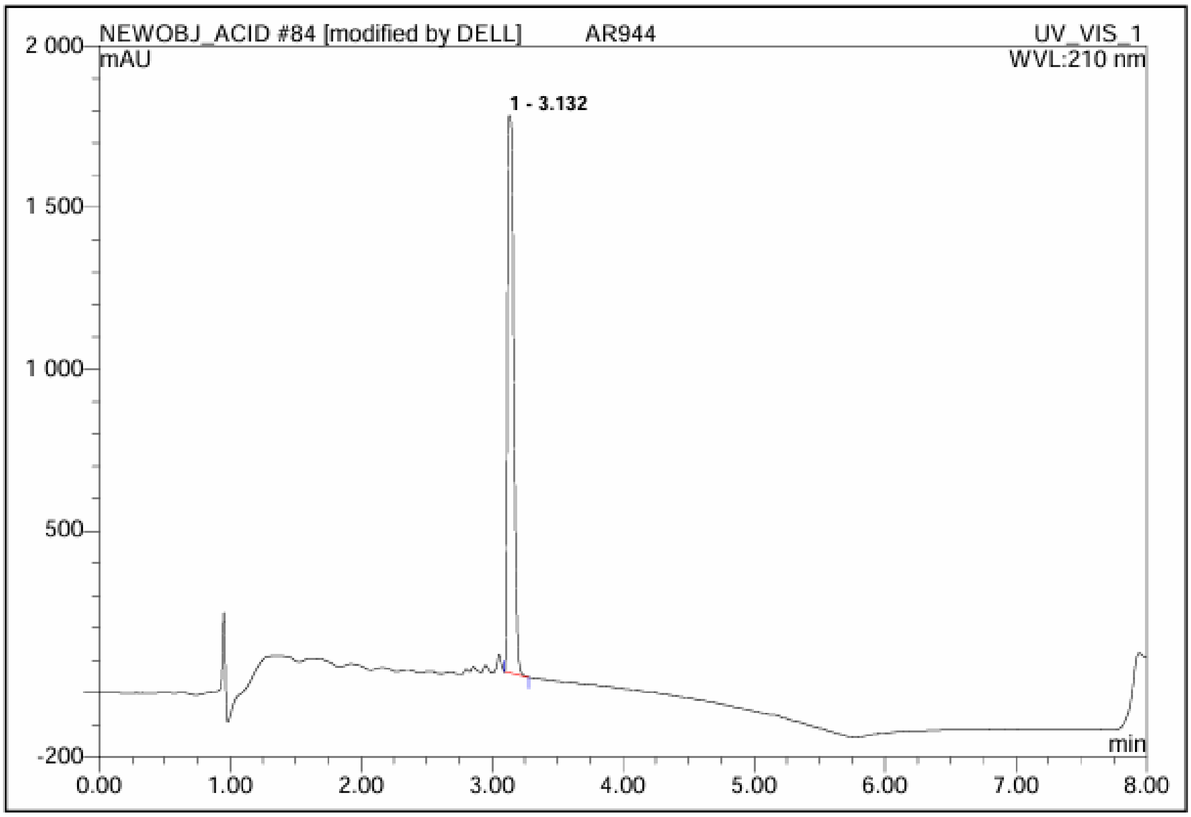

| No. | Ret.Time min | Peak Name | Height mAU | Area mAU*min | Rel.Area % | Amount | Type |
|---|---|---|---|---|---|---|---|
| 1 | 3.13 | n.a. | 1725.070 | 94.603 | 100.00 | n.a. | BMB |
| Total: | | | 1725.070 | 94.603 | 100.00 | 0.000 | |

*(10-((2-((2R,4S,5R)-4-hydroxy-5-(hydroxymethyl)tetrahydrofuran-2-yl)-1-oxo-1,2-dihydrobenzo[4,5]imidazo[1,2-c]pyrimidin-7-yl)oxy)decyl)triphenylphosphonium bromide* **6g**

**284 AR946**

| | | | |
|---|---|---|---|
| *Sample Name:* | **AR946** | *Injection Volume:* | **20.0** |
| *Vial Number:* | **RH10** | *Channel:* | **UV_VIS_1** |
| *Sample Type:* | **unknown** | *Wavelength:* | **210.0** |
| *Control Program:* | **NewObj_short** | *Bandwidth:* | **4** |
| *Quantif. Method:* | **default** | *Dilution Factor:* | **1.0000** |
| *Recording Time:* | **20.9.2025 0:46** | *Sample Weight:* | **1.0000** |
| *Run Time (min):* | **8.00** | *Sample Amount:* | **1.0000** |

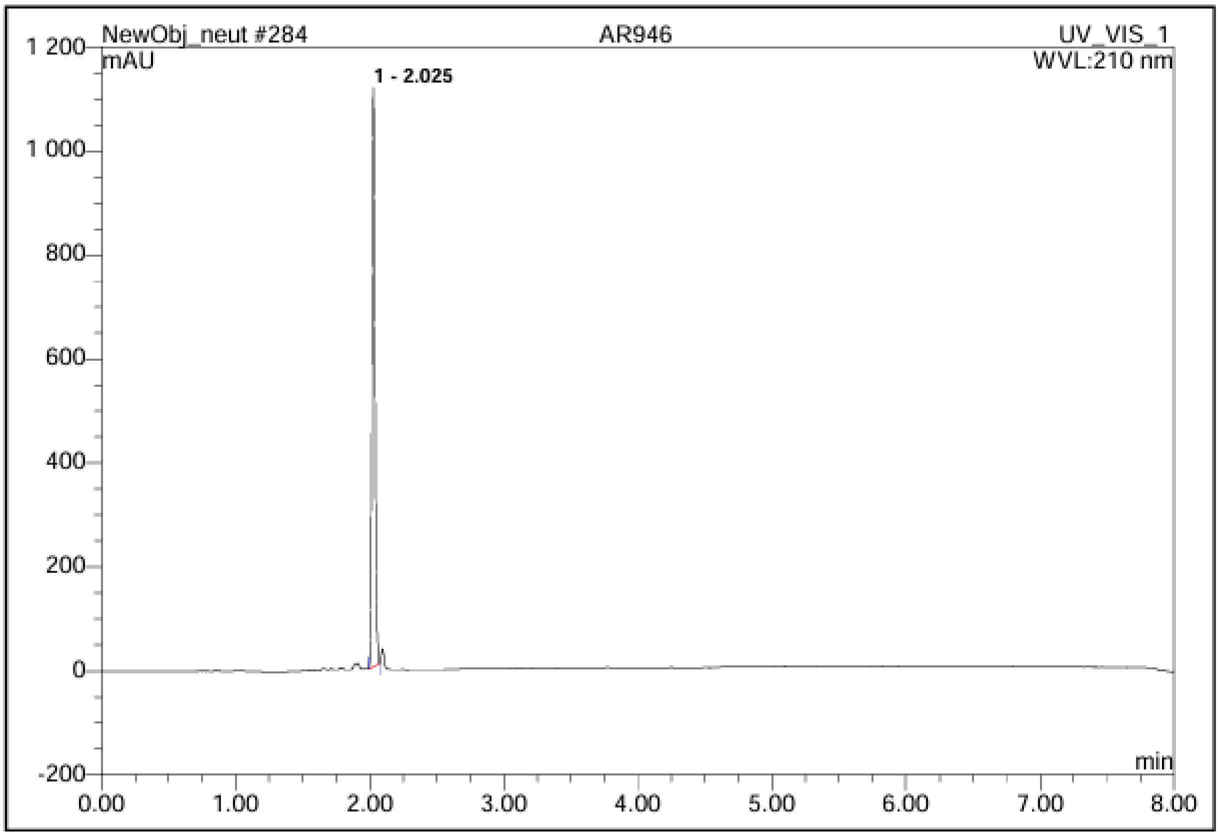

| No. | Ret.Time min | Peak Name | Height mAU | Area mAU*min | Rel.Area % | Amount | Type |
|---|---|---|---|---|---|---|---|
| 1 | 2.03 | n.a. | 1112.725 | 29.030 | 100.00 | n.a. | BMB |
| Total: | | | 1112.725 | 29.030 | 100.00 | 0.000 | |

*1-(6-((3-(4-(((2-((2R,4S,5R)-4-hydroxy-5-(hydroxymethyl)tetrahydrofuran-2-yl)-1-oxo-1,2-dihydrobenzo[4,5]imidazo[1,2-c]pyrimidin-7-yl)oxy)methyl)-1H-1,2,3-triazol-1-yl)propyl)amino)-6-oxohexyl)-3,3-dimethyl-2-((E)-3-((E)-1,3,3-trimethylindolin-2-ylidene)prop-1-en-1-yl)-3H-indol-1-ium 2,2,2-trifluoroacetate* **8a**

**82 AR941-cy3**

| | | | |
|---|---|---|---|
| *Sample Name:* | **AR941-cy3** | *Injection Volume:* | **20.0** |
| *Vial Number:* | **BF8** | *Channel:* | **UV_VIS_1** |
| *Sample Type:* | **unknown** | *Wavelength:* | **210.0** |
| *Control Program:* | **NewObj_short** | *Bandwidth:* | **4** |
| *Quantif. Method:* | **default** | *Dilution Factor:* | **1.0000** |
| *Recording Time:* | **4.10.2025 1:43** | *Sample Weight:* | **1.0000** |
| *Run Time (min):* | **8.00** | *Sample Amount:* | **1.0000** |

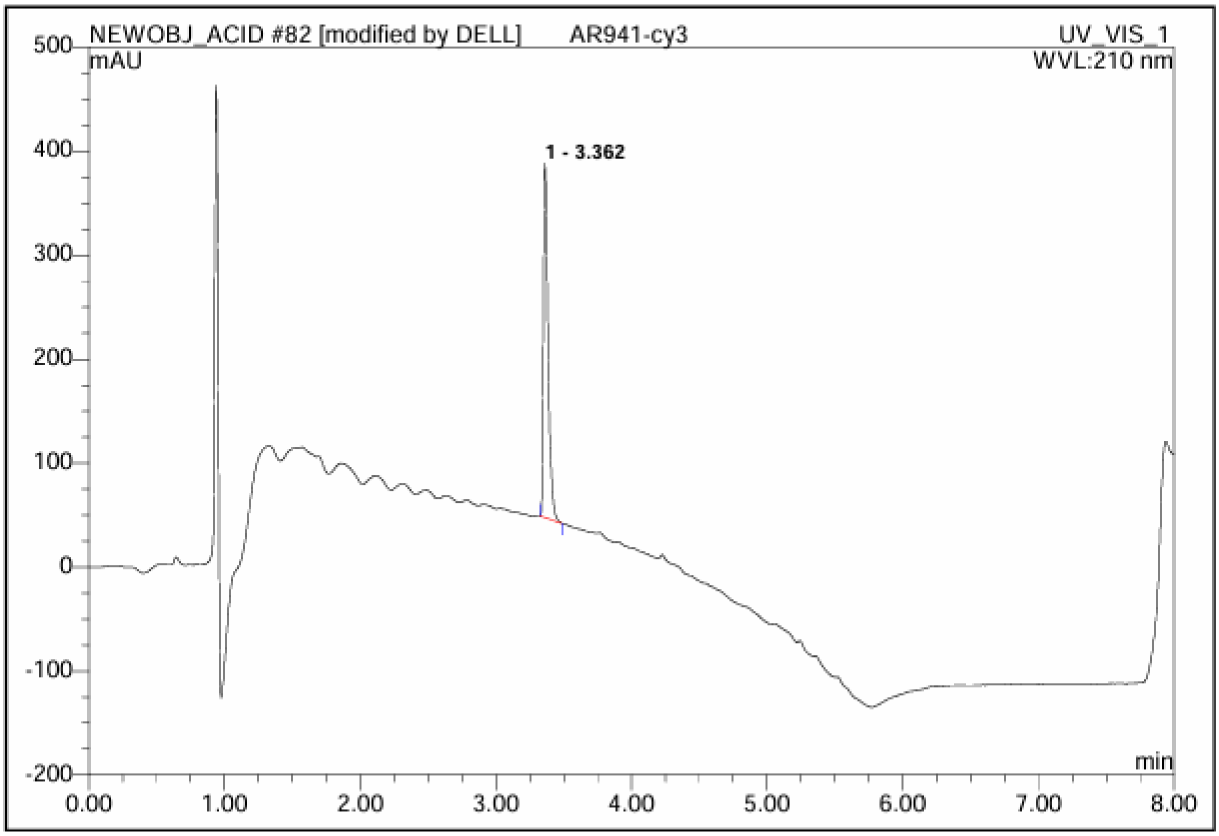

| No. | Ret.Time min | Peak Name | Height mAU | Area mAU*min | Rel.Area % | Amount | Type |
|---|---|---|---|---|---|---|---|
| 1 | 3.36 | n.a. | 340.788 | 13.554 | 100.00 | n.a. | BMB |
| **Total:** | | | 340.788 | 13.554 | 100.00 | 0.000 | |

*1-(6-((3-(4-(4-((2-((2R,4S,5R)-4-hydroxy-5-(hydroxymethyl)tetrahydrofuran-2-yl)-1-oxo-1,2-dihydrobenzo[4,5]imidazo[1,2-c]pyrimidin-7-yl)oxy)butyl)-1H-1,2,3-triazol-1-yl)propyl)amino)-6-oxohexyl)-3,3-dimethyl-2-((E)-3-((E)-1,3,3-trimethylindolin-2-ylidene)prop-1-en-1-yl)-3H-indol-1-ium 2,2,2-trifluoroacetate* **8b**

**80 AR942-cy3**

| | | | |
|---|---|---|---|
| *Sample Name:* | **AR942-cy3** | *Injection Volume:* | **20.0** |
| *Vial Number:* | **BB1** | *Channel:* | **UV_VIS_1** |
| *Sample Type:* | **unknown** | *Wavelength:* | **210.0** |
| *Control Program:* | **NewObj_short** | *Bandwidth:* | **4** |
| *Quantif. Method:* | **default** | *Dilution Factor:* | **1.0000** |
| *Recording Time:* | **4.10.2025 0:02** | *Sample Weight:* | **1.0000** |
| *Run Time (min):* | **8.00** | *Sample Amount:* | **1.0000** |

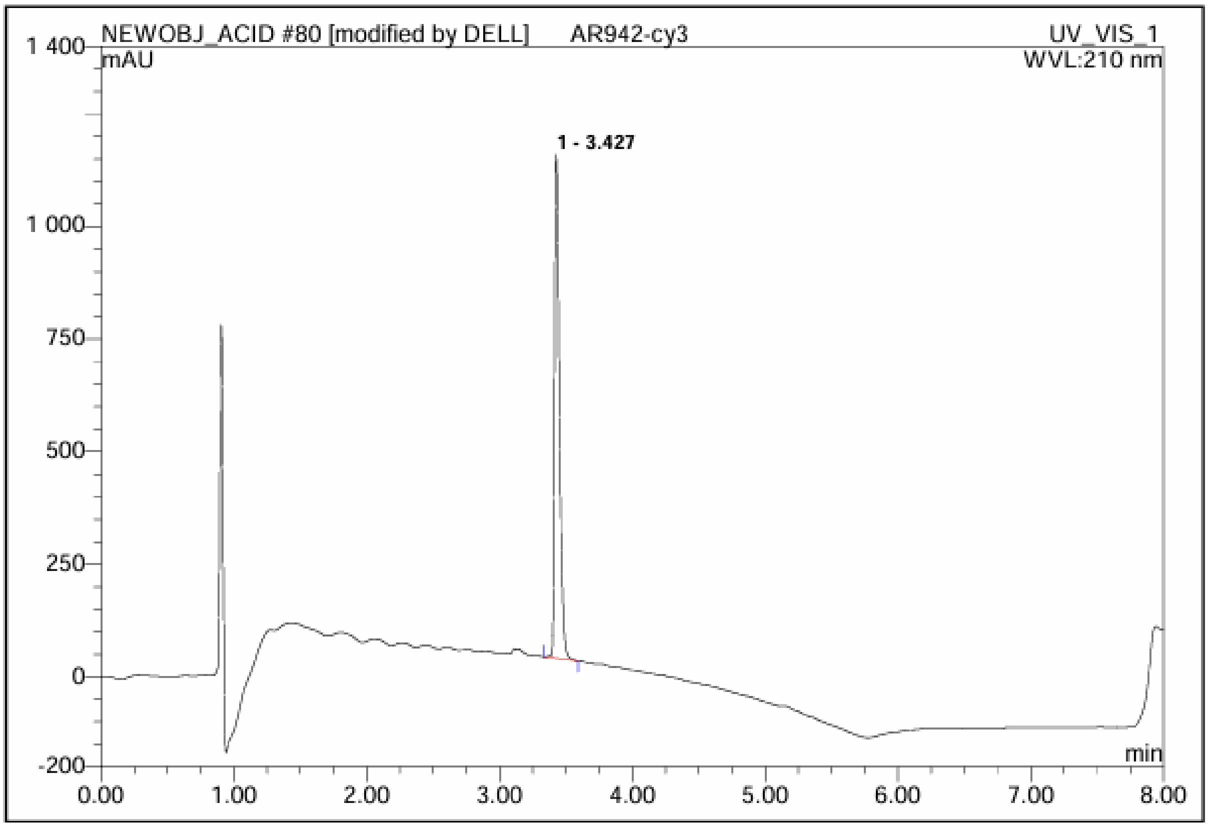

| No. | Ret.Time min | Peak Name | Height mAU | Area mAU*min | Rel.Area % | Amount | Type |
|---|---|---|---|---|---|---|---|
| 1 | 3.43 | n.a. | 1118.203 | 46.874 | 100.00 | n.a. | BMB |
| Total: | | | 1118.203 | 46.874 | 100.00 | 0.000 | |

*1-(6-((3-(4-(((2-((2R,4S,5R)-4-hydroxy-5-(hydroxymethyl)tetrahydrofuran-2-yl)-1-oxo-1,2-dihydrobenzo[4,5]imidazo[1,2-c]pyrimidin-7-yl)oxy)methyl)-1H-1,2,3-triazol-1-yl)propyl)amino)-6-oxohexyl)-3,3-dimethyl-2-((1E,3E)-5-((E)-1,3,3-trimethylindolin-2-ylidene)penta-1,3-dien-1-yl)-3H-indol-1-ium 2,2,2-trifluoroacetate* **8c**

**83 AR941-cy5**

| | | | |
|---|---|---|---|
| *Sample Name:* | **AR941-cy5** | *Injection Volume:* | **20.0** |
| *Vial Number:* | **BH11** | *Channel:* | **UV_VIS_1** |
| *Sample Type:* | **unknown** | *Wavelength:* | **210.0** |
| *Control Program:* | **NewObj_short** | *Bandwidth:* | **4** |
| *Quantif. Method:* | **default** | *Dilution Factor:* | **1.0000** |
| *Recording Time:* | **4.10.2025 2:43** | *Sample Weight:* | **1.0000** |
| *Run Time (min):* | **8.00** | *Sample Amount:* | **1.0000** |

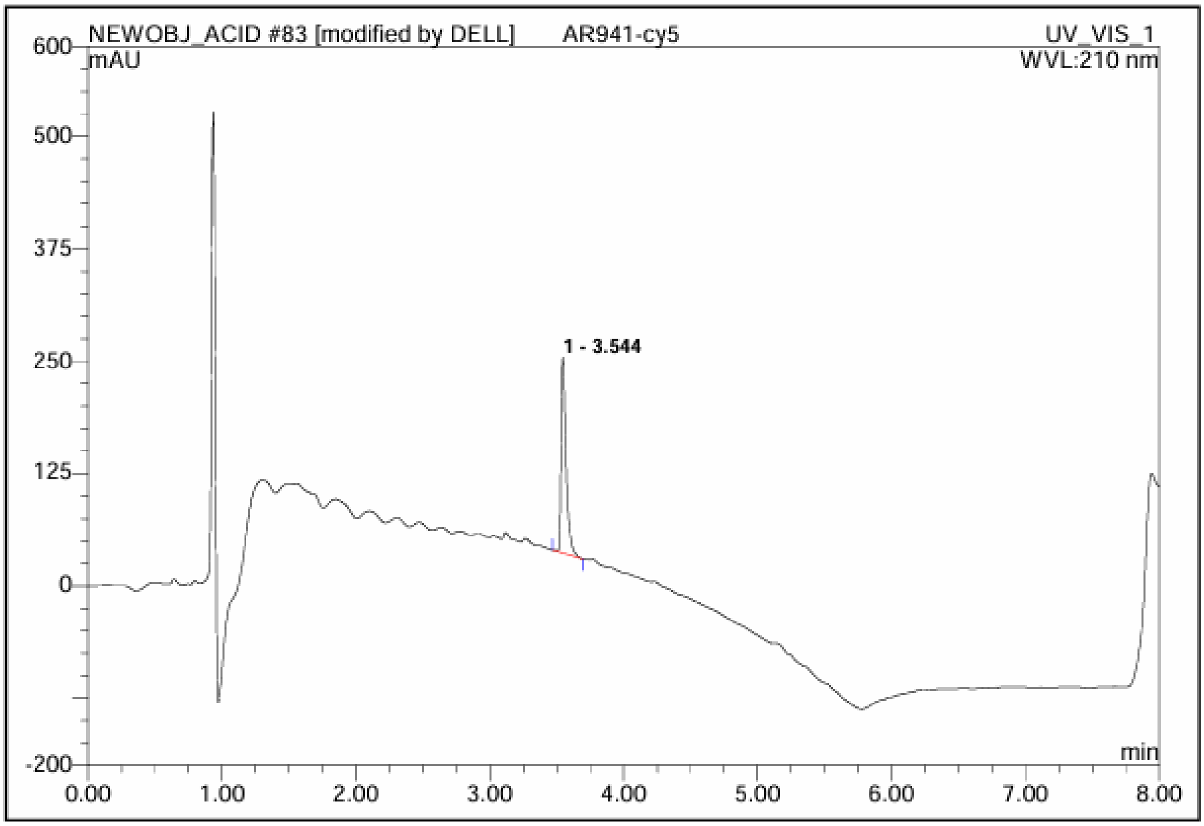

| No. | Ret.Time min | Peak Name | Height mAU | Area mAU*min | Rel.Area % | Amount | Type |
|---|---|---|---|---|---|---|---|
| 1 | 3.54 | n.a. | 218.159 | 9.192 | 100.00 | n.a. | BMB |
| **Total:** | | | 218.159 | 9.192 | 100.00 | 0.000 | |

*1-(6-((3-(4-(4-((2-((2R,4S,5R)-4-hydroxy-5-(hydroxymethyl)tetrahydrofuran-2-yl)-1-oxo-1,2-dihydrobenzo[4,5]imidazo[1,2-c]pyrimidin-7-yl)oxy)butyl)-1H-1,2,3-triazol-1-yl)propyl)amino)-6-oxohexyl)-3,3-dimethyl-2-((1E,3E)-5-((E)-1,3,3-trimethylindolin-2-ylidene)penta-1,3-dien-1-yl)-3H-indol-1-ium 2,2,2-trifluoroacetate* **8d**

**81 AR942-cy5**

| | | | |
|---|---|---|---|
| *Sample Name:* | **AR942-cy5** | *Injection Volume:* | **20.0** |
| *Vial Number:* | **BD7** | *Channel:* | **UV_VIS_1** |
| *Sample Type:* | **unknown** | *Wavelength:* | **210.0** |
| *Control Program:* | **NewObj_short** | *Bandwidth:* | **4** |
| *Quantif. Method:* | **default** | *Dilution Factor:* | **1.0000** |
| *Recording Time:* | **4.10.2025 0:14** | *Sample Weight:* | **1.0000** |
| *Run Time (min):* | **8.00** | *Sample Amount:* | **1.0000** |

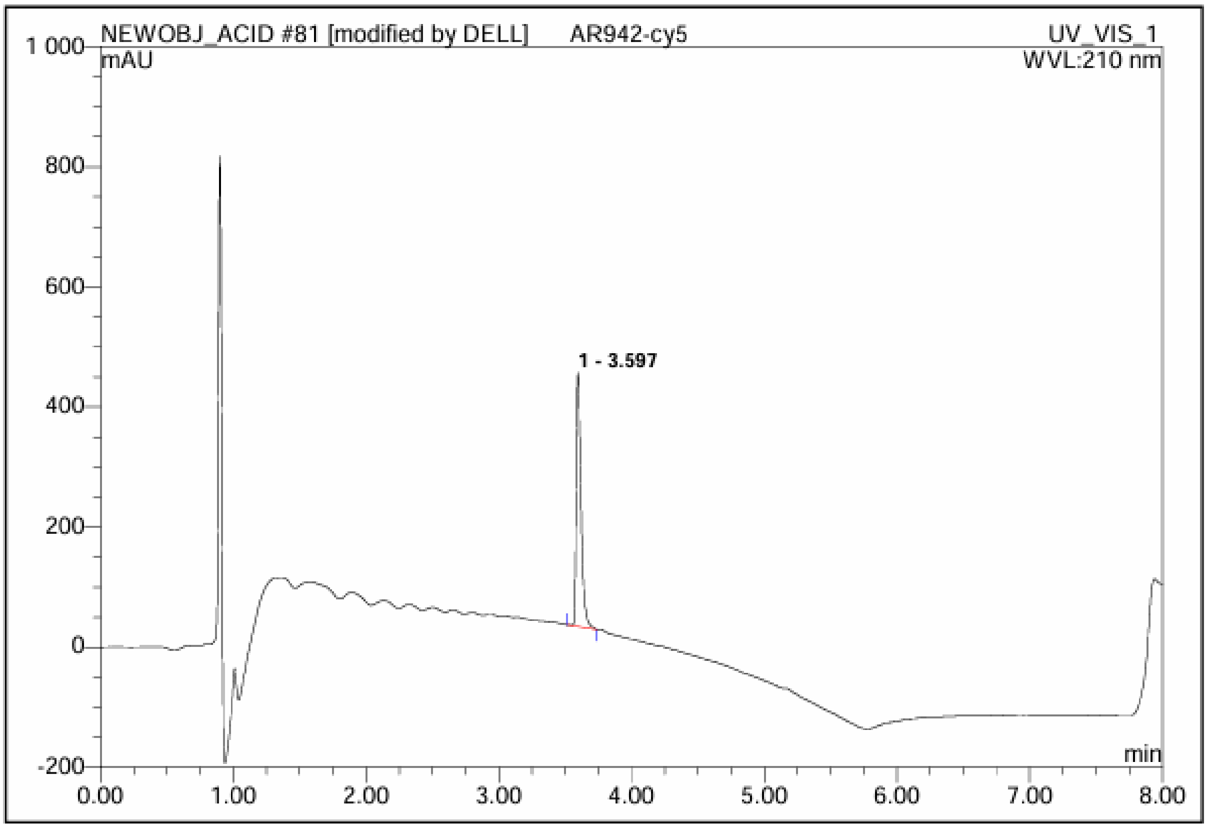

| No. | Ret.Time min | Peak Name | Height mAU | Area mAU*min | Rel.Area % | Amount | Type |
|---|---|---|---|---|---|---|---|
| 1 | 3.60 | n.a. | 423.064 | 17.553 | 100.00 | n.a. | BMB |
| Total: | | | 423.064 | 17.553 | 100.00 | 0.000 | |
